# Identifying Interpretable Latent Factors with Sparse Component Analysis

**DOI:** 10.1101/2024.02.05.578988

**Authors:** Andrew J. Zimnik, K. Cora Ames, Xinyue An, Laura Driscoll, Antonio H. Lara, Abigail A. Russo, Vladislav Susoy, John P. Cunningham, Liam Paninski, Mark M. Churchland, Joshua I. Glaser

**Affiliations:** Department of Neuroscience, Columbia University Medical Center, New York, NY, USA; Zuckerman Institute, Columbia University, New York, NY, USA; Grossman Center for the Statistics of Mind, Columbia University, New York, NY, USA; Center for Theoretical Neuroscience, Columbia University, New York, NY, USA; Department of Neurology, Northwestern University, Chicago, IL, USA; Interdepartmental Neuroscience Program, Northwestern University, Chicago, IL, USA; Department of Electrical Engineering, Stanford University, Stanford, CA, USA; Allen Institute for Neural Dynamics, Allen Institute, Seattle, CA, USA; Department of Physics, Harvard University, Cambridge, MA, USA; Center for Brain Science, Harvard University, Cambridge, MA, USA; Department of Statistics, Columbia University, New York, NY, USA; Kavli Institute for Brain Science, Columbia University Medical Center, New York, NY, USA; Department of Computer Science, Northwestern University, Evanston, IL, USA

## Abstract

In many neural populations, the computationally relevant signals are posited to be a set of ‘latent factors’ – signals shared across many individual neurons. Understanding the relationship between neural activity and behavior requires the identification of factors that reflect distinct computational roles. Methods for identifying such factors typically require supervision, which can be suboptimal if one is unsure how (or whether) factors can be grouped into distinct, meaningful sets. Here, we introduce Sparse Component Analysis (SCA), an unsupervised method that identifies interpretable latent factors. SCA seeks factors that are sparse in time and occupy orthogonal dimensions. With these simple constraints, SCA facilitates surprisingly clear parcellations of neural activity across a range of behaviors. We applied SCA to motor cortex activity from reaching and cycling monkeys, single-trial imaging data from *C. elegans*, and activity from a multitask artificial network. SCA consistently identified sets of factors that were useful in describing network computations.

## Introduction

The study of computation by neural populations has experienced a conceptual shift. Traditionally, one first characterized single-neuron response properties, then extrapolated to determine how the population would behave. This approach worked well in situations where population-level function (e.g. saccade generation) was a straightforward extension of single-neuron responses (e.g. tuning for target amplitude and direction in the superior colliculus)^1,2^. Yet with time, it became appreciated that this strategy often needs to be inverted. In many areas, single-neuron responses are complex, heterogeneous, and make little sense except in the context of population-level computation. This realization led to an increased desire for tools for investigating computation at the population level.

Population-level computations, in many biological and artificial networks, involve ‘latent factors’^3–10^. Latent factors are simply signals that are shared across the population. In many networks, the factors are the computationally relevant signals, as they are consistent across trials, while the activity of individual neurons is noisy and variable^11^. The evolution and interactions of factors can yield explanations of how a particular computation is accomplished (e.g.,^11–20^). However, factors cannot be observed directly; instead, they must be estimated from population activity using dimensionality reduction. This need for inference poses a hurdle. While there are typically many statistically valid estimates of factors, not all provide equally interpretable views of the underlying computational mechanisms.

As a concrete example of the importance and utility of extracting interpretable factors, consider motor cortical activity during delayed reaching (Fig. 1A). Individual-neuron responses are complex and difficult to interpret^21^. Most neurons are active during both the delay period and during movement, with no clear similarity between these two activity patterns^22–25^. Yet at the population level, one can identify two distinct sets of factors. ‘Preparatory factors’ (Fig. 1A, *left*) are active solely during the delay, while ‘execution-related factors’ (Fig. 1A, *right*) are active solely during the reach^26,27^. This separation maps onto basic hypotheses regarding network computation, where execution dynamics generate activity patterns that form descending commands^15^, while preparatory factors seed an initial state from which upcoming execution-related factors evolve^22,28,29^. This framework, and the ability to empirically separate preparatory and execution factors, has made it possible to address a variety of outstanding questions regarding how movements are prepared^27^, executed^15^, and arranged in sequences^30^. This progress was possible only because it was suspected, in advance, that preparation and execution might be distinct processes subserved by distinct sets of factors. Careful task design also allowed researchers to know when, within each trial, these two processes occurred and where the likely boundary was between them.

**Figure 1.**
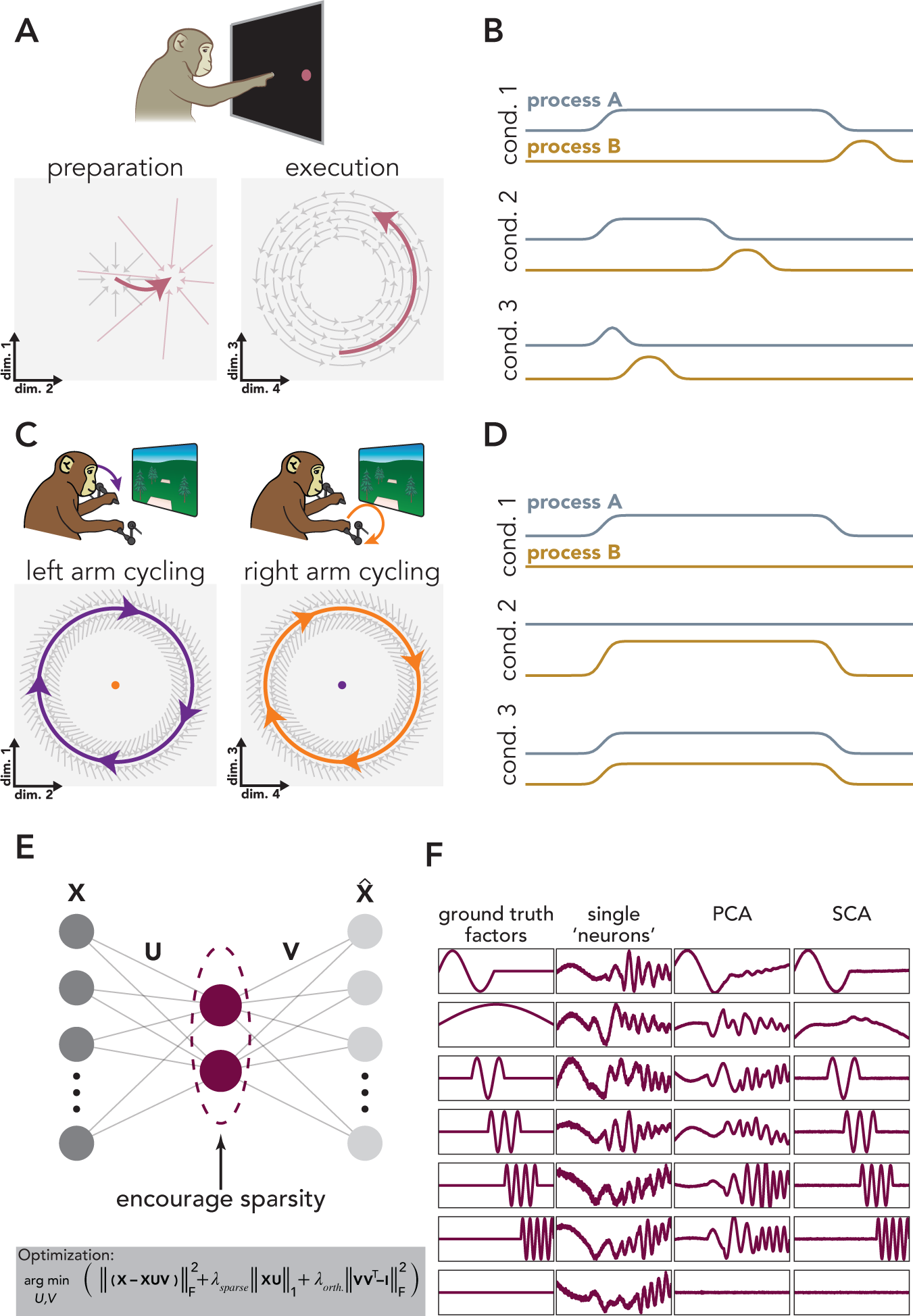
SCA seeks non-synchronous factors. **A.** Non-synchronous factors during reaching. Left: preparation: Condition-specific inputs (*small red arrows*) cause preparatory factors (*thick red arrow*) to settle at a single fixed point. Absent this input, activity would decay to a baseline determined by the intrinsic dynamics (*gray arrows*). Right: execution dynamics shared across conditions (*gray arrows*) and the initial state of the system (*beginning of red arrow*) determine the trajectory of the execution-related factors (*red arrow*). **B.** Time course of two non-synchronous neural processes that are not independent (e.g., **A**). The two processes exhibit a degree of temporal flexibility because the duration of Process A (*silver trace*) can differ across conditions. However, Process B (*gold trace*) always follows the end of process A. **C.** Non-synchronous factors during cycling. Motor cortical dynamics (*gray arrows*) maintain stable limit cycles that generate left-arm moving (*purple*) and right-arm moving (*orange*) factors. These two processes occur in orthogonal subspaces; the space that captures a large amount of left-arm variance (left) only accounts for a small fraction of the neural variance when the right-arm is moving (*orange dot*, left space), and *vice versa*. **D.** Time course of two non-synchronous processes that are independent (e.g., **C**). **E.** SCA architecture and cost function. **F.** SCA recovers ground-truth latent factors in idealized data. We constructed a population of 50 ‘neurons’ via orthonormal mappings from the ground-truth factors, plus Gaussian noise.

Reach preparation and execution provide a specific example of a general need: parsing population activity into dimensions that respect basic divisions between the computational roles of different factors, if such divisions exist. It has become common to use unsupervised dimensionality reduction methods, such as principal component analysis (PCA) as a first step in estimating factors. Yet a well-known hurdle is that unsupervised methods frequently fail to provide the desired parcellation; PCA-estimated factors are often as ‘mixed’ as the individual neurons one started with. The field has thus developed a variety of approaches that can be more appropriate in specific situations. A simple form of supervision is to manually label times, dividing data into groups based on prior knowledge or assumptions, before applying an otherwise-unsupervised method (e.g.,^17,19^). Alternatively, explicitly supervised methods such as demixed PCA (dPCA) use labeling based both on time within a trial and on the condition label of that trial^31^. Related methods such as targeted dimensionality reduction (TDR) and its successor, model-based TDR,^32^ use regression to identify factors that covary with task-related variables^12^. Finally, methods such as jPCA^15^ avoid labels, but seek hypothesized structure that is typically dataset- or experiment-specific (e.g., factors that obey linear dynamics). All these approaches are limiting in the many situations where one may not know the correct labels, temporal divisions, or potential forms of population-level structure.

Here, we present an unsupervised method for identifying interpretable latent factors. Sparse Component Analysis (SCA) seeks factors that are sparse in time and evolve within orthogonal dimensions. In seeking this simple structure, SCA is able to recover factors that only reflect a single computational role. SCA does so by leveraging the fact that many neural computations involve distinct processes, each associated with different sets of latent factors, which are not always active at the same time. SCA is a variant of dictionary learning^33,34^, with a simple cost function that reflects very general assumptions about neural computations and therefore can be usefully applied to a wide range of datasets.

We applied SCA to data collected from three experimental models (rhesus monkeys, *C. elegans*, and artificial networks) during a variety of behavioral tasks. SCA identified factors that have been previously described using supervised methods, verifying that the method can recover expected features without the need for supervision. SCA also identified previously undescribed factors, validating the method as a tool for scientific discovery. In monkey motor cortical activity collected during a reaching task, SCA identified separate factors related to reach preparation, execution, and postural maintenance. In activity recorded during unimanual cycling, a potentially more complicated behavior that involves more extended movements, SCA produced surprisingly analogous factors to those observed during reaching – factors related to movement preparation, movement execution, and postural maintenance. When applied to bimanual cycling activity, SCA identified factors that were selective for one of the two arms. Surprisingly, many of these factors were active during both unimanual movements and bimanual movements, demonstrating that SCA will not ‘invent’ sparsity that is not present in data. Next, we applied SCA to neural data collected from *C. elegans* during mating. SCA identified factors, and individual neurons, related to specific mating motifs. Finally, in a multi-task RNN, SCA recovered factors that reflected the compositional nature of the network’s computations^13,35^.

## Results

### An unsupervised method for identifying interpretable factors

When attempting to understand a given neural computation, it is often useful to partition neural activity into signals that reflect distinct computational roles. Typically, this partitioning is accomplished via careful task design. By controlling the timing of key task events, researchers determine when different neural processes occur. Once distinct processes are confined to specific task epochs, unsupervised dimensionality reduction methods can be used to identify the latent factors active during that epoch. For example, in a delayed reaching task, researchers control the duration of the delay period and therefore determine when preparatory and execution-related processes occur^23,24^. These two processes not only occur at different times, but also within different sets of dimensions; the neural dimensions occupied during preparation are nearly orthogonal to those occupied during execution^26^. Preparatory and execution-related factors can therefore be identified by training PCA on delay-period or execution-period activity (Fig. 1A).

Preparatory and execution-related factors exhibit a property that may be common to sets of factors that perform different computational roles: they are ‘non-synchronous’, meaning that these two sets of factors can be active at different times. In other words, preparation and execution exhibit temporal flexibility relative to one another. Preparation can be lengthy or brief^27,36,37^, preparation for a second movement can occur simultaneously with execution of a first^30,36^, or can occur after an initial movement ends^30^. This flexibility is incomplete – preparation must presumably always precede execution – yet it is still the case that the envelopes of preparation and execution are quite different in different situations (Fig. 1B). A second example of non-synchronous factors is observed during bimanual coordination (Fig. 1C,D). Distinct sets of factors are active when the left or the right arm moves^38,39^ and are co-active when both arms move simultaneously ^40^. As was true for preparatory and execution factors, left-arm and right-arm factors evolve in orthogonal sets of dimensions^38,39^. Further examples of non-synchronous factors that occupy orthogonal dimensions have been observed across brain regions and species, during a wide variety of tasks (e.g.,^16,17,19,31,41–43)^. In the past, discerning the presence of non-synchronous factors required either clever task design (e.g., the classic delayed-reach paradigm) or supervised methods to separate one factor from another (e.g., dPCA, TDR). A natural goal is thus a method that identifies non-synchronous factors in an unsupervised manner.

We designed SCA (Fig. 1E) to accomplish this goal. SCA was designed to identify factors that are active at different times and/or overlap in time but display the temporal flexibility described above. SCA finds:

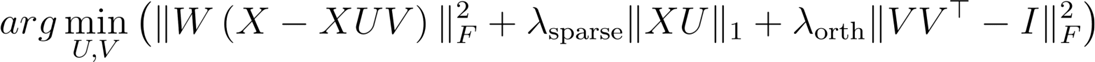

The first term ensures that the learned latent factors capture a large fraction of the neural variance, the second term encourages the factors to be sparse in time (either within a trial or across conditions), and the final term encourages the loadings to be orthogonal.

Let **X***^T×N^* be a matrix of neural data, where *N* and *T* indicate the number of recorded neurons and number of time points, respectively. If there are multiple experimental conditions, neural data within **X** can be concatenated across conditions, so *T* represents the total number of time points across conditions. In order to find the low-dimensional factors, we learn the factors via a mapping, **U***^N×K^*, from the neural activity space (Fig. 1E), where *K* indicates the number of requested factors. **V***^K×N^* is a matrix that reconstructs the high-dimensional neural activity from the factors. We also include offset terms in both these mappings (omitted for brevity in the equation below, see *Methods* for full equation). We weight the samples in **X**, via **W***^T×T^*, such that the reconstruction of timepoints of high firing-rate neural activity are not given inherent precedence over low firing rate activity (e.g., the reconstruction of preparatory activity does not suffer from the attempt to reconstruct the relatively higher firing rates associated with execution-related activity, see details in *Methods*).

It may seem odd to encourage sparsity in the factors, as the factors of interest are often non-sparse, and may be active during many, or even most, task epochs. However, encouraging sparsity means that the cost–function is minimized when non-synchronous factors are isolated in different dimensions. For example, a solution in which factor 1 is active only during preparation and factor 2 is active only during movement is preferred to one in which both dimensions are active at both times. Thus, SCA aims to yield factors that can be naturally grouped in ways that aid scientific interpretation.

Importantly, this cost function is agnostic regarding the experimental ‘meaning’ of different times. For example, SCA has no knowledge of whether activity corresponds to an imposed preparatory period versus an execution epoch or whether activity occurs during one experimental condition versus another. The same cost function can therefore be used across a great variety of experimental situations, as we illustrate below.

To begin, we verified that SCA recovers the expected factors from a population of artificially-generated ‘neurons’ (Fig. 1F). By design, the activity of these individual neurons reflects the influence of multiple non-synchronous factors (Fig. 1F, second column). PCA, as expected, does not resolve this mixing (Fig. 1F, third column); individual PCA factors and neurons are both combinations of the factors. SCA, however, recovers the ground truth factors (Fig. 1F, fourth column). While it is useful to validate that SCA can recover non-synchronous factors in idealized data, in empirical neural data, we do not have access to the ‘ground truth’ factors. In such cases, why are SCA factors preferable over PCA factors if both are statistically valid views of the data? SCA factors are preferable if they invite testable interpretations about how a given computation is performed. We demonstrate below, using a variety of datasets, that SCA often provides such factors.

### Center-out reaching

Motor cortical activity during center-out reaching has been studied (and modeled) extensively^14,15,27,44,45^. This task is thus a useful testbed for SCA because the field has developed a reasonably mature understanding of the neural mechanisms that underlie this behavior. Reaching involves three distinct processes, each reflected by a set of latent factors. Preparatory factors become active shortly after the onset of the target and remain relatively static throughout the delay period^27,46^. Shortly after the delivery of the go-cue (and approximately 150 ms before the hand begins to move), activity in preparatory dimensions collapses while execution-related factors simultaneously become active^26,27^. Finally, factors related to postural-maintenance become active while the animal holds its hand at the peripheral target^47,48^. To determine whether SCA could identify preparatory, execution-related, and posture-related factors, without the supervision such parcellation previously required, we applied SCA to trial-averaged activity from motor cortex, recorded while monkeys performed a delayed center-out reaching task (Fig. 2A,B).

**Figure 2.**
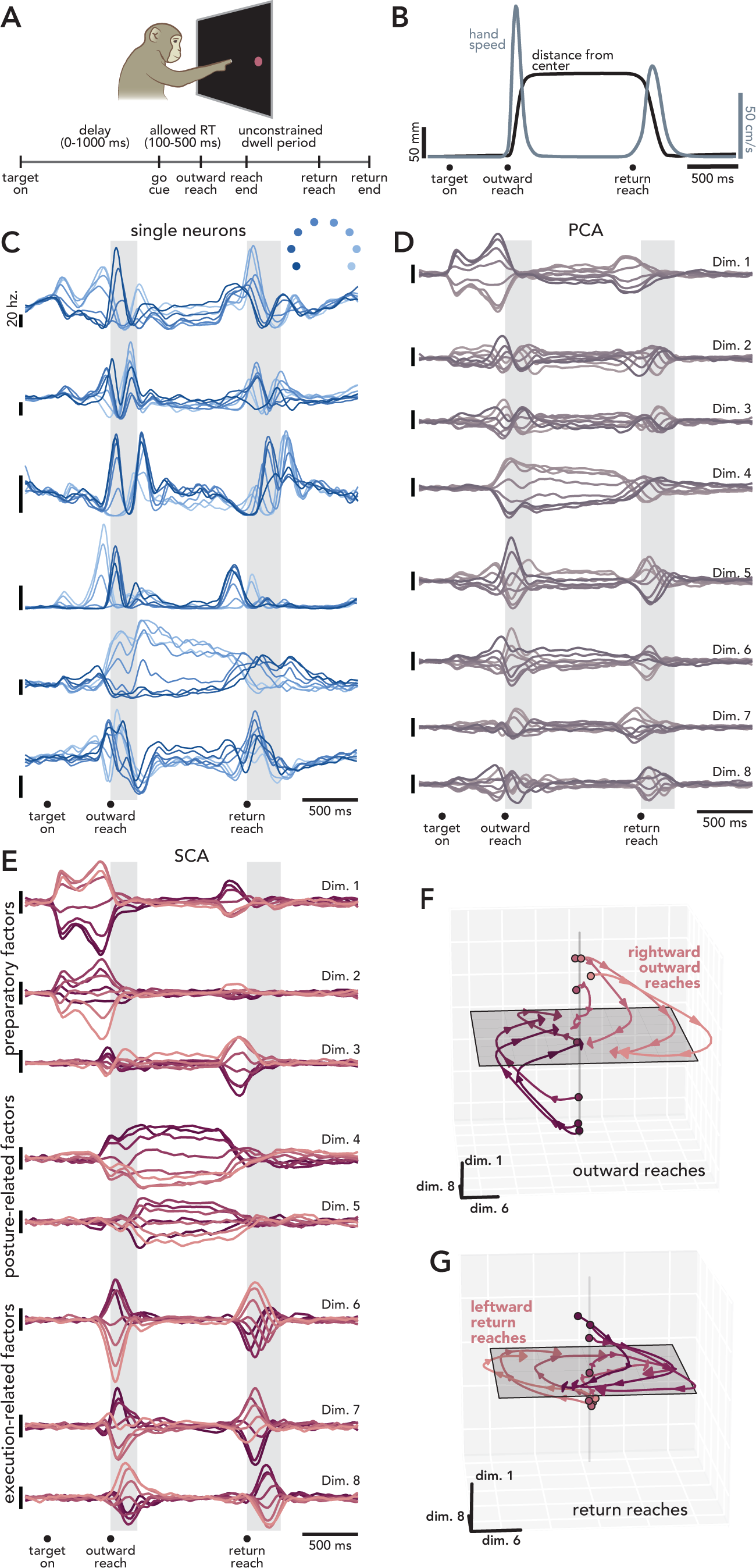
SCA identifies preparatory, execution-related, and posture-related factors in motor cortical reaching data. **A.** Trial timing. **B.** Example behavior for one condition (averaged across trials). **C.** Six example motor cortical neurons. Each trace corresponds to a reach direction. *Gray rectangles* indicate the median duration of all reaches. **D.** Top eight PCA factors. The factors have been ordered by time of maximum occupancy. **E.** Projections in eight SCA dimensions. As in **D**, factors have been ordered by time of maximum occupancy. **F.** Outward reach activity projected into three SCA dimensions. **G.** Projection of return reach activity in the same space as **F**.

In this task, monkeys held their hand at a central touchpoint and were shown one of eight peripheral targets. Following an unpredictable go cue, the animals quickly captured the instructed target and maintained this position until a juice reward was delivered. After receiving a reward, the monkey returned to the starting position to begin the next trial. This return reach was self-initiated; the task did not require the monkey to return to the touchpoint within a certain amount of time.

The fact that reaching is composed of three distinct processes was not apparent either at the level of single units (Fig. 2D) nor factors identified via PCA (Fig. 2D). In both, neural activity relating to preparatory, execution-related, and posture-related processes are mixed. For example, the neuron shown in the top row of Fig. 2C was active during the delay period, the outward reach, the hold period (when the monkey was holding the captured target), and the return reach. Many of the PCA factors were also continuously active throughout the trial (e.g., Fig. 2D, *factors 2 and 3*). While individual neurons and PCA factors hint at the existence of discrete processes – for example, the ‘tuning’ of individual neurons often changed between the delay period and the execution epoch (e.g., Fig. 2C, *top row*) and some individual PCA factors were more active during a single task epoch (e.g., Fig. 2D, *top row*) – these hints are somewhat subtle. Neither view of the data readily offers an interpretable view of the underlying computation.

SCA, conversely, produced a remarkably clear view of the divisions between neural processes (Fig. 2E). Without supervision, SCA found factors that were primarily active during only one task epoch: preparation, reach execution, or postural-maintenance. Consider activity before and during the outward reach (Fig. 2E). Shortly after target onset, activity emerges in dimensions 1 and 2 and is maintained throughout the delay period, a pattern expected for preparatory activity during this task^22,27,46^. Shortly before reach onset, activity in these first two dimensions collapses, while activity in dimensions 4, 6, 7, and 8 grows. Dimensions 6, 7, and 8 are occupied briefly – for approximately the same duration as the outward reach – while activity in dimensions 4 and 5 remain high. These patterns are consistent with dimensions 1 and 2 being preparatory dimensions, dimensions 6, 7, and 8 being execution-related dimensions, and dimensions 4 and 5 being posture-related dimensions.

The factors’ patterns of activity around the return reach agree with this interpretation. The putative posture-related factors (dimensions 4 and 5) remain active while the monkey maintains his hand position at the peripheral target. Approximately 250 ms before the self-initiated return reach, activity in these posture dimensions begins to recede. At the same time, activity reemerges in dimension 1, and grows more dramatically in dimension 3. That dimension 3 is only occupied prior to the return reach is not unexpected given kinematic differences between the outward and return reaches, a point we return to below. Before the onset of the return reach, activity in dimensions 1 and 3 decreases, while dimensions 6, 7, and 8 (the putative execution-related dimensions) become transiently occupied, as they were during the outward reach. Similar results were found using data from a second monkey (Figure S1); each SCA factor was primarily active during the preparatory, execution, or posture epoch.

In order to visualize the relationship between factors, it is often useful to plot activity in a state space (Fig. 2F,G). Here, we’ve plotted activity in a three dimensional space defined by one preparatory dimension (SCA dimension 1) and two execution-related dimensions (SCA dimensions 6 and 8). This view highlights both the relationship between preparatory activity and execution activity as well as the relationship between execution-related factors. During outward reaches (Fig. 2F), preparatory activity sets the initial state for execution-related activity^22^. Just before movement, preparatory activity diminishes, while activity emerges in execution-related dimensions (Fig. 2F, *gray plane*). Activity within this plane appears to be dictated by a single set of dynamics, which are shared across conditions^15^. During return reaches (Fig. 2G), the same process unfolds in the same three dimensions (although the preparatory activity is weaker in this dimension, see below). The ordering of conditions is largely reversed between outward and return reaches. This reversal reflects task geometry; conditions that begin with an outwards reach to the right are followed by a return reach to the left.

### Quantitative Comparisons and Characterization

Qualitatively, SCA appears to produce more interpretable factors than PCA. Specifically, SCA factors seem to correspond to distinct computational roles, while PCA factors reflect a mixture of computational processes. To validate this impression, we performed a number of quantitative comparisons between SCA and PCA factors. We compared SCA factors to factors recovered by weighted PCA (wPCA, see *Methods*), a PCA variant that is more comparable to SCA in that it also uses sample-weighting. The only substantial difference between these two methods is that wPCA does not attempt to identify sparse factors.

First, we found that the improved interpretability of SCA factors does not come at the cost of decreased reconstruction accuracy. Although SCA factors are only active during a single epoch, SCA and wPCA factors account for virtually identical amounts of total neural variance (Fig. 3A). Additionally, SCA and wPCA perform equally well when asked to generalize to held-out neurons and conditions (Fig. 3A, *inset*).

**Figure 3.**
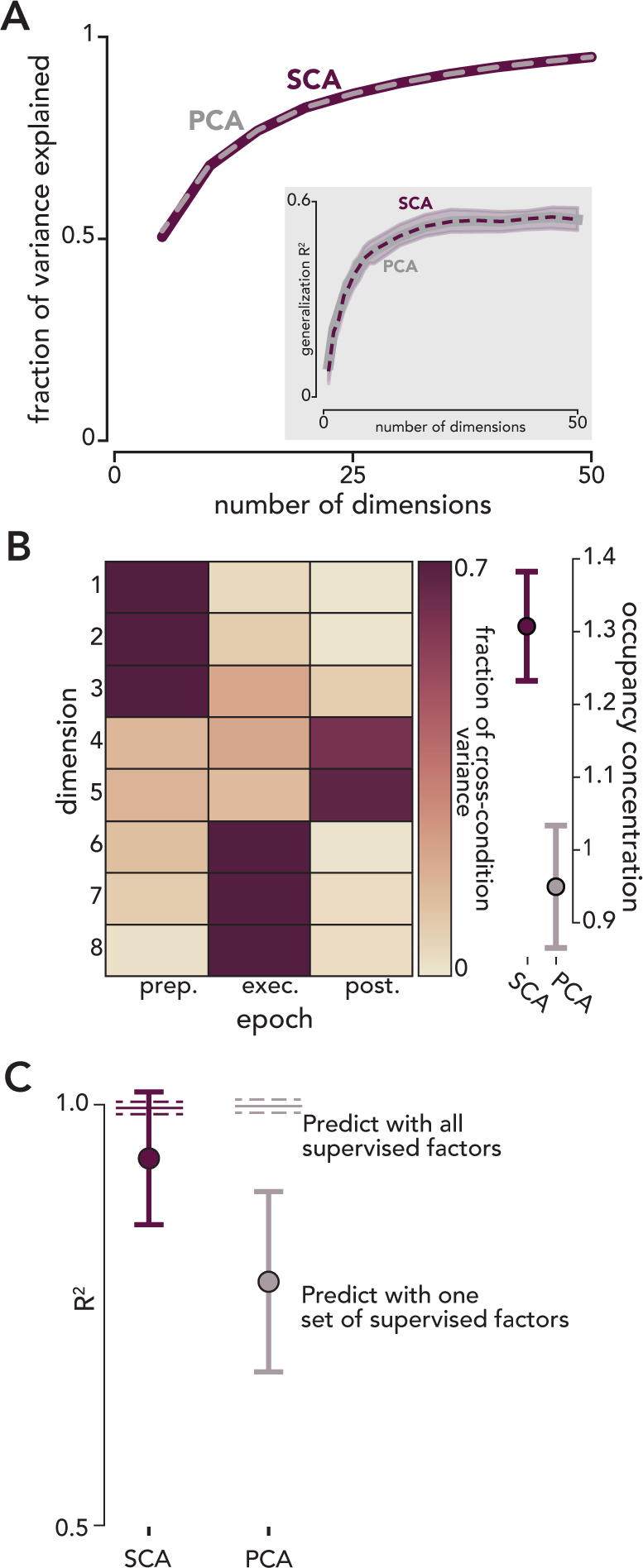
Quantitative comparisons of SCA and PCA when applied to reaching data. **A.** SCA and PCA account for virtually identical fractions of neural variance. Inset. SCA and PCA were trained on the activity from a subsample of neurons during all but one condition. R^2^ was calculated between the predicted and actual activity of held-out neurons during held-out conditions (see *Methods*). *Solid* and *dashed traces* correspond to the mean R^2^ across conditions (PCA and SCA, respectively) and *shading* indicates standard error. **B.** Left: Fraction of total occupancy (cross-condition variance) accounted for by each epoch, for each SCA dimension. Right: Mean and standard deviation of occupancy concentration (see *Methods*) calculated across bootstrapped neural populations (p < 0.01, n=100 resampled populations). **C.** Reconstructing a single SCA or PCA factor from preparatory, execution, or posture-related factors. *Error bars* correspond to mean and standard deviation across resampled populations (n=100). Reconstructions of SCA factors were significantly higher than those of PCA factors (p < 0.01, *bootstrap test*). *Horizontal lines* correspond to reconstruction performance when using all sets of supervised factors (preparatory, execution-related, and posture-related) to reconstruct each SCA or PCA factor. The difference in R^2^ between predicting factors using a single set of supervised factors vs. all supervised factors (i.e., the difference between the *solid horizontal line* and the *filled circle*) was significantly smaller for SCA than PCA (p < 0.01, *bootstrap test*).

To quantify the degree to which SCA identifies latents related to a single computation, we measured the cross-condition variance (i.e., ‘occupancy’, see *Methods*) of each dimension during preparatory, execution, and posture-related epochs (Fig. 3B, left). If a factor is only active (i.e., that dimension is ‘occupied’) during a single task epoch, then the occupancy will be high during one epoch and low during the other two. Indeed, all SCA dimensions were primarily occupied during a single epoch; in all 8 dimensions, a single epoch accounted for greater than 60% of that dimension’s total occupancy. Across all dimensions, occupancy was significantly more concentrated in SCA factors than wPCA factors (Fig. 3B, right, p < 0.01 *bootstrap test*). This result did not trivially follow from SCA prioritizing sparsity. We performed a control analysis, where we trained SCA and wPCA only on delay-period activity, divided the delay period into three ‘epochs’, and calculated the occupancy concentration of the resulting factors. Here, occupancy concentration did not significantly differ between the SCA and wPCA factors (p = 0.68, *bootstrap test*). SCA cannot recover non-synchronous factors when they do not exist. Excepting the very end of the delay period (when execution-related factors become active), only preparatory factors are active during the delay period^26,27^, so performing SCA and wPCA on delay-period activity produces factors that are comparably sparse.

To directly compare SCA factors to those recovered by a supervised method, we used the method introduced by Elsayed *et al.*^26^ to identify three orthogonal subspaces, one for each task epoch (Fig. 3C). We wish to know whether SCA naturally discovers the same divisions – factors specific to preparation, execution and posture – sought by the supervised method. If so, then each SCA factor should be a linear combination of supervised-method factors that are restricted to be of the same type (e.g. only preparation-epoch factors or only execution-epoch factors). This was indeed the case. Across all SCA dimensions, the mean best fit was 0.93±0.04 (R^2^±std). This fit was just slightly worse than that achieved with no restrictions: fitting with all sets of supervised dimensions yielded an R^2^ of 0.99. Thus, SCA found a very similar division of factors as the supervised method, despite having no prior knowledge of epoch boundaries (or even how many epochs there might be). wPCA, which is not designed to find sets of distinct factors, found more ‘mixed’ factors. Repeating the analysis above, the fit to PCA factors was 0.79±0.04 when restricted to supervised factors of a single type, versus 0.99 when using all supervised factors. In addition to better matching supervised-factors, SCA also provides more consistent factors across resampled neural populations (Fig. S2).

SCA performed similarly across a range of hyperparameters. SCA has three hyperparameters: λ_sparse_, which scales the penalty for non-sparse latent factors, λ_orth_, which scales the penalty for non-orthogonal decoding dimensions, and a requested number of latent factors. We found that the main result from the reaching datasets – that preparatory, execution, and posture-related processes occur in largely non-overlapping subspaces – was robust to relatively large changes in all hyperparameters (Figure S3). The largest difference that we observed with changes in hyperparameters was whether activity related to outwards and return reaches was captured within the same or different dimensions (Figures S4, S5). For example, consider the execution-related dimensions from monkey B (Fig. 2C, *dimensions 6, 7, 8)*. Requesting eight dimensions produces latents where execution-related activity for outward and return reaches is captured within the same dimensions, but requesting twenty-four dimensions produces dimensions where these patterns are somewhat segregated (Figure S4). While many dimensions are occupied for both outward and return movements (e.g., Figure S4 *dimensions 5*, *6*, and *15*), there are others that are only occupied during the outward reach or return reach (e.g., Figure S4, *dimension 7 and 14)*.

SCA can segregate activity related to outward and return movements because the neural spaces that are occupied during these two reaches are only partially overlapping. The return reaches are notably slower than the outward reaches (29% and 52% longer durations and 30% and 49% lower peak speeds, monkeys B and A, respectively). The alignment index is a concise method for measuring the overlap between two k-dimensional spaces^26^. Two spaces that are completely orthogonal have an alignment index of 0, while two spaces with perfect alignment have an alignment index of 1. The alignment index between delay-period activity and outward-reach related activity is 0.1, in agreement with previous results^26,27,30^. In contrast, the alignment index between outward-reach activity and return-reach activity is 0.43. SCA reflects the true alignment between activity during these task periods. Regardless of the number of dimensions requested, preparatory and execution-related factors are segregated into different dimensions, which is the desired behavior given the near-orthogonality of the dimensions occupied during preparation and execution. On the other hand, during outward and return reaches, neural activity occupies partially overlapping neural dimensions, and thus the degree to which outward- and return-reach related activity are separated by SCA depends on the number of dimensions requested. If few SCA factors are recovered, activity relating to outward and return reaches is largely placed within the same dimensions, while requesting a greater number of dimensions leads to greater separation.

Importantly, SCA, unlike alternative methods that also yield temporally sparse factors, does not seem to find ‘overly-sparse’ factors – i.e., prioritizing sparsity does not lead to a distorted view of the data. Independent component analysis (ICA) is an unsupervised dimensionality reduction method that seeks to minimize the mutual information between factors^49^ and often yields factors that are sparse in time^50^. In reaching data, we found that the sparsity of a given ICA factor is heavily influenced by the total number of requested dimensions (Figure S4). When using ICA, requesting a large number of dimensions will result in variance being ‘spread out’ across all dimensions. SCA does not exhibit this behavior. The orthogonality term in the SCA cost function discourages the recovery of factors with similar loadings (Figure S4). Therefore, requesting a large number of dimensions yields factors that are qualitatively similar to those returned when fewer dimensions are requested, in addition to many factors that only reflect noise.

A related concern is that SCA might provide a distorted view of factor dynamics. Suppose factors reflect purely oscillatory dynamics: trajectories rotate from dimension-one into dimension-two, and then back into dimension-one. One might worry that SCA could distort such structure, making dynamics appear more ‘non-normal’: activity rotating from dimension-one into dimension-two, and then into dimension-three without returning^51–53^. In fact, such distortion appears minimal. We applied SCA to activity from an RNN that uses oscillatory dynamics to generate empirical patterns of EMG (Figure S6). SCA recovered the oscillatory latents. This argues that, for the monkey whose data is shown in Fig. 2E, there is indeed a large ‘non-normal’ component to the dynamics, with activity rotating from each dimension into the next, and mostly not returning. In contrast, for an additional reaching monkey, SCA recovered strongly oscillatory latent factors (Figure S6). This suggests that the exact form of the rotational dynamics during reaching is heterogeneous across animals and that SCA can help identify this heterogeneity (see Figure S6 for further discussion).

### Unimanual Cycling

Motor cortex activity during delayed center-out reaching was a natural choice for validating SCA because the distinct component processes that produce a reach have been characterized using supervised methods. We could thus ask whether SCA recovered factors that reflect the known computational division of labor. We now turn to a behavior, cycling^54^, where less is known about potential divisions. Unlike delayed reaching, which involves brief (∼250 ms) movements whose timing is tightly controlled by the task requirements, cycling movements unfold over multiple seconds. Here, the number, or even existence, of distinct sets of computationally distinct factors has not been established.

In this task, monkeys used a hand-held pedal to traverse a virtual track (Fig. 4A). On each trial, the monkey performed either a seven, four, two, one, or one-half cycle movement to travel from a starting position to a final target. The monkey performed forward and backward cycles in multi-trial blocks, with the virtual environment cuing the monkey as to the required cycling direction. Each trial began with the pedal in one of two positions, either at the top of the cycle or the bottom (‘top-start’ and ‘bottom-start’, respectively). As in the reaching task, each trial began with a variable delay period. During this epoch, a target appeared in the distance, conveying the number of cycles necessary to complete the trial.

**Figure 4.**
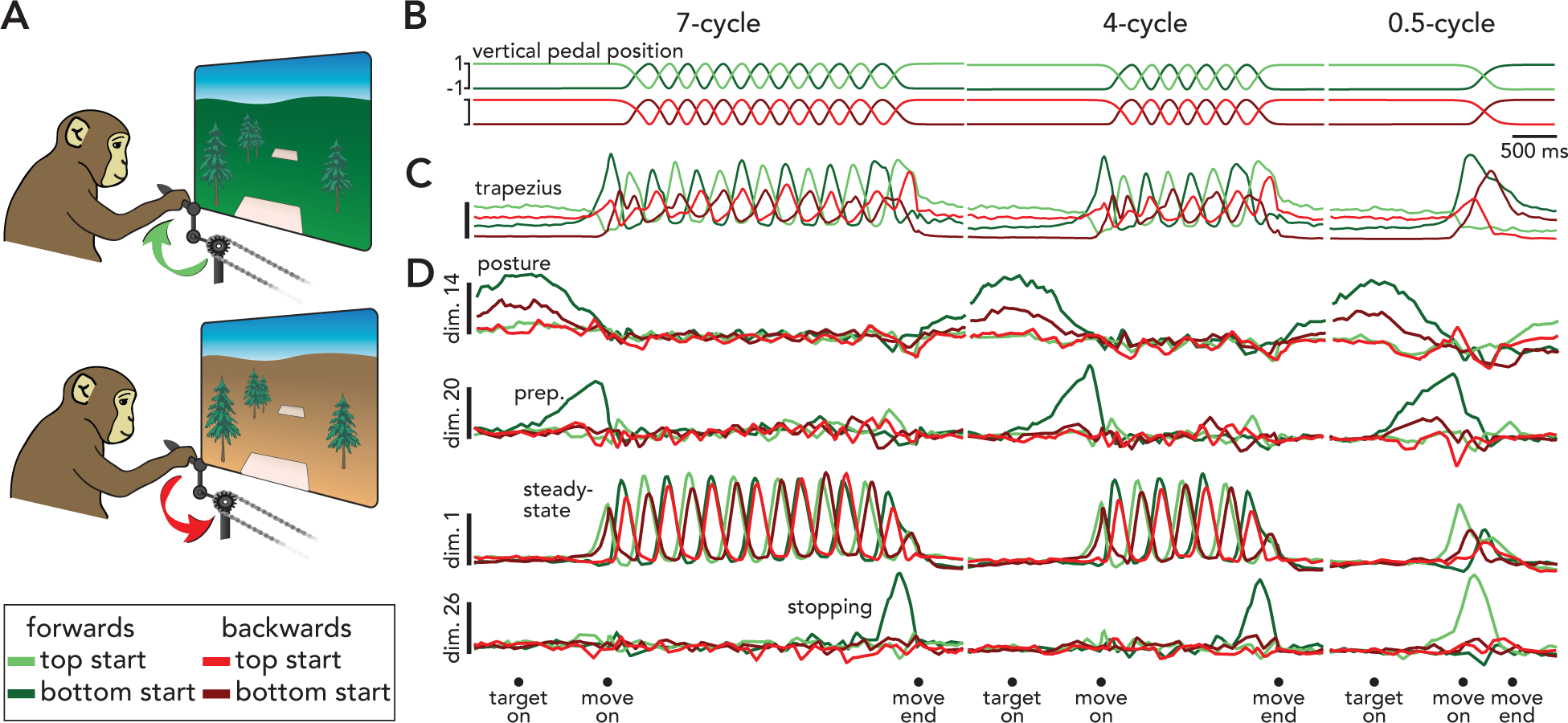
SCA identifies preparatory, execution-related, posture-related, and stopping-related factors in motor cortical unimanual cycling data. **A.** Top and Middle: Task cartoon. Monkeys pedaled forward (top) or backward (middle), pedaled for seven, four, two, one, or half a cycle, and began each trial at either the top of a cycle or the bottom. Bottom: legend for **B-D**. **B.** Left: vertical pedal position during four conditions: seven-cycle, forward (*green traces*) and backward (*red traces*), top-start (*light traces*) and bottom-start (*dark traces*). Center: vertical pedal position during four four-cycle conditions. Right: pedal position during four half-cycle conditions. **C.** Trial-averaged EMG activity from the trapezius muscle. Same format as **B**. **D.** Projections in example putative posture-related (dimension 14), preparatory (dimension 20), steady-state (i.e., execution-related, dimension 1), and stopping-related (dimension 26) SCA dimensions.

We applied SCA to trial-averaged motor cortical activity from all twenty conditions. The recovered factors fell into four categories, putatively related to postural maintenance, movement preparation, steady-state cycling, and stopping (*top* to *bottom* in Fig. 4D). One example factor from each category is plotted in Fig. 4D. All SCA factors are plotted in Figure S7. Note that SCA does not assign factors to different categories, nor does the method even ‘know’ if distinct categories exist. This illustrates an appealing feature of SCA; by simply prioritizing sparsity, the method can find factors that naturally cluster into categories amenable to interpretation. This clustered structure is not imposed by SCA, but is inherent to the data and revealed by SCA. In the case of reaching, factors clustered into categories (preparatory, execution, posture) that agreed with prior analysis and interpretation. For cycling, less is presently known and the labels presently applied to each set of factors should thus be considered tentative and descriptive.

For example, we refer to factors like that in the top row of Fig. 4D as ‘posture’ factors, because they are active only when the monkey maintains its arm at a certain position. Whether such factors generate postural muscle activity and/or reflect proprioceptive feedback remains to be determined. Yet even in advance of determining the source of the posture-related information, inspection reveals a consistent relationship between posture factors and the specific posture being held. For example, the ‘posture’ factor in Fig. 4D is most active when the animal’s arm is stationary at the bottom of a cycle, which occurs at the start and end of bottom-start, 7-cycle and 4-cycle conditions (represented by the *dark red* and *dark green* traces). At the end of 0.5-cycle conditions, this pattern appears to reverse, with bright green and red traces (top-start forward and backward, respectively) showing the greatest activity. This pattern makes sense: for the 0.5-cycle conditions, top-start conditions end at the bottom of the cycle.

The second set of SCA factors likely reflect preparatory processes (e.g., Fig. 4D, dimension 20). As during reaching, preparatory factors become active (and differentiate among conditions) during the delay period, peak approximately 140 ms before movement onset, and remain inactive during movement. The preparatory state just prior to movement onset was similar regardless of the number of upcoming cycles. This is true not only for factor 20 (Fig. 4D), but for all preparatory factors (Figure S7). This observation agrees with prior work: motor cortex activity does not reflect behavior that will occur multiple seconds in the future, in contrast to the supplementary motor area^55^. Notice that activity in preparatory dimensions remains low even after the movement ends, whereas activity in the posture dimensions tends to increase at the end of the trial.

SCA also recovers factors related to steady-state cycling (e.g., Fig. 4D, dimension 1). These factors are quiescent during the delay period and become active at approximately the same time that activity in the preparatory dimension begins to fall – analogous to the evolution of preparatory and execution factors during reaching. As in this example, steady-state factors were similarly active for top-start and bottom-start conditions, with responses that were 180 degrees out of phase, consistent with behavior (Figure S7, *light* and *dark traces*). This is consistent with the observation that the steady-state factor-trajectories are similar regardless of starting position^54^ and illustrates that SCA does not separate conditions into different dimensions if the data do not support that division. In contrast, forward and backward cycling occupied overlapping, but not identical, subspaces. Some steady-state factors (Fig. 4D, dimension 1) are active during both, while others are primarily active for only one direction (e.g., Figure S7, *dimension 3*). In agreement with the above observations, the alignment index was high when comparing top-start and bottom-start cycling (0.85±0.05, mean and standard deviation across pairs of top-start/bottom-start conditions) and smaller when comparing forward and backward cycling (0.40±0.03).

The final set of SCA factors were only active during the final cycle of a movement. For example, the factor in dimension 26 (Fig. 4) was active only when the animals stopped at the bottom after cycling forward. For 7- and 4-cycle bouts, this occurred during bottom-start conditions (due to the integer number of cycles). During 0.5-cycle bouts, this occurred during top-start conditions (which start at the top of the cycle and end at the bottom). SCA does not find separate dimensions for each individual cycle – e.g., no factors were active only during the third cycle of a seven-cycle bout. Nor were there factors that were active for 7-cycle bouts but not 4-cycle bouts. This is consistent with the prior finding that motor cortex activity repeats across steady-state cycles (i.e., all but the first and last cycle) regardless of distance^55^. This again illustrates that SCA does not create separation where none exists. The presence of a distinct stopping dimension is suggested by prior observations^55^ and agrees with analyses based on the alignment index (Figure S8). Alignment is high across all steady-state cycles and considerable when comparing initial and steady-state cycles. Alignment is much lower between steady-state cycles and the final cycle. Thus, SCA usefully identifies a feature – factors active only when stopping – whose existence is confirmed by additional analyses. SCA produced similar results for a second monkey (monkey D, Figures S9, S10): factors naturally fell into posture-related, preparatory, steady-state, and stopping-related categories (Figure S10).

In cycling, as in reaching, SCA provided a clean segregation of neural activity into groups of interpretable factors. Three of these groups (posture-related, preparatory, and steady-state) mirror the sets of factors recovered from reaching (posture-related, preparatory, and execution-related). Additionally, SCA recovered a fourth category, stopping-related factors. The existence of these stopping factors may not have been predicted from behavior alone, yet their relationship to behavior is clear.

### Limitations of SCA

We pause here to offer two practical caveats. The first is to note the instances when SCA will (and won’t) be successful. In both reaching and cycling, SCA was able to parse neural activity into preparatory, execution-related (steady-state), and posture-related factors because the factors related to these processes were non-synchronous: these processes did not *always* occur at the same time. Consider the idealized data shown in Fig. 1F. Here, there was significant overlap between when the true factors were active, yet SCA was able to recover these factors because for any pair, there were times when one factor was active and the second was not. Similarly, in the reaching dataset (Fig. 2E), SCA was able to separate posture-related factors from execution-related factors even though posture-related activity (particularly the activity in dimension 4) emerged at the same time as execution-related activity (dimensions 6-8). SCA cannot accurately segregate two computationally-distinct factors if those factors are perfectly co-active. For example, prior work^54^ has demonstrated that the largest signals during steady-state cycling primarily serve an internal computational role: maintaining low-’tangling’. Other factors, identifiable via regression, are more directly related to producing descending motor commands. These two sets of factors (‘low-tangling factors’ and ‘muscle-readout factors’) are close to orthogonal and serve two distinct computational purposes. However, SCA has no way of disentangling these two components as they occur simultaneously at all times, across all conditions.

The second consideration involves interpreting SCA factors. Even if SCA successfully parses a given factor such that it only reflects a single computational role, the precise mechanistic role that factor plays can be ambiguous, especially if that factor is only observed during a small number of self-similar experimental conditions. As a concrete example, we return to motor cortical activity during a reaching task^27^. In this task, the monkey performed center-out reaches either with or without a delay period. If we apply SCA only to activity recorded during non-delayed reaching, it is unclear whether there is preparatory activity (Figure S11). There exist two factors (1 and 2) that appear likely to be related to preparation yet could just as easily reflect early execution-related activity. Only when activity from delayed conditions is projected into the same dimensions does it become clear that these factors do indeed reflect preparatory computations. Thus, even though SCA correctly separated preparatory from execution-related factors using only non-delayed conditions, additional data (from delayed conditions) was required to know how to group factors based on their functional role. This highlights the fact that while SCA often recovers factors that are amenable to interpretation, it does not itself supply that interpretation. This is very much intended: the goal of SCA is to remain unsupervised, while still aiding interpretation in ways that previously required the use of more supervised methods.

### Bimanual cycling

In the reaching and cycling data above, SCA identified factors that were active at different times within a trial, presumably reflecting different computational roles (e.g., Fig. 1A,B). We now ask whether SCA can identify factors that are potentially differentially active across experimental conditions (e.g., Fig. 1C,D). We analyzed data from two monkeys performing a bimanual version of the cycling task^38^. Each trial required the monkey to perform seven complete cycles, cycling either forward or backward, starting at either the top or bottom of the cycle. Depending on the condition, the monkey cycled with the left arm, right arm, or both arms simultaneously (Fig. 5A,B). Monkeys performed two types of bimanual movements: with the arms in phase (bimanual-0) and with the arms 180° out of phase (bimanual-180).

**Figure 5.**
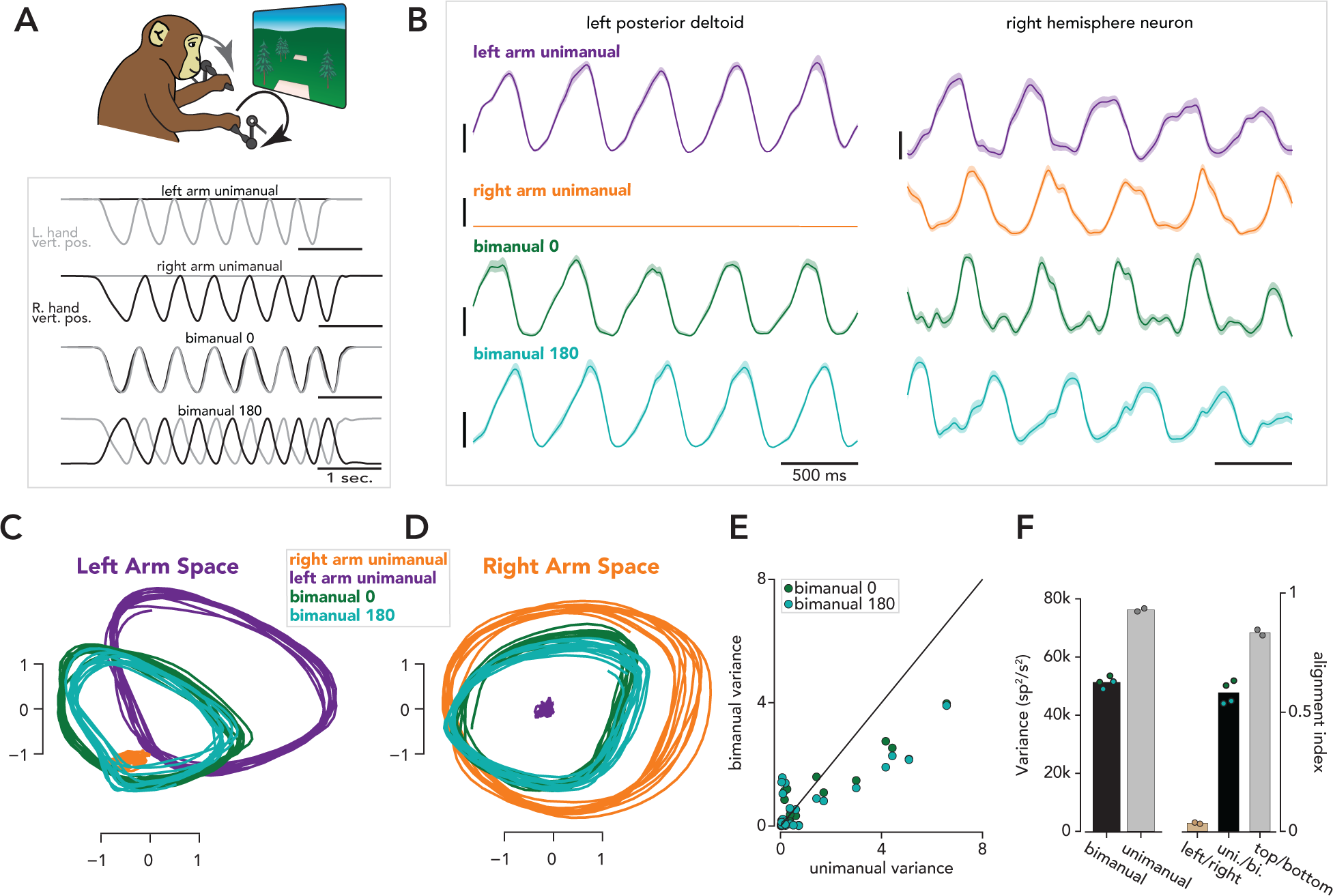
SCA identifies distinct left-arm and right-arm factors during unimanual cycling and partial overlap in the factors active during unimanual and bimanual cycling. **A.** Task schematic (top) and trial structure (bottom). Monkeys performed seven-cycle movements with either the left arm only (left arm unimanual), right arm only (right arm unimanual), both arms with a 0 degree phase offset between arms (bimanual 0), or a 180 degree phase offset (bimanual 180). Conditions either began at the top or bottom of a cycle. Traces correspond to hand positions during four example trials. **B.** Left. Trial-averaged EMG (mean and standard deviation) from the posterior deltoid during four conditions. Right. Example motor cortex neuron. **C.** All forward conditions plotted in a space that is spanned by the two SCA dimensions that accounted for the largest fraction of left arm unimanual variance. **D.** Same as **C**, but for SCA dimensions that captured a large fraction of right arm unimanual variance. **E.** Bimanual and unimanual variance in 50 SCA dimensions. **F.** Left: total neural variance during bimanual and unimanual conditions. *Blue* and *green* dots correspond to variance during a single bimanual condition, and *gray dots* correspond to the sum of the variances during left arm unimanual and right arm unimanual conditions. The two dots indicate variances during top-start and bottom-start conditions. Bar heights correspond to means across dots. Right: Alignment indices between left and right unimanual conditions (*tan*, each dot corresponds to a pair of top start/bottom start conditions), unimanual and bimanual conditions (*black*, each dot corresponds to the comparison between a single bimanual condition and a pair of unimanual conditions), and top start and bottom start unimanual conditions (*gray*, each dot corresponds to a top start/bottom start pair).

Neural activity was recorded from the primary motor cortices of both hemispheres. As illustrated by the example neuron in Figure 5B, most M1 neurons are strongly active during both ipsilateral and contralateral movements^38,39,56,57^. At the same time, it is well-established that the impacts of M1 microstimulation and inactivation are almost purely contralateral. A potential resolution to this seeming paradox is that left-arm and right-arm signals may be carried by distinct population-level factors that have distinct downstream effects^38,39^. Can SCA identify putative candidate factors, without knowing the underlying hypothesis and accompanying condition-labels?

We applied SCA to steady-state activity (i.e., activity from cycles 2-6) from all recorded neurons, during all 16 conditions (two cycling directions by two starting positions by four handedness requirements). The factors recovered by SCA point towards a marked division between the neural processes that generate left- and right-arm movements. Figure 5C,D plots the evolution of four SCA factors in state-space. Left-arm factors (e.g., Fig. 5C) were active for all conditions during which the left arm moved (left-arm unimanual, bimanual-0, and bimanual-180 conditions, *purple, light green, and dark green traces,* respectively) and were inactive when the right arm moved alone (*orange traces*). Right-arm factors (e.g., Fig. 5D) were active for all conditions during which the right arm moved (right-arm unimanual, bimanual-0, and bimanual-180, *orange, light green, and dark green traces,* respectively) and were inactive when the left arm moved alone (*purple*). This near-complete arm specificity was true of most SCA dimensions; if a factor was active when the left arm moved alone, it was rarely active when the right arm moved alone (Figure S12). The existence of arm-specific factors agrees with prior findings from both cycling^38^ and reaching^39^.

SCA revealed that the same factors that drive unimanual cycling also contribute to bimanual cycling. Factors active when the left-arm moves alone (e.g., Fig. 5C, *purple traces*) are also active during both bimanual conditions (Fig. 5C, *light and dark green traces*). The same is true for right-arm factors (Fig. 5D). A similar result has recently been observed during reaching^40^. SCA recovered additional factors that were only active during bimanual cycling (Figure S12). The set of bimanual-only factors reflected the form of coordination: bimanual-0 versus bimanual-180. Interestingly (and consistent with prior reaching results^40^) total neural variance during bimanual conditions is lower than the sum of the variance during the component unimanual conditions (Figure 5F, *left*). This was true of most individual dimensions as well (Figure 5E) and is reflected in the modestly smaller bimanual-condition orbits (*dark and light green trajectories*) in Figure 5C,D.

The SCA factors imply a specific computational division of labor. Computations related to movements of the left or right arm are separate and distinct, as suggested by the existence of distinct left-arm and right-arm factors (Fig. 5C,D). These computations can occur separately (as is the case during unimanual movements) or simultaneously (as is the case during bimanual movements). SCA also suggests that the computations that produce bimanual movements are similar, yet not identical, to those that produce unimanual movements, as evidenced by the existence of ‘bimanual-only’ factors and the fact that factors active during unimanual movements are also active during bimanual movements. Importantly, SCA did not find unique sets of factors for all possible distinctions. For example, SCA did not recover any ‘top-start only’ or ‘third-cycle only’ factors, consistent with the results in Figure 4. We validated the computational organization suggested by SCA using the PCA-based alignment index. When comparing the population response during left-arm and right-arm unimanual conditions, the alignment index was very low (0.03, Fig. 5F, right, *light brown bar*, computed for forward cycling), while the alignment index between unimanual and bimanual movements was higher (0.58, Fig. 5F, right, *black bar*). The alignment between top-start and bottom-start conditions was higher still (0.83, Fig. 5F, right, *gray bar*). Figure 5 presented results from forward cycling conditions in one monkey (monkey F). As noted above, SCA was trained on all 16 conditions simultaneously, and our primary findings held true both for backward conditions (Figure S12) and for a second monkey (monkey E, Figures S13, S14).

In a second monkey, SCA recovered factors relating to a behavioral idiosyncrasy. Monkey E increased his pedaling speed toward the end of each trial (Figure S14). Prior work, where cycling speed was intentionally varied, makes the strong prediction that increasing speed should be accompanied by a translation in state space, yielding a spiral trajectory^58^. Because the translation should be orthogonal to the rhythmic trajectory itself, there should exist at least one factor with a pronounced ramp, yet minimal rhythmic variation. With SCA, this predicted structure was readily identifiable (Figure S15).

### SCA identifies factors related to specific mating behaviors in *C. elegans*

In mammalian cortex, neurons tend to exhibit heterogeneous responses with ‘mixed selectivity’^59–65^, consistent with single-neuron rates reflecting random projections of population factors^21,42,66,67^. Is SCA useful in networks where neurons are more specialized? To probe the scope of SCA’s utility, we applied the method to single-trial calcium-imaging data recorded during *C. elegans* mating^68^, a complex behavior composed of discrete behavioral motifs (Fig. 6A). Prior work has mapped the individual neurons in the posterior brain of male *C. elegans* (the ganglia responsible for generating mating-related behaviors^69–71^) and found that the activity of many neurons is tightly linked to a few (or even a single) behavioral motif^68,72–74^.

**Figure 6.**
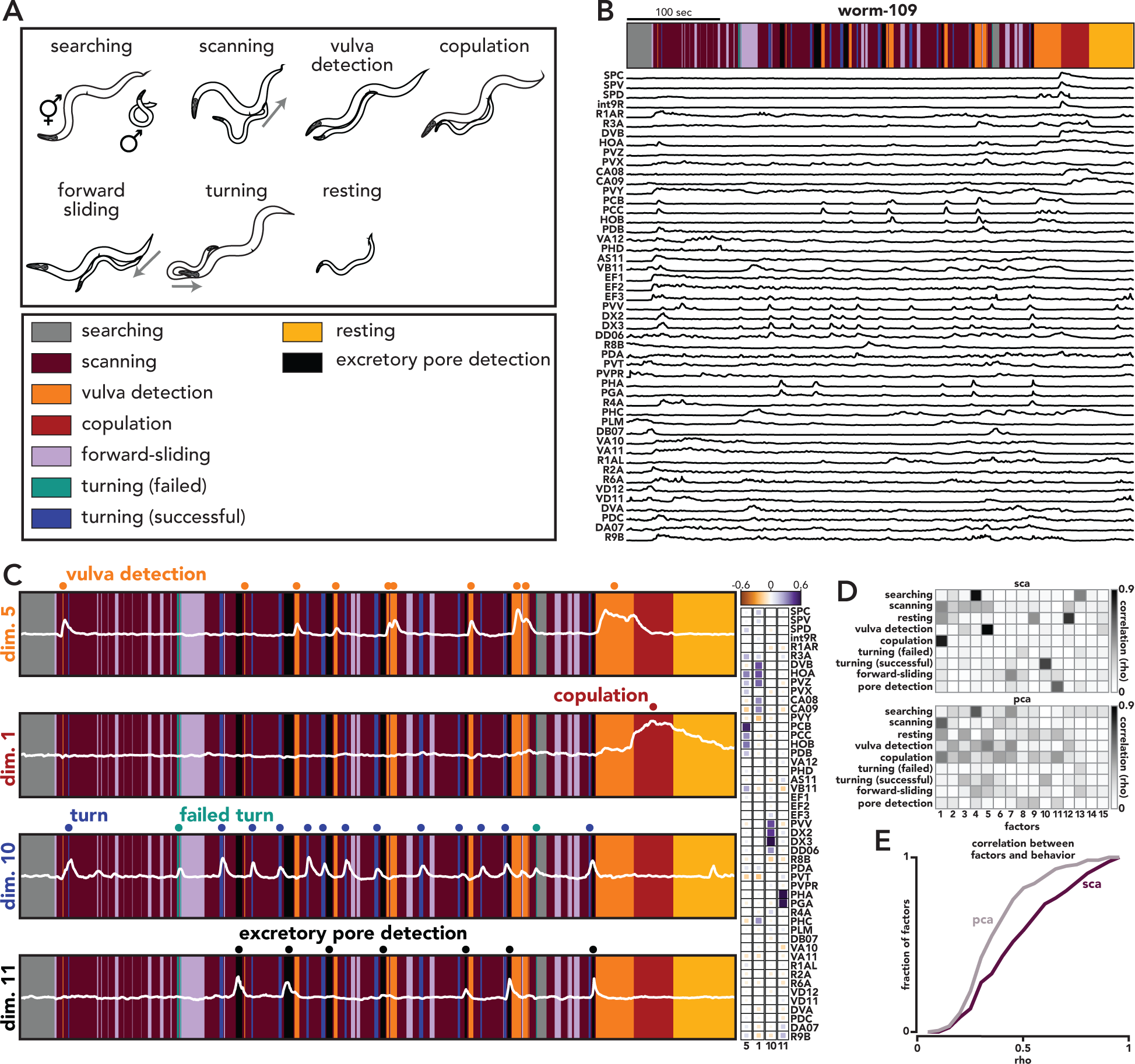
SCA identifies factors related to specific mating motifs in *C. elegans.* **A.** Behavioral motifs exhibited by worm-109. Continuous behavior was automatically classified; not all worms exhibited all behavioral motifs. **B.** Top: ethogram for a single mating bout in one worm (worm-109). Bottom: Activity from forty-nine neurons during the same mating bout. **C.** Left: Four SCA factors, all from worm-109. Activity of these four factors most closely corresponds to vulva detection, copulation, turning, and excretory pore detection (top to bottom, respectively). Right: Loadings for the four SCA dimensions plotted on the left. **D.** Correlations between all worm-109 factors and all behaviors. Top: SCA factors, Bottom: PCA factors. **E.** Maximum correlations between activity in a single SCA or PCA dimension and behavior (p < 0.001, *rank-sum test*, maximum calculated across behaviors, see *Methods*). The rightwards shift of the *purple trace* (relative to the *gray trace*) indicates that the correlations between SCA factors and behavior were higher than those between PCA factors and behavior.

We analyzed behavioral and calcium imaging data from the posterior brain of eight male worms during mating (49-54 simultaneously recorded and identified neurons, Fig. 6B, *bottom traces*). The worms’ behavior throughout the trial (i.e., a mating bout) was classified into one of twelve motifs (see *Methods*, Fig. 6B, *top*). We applied SCA to the neural activity of each individual worm separately. Note that, as in the previous analyses, SCA was not provided any task-related timing information (i.e., when different motifs occurred).

Activity in many individual SCA dimensions was tightly related to a single behavioral motif. Consider the top row of Fig. 6C, dimension 5 (for all SCA factors from all worms see Figures S16, S17). This factor’s magnitude (*white trace*) is low except around vulva detection (*orange rectangles*). The three remaining factors (Fig. 6C, dimensions 1, 10, and 11) have similarly tight relationships with the onset of copulation (*red)*, turning (*blue*), and excretory pore detection (*black*), respectively. The loadings learned by SCA are consistent with previously described functional roles of many individual neurons (Fig. 6C, *right*). Neurons PCB, PCC, and HOB are sensory neurons known to contribute to vulva detection^68,73^, and these neurons contribute heavily to the factor that is active during vulva detection (Fig. 6C, *top trace and leftmost column*). Similarly, neuron PVV contributes heavily to the factor related to turning (Fig. 6C, *third trace and third column*), in agreement with prior work, which demonstrated that PVV drives turning behavior during mating^68^.

We measured the correlation between each SCA factor and all behaviors (see *Methods*). The majority of SCA factors from worm-109 were closely related to a single behavior (Fig. 6D, *top*); most wPCA factors, on the other hand, were weakly correlated with multiple behaviors (Fig. 6D, *bottom)*. Across all SCA factors, in all worms, the mean correlation between the factors and the most closely related behavior was 0.47 ±0.02 (standard error) (Fig. 6E). Performing the same analysis with PCA factors yielded significantly lower correlations (0.37 ±0.01, p < 0.001, rank sum test), indicating that the SCA factors bore a closer relationship to the timing of individual motifs than did wPCA factors.

We next asked whether SCA could recover similar neural dimensions from different worms. If a given set of neurons are responsible for generating a specific motif across animals, then SCA should recover factors that are primarily related to that motif, and the accompanying loadings should be similar between worms. As described above, many SCA factors were tightly related to specific motifs. For example, we found SCA factors that were closely related to copulation in all worms that exhibited this behavioral motif (six of eight, Fig. 7A). Similarly, factors related to vulva detection, turning, and excretory pore detection were also common across worms (Fig. 7A).

**Figure 7.**
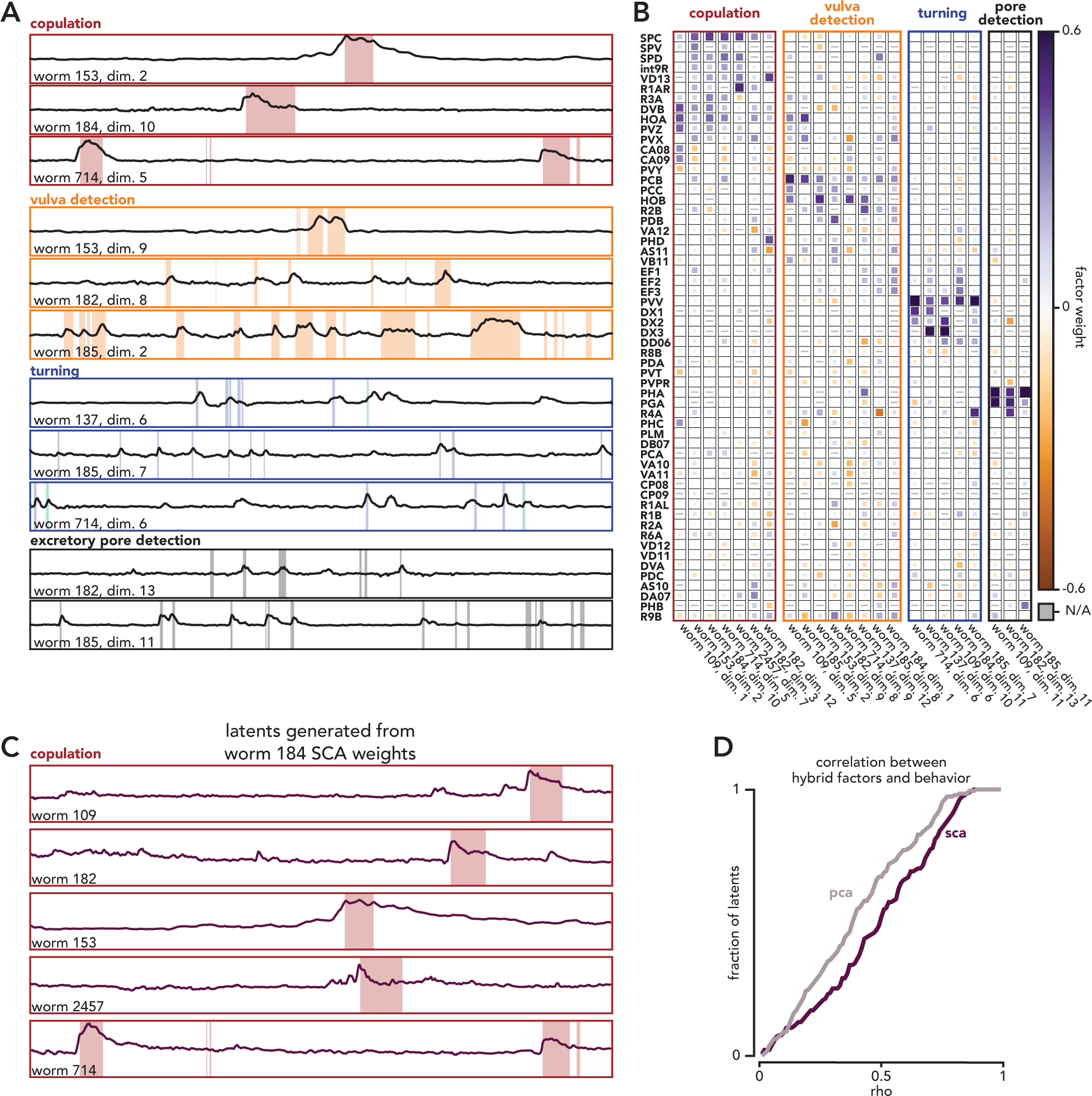
SCA identifies similar dimensions across individual worms. **A.** Example SCA factors related to specific behavioral motifs. **B.** Loadings for SCA dimensions related to the same behavioral motif. Note that there are different numbers of columns for each motif because not all worms exhibited the same motifs, and SCA did not always identify a dimension that clearly corresponded to a particular motif. The latter is partially due to different combinations of neurons being recorded across worms. **C.** ‘Hybrid factors’ generated from SCA weights from worm 184 and the neural activity of five other worms that copulated. **D.** Correlations between hybrid factors generated from SCA or PCA weights and behavior (p < 0.001, *rank-sum test*). The rightwards shift of the *purple trace* (relative to the *gray trace*) indicates that the correlations between SCA factors and behavior were higher than those between PCA factors and behavior.

The SCA loadings were also similar across worms. Qualitatively, SCA recovered similar relationships between individual neurons and particular motifs, across worms. For example, for copulation-related factors, SCA tended to learn large weights for neurons SPC, SPD, and HOA, neurons that are known to play an important role in copulation (Fig. 7B, *red box*)^68,72^. Similarly, for turning-related SCA dimensions (Fig. 7B, *blue box*), SCA recovered large weights for neuron PVV (a neuron known to be important for turning). To quantify the similarity between SCA loadings, we measured the angle between the loading vectors of individual worms (Figure S18). When compared to the angle between wPCA loadings, we found a modest but highly significant difference between SCA angles and wPCA angles (Figure S18), indicating that the SCA loadings are indeed more replicable across worms than wPCA loadings.

To further test the generalizability of the SCA loadings, we used the SCA loadings from one worm to predict the behavior of a second worm (Fig. 7C). Given an SCA factor that was related to a particular motif in one worm (worm-A) and a second worm that exhibited the same motif (worm-B), we generated a ‘hybrid factor’ using the SCA loadings from worm-A and the neural activity from worm-B. We then asked how well the hybrid factor correlated with the behavior of worm-B. For an example, see Fig. 7C. Here, we used the SCA loadings from worm-184 and the neural activity of the other five worms that copulated to generate five hybrid factors (Fig. 7C, *traces*). These hybrid factors were well correlated with copulation timing (rho = 0.59±0.13, mean and standard deviation, across worms). Across all worms and motifs, we found that hybrid factors generated from SCA loadings were significantly more correlated with behavior than the comparable factors generated from wPCA loadings (mean rho = 0.48±0.02 versus 0.39±0.02, p < 0.001, rank sum test, Fig. 7D).

### Identifying compositionality in a multitask RNN

As demonstrated above, SCA leverages the fact that many neural computations (e.g., reaching) are composed of multiple distinct processes (e.g., preparation and execution). This discrete computational structure opens the possibility of reusing component processes across multiple tasks. Component processes (i.e., component computations) could be used as ‘building blocks’ that are reassembled or reordered to perform different tasks. While such ‘compositionality’ has yet to be observed in neural data (partially due to the difficulty in training a single animal to perform multiple tasks simultaneously), reuse of component computations has been observed in artificial networks. Driscoll *et al.*^13^, extending earlier work by Yang *et al.*^35^, found that single networks trained to perform fifteen different cognitive tasks learned a small number of component computations. These computational building blocks were used across multiple tasks and even allowed networks to quickly learn new tasks. Importantly, Driscoll *et al.,* found that compositionality was ubiquitous across networks, even when such structure was obscured at the level of individual neurons. In networks composed of units with non-negative activation functions (soft plus or rectified tanh), individual units primarily participated in a small number of component computations. In tanh networks, however, units tended to be active during most component computations.

To determine whether SCA could recover the compositionality masked at the level of individual neurons, we performed SCA on the activity of a tanh network trained to perform 15 cognitive tasks (Fig. 8A, see *Methods* and Figure S19 for a thorough description of each task). In all tasks, the network received scaled, directional inputs (in the form of the cosine and sine of different angles) and generated a 2-dimensional directional output. As was previously reported^13^, the majority of neurons were active during most tasks (Fig. 8B).

**Figure 8.**
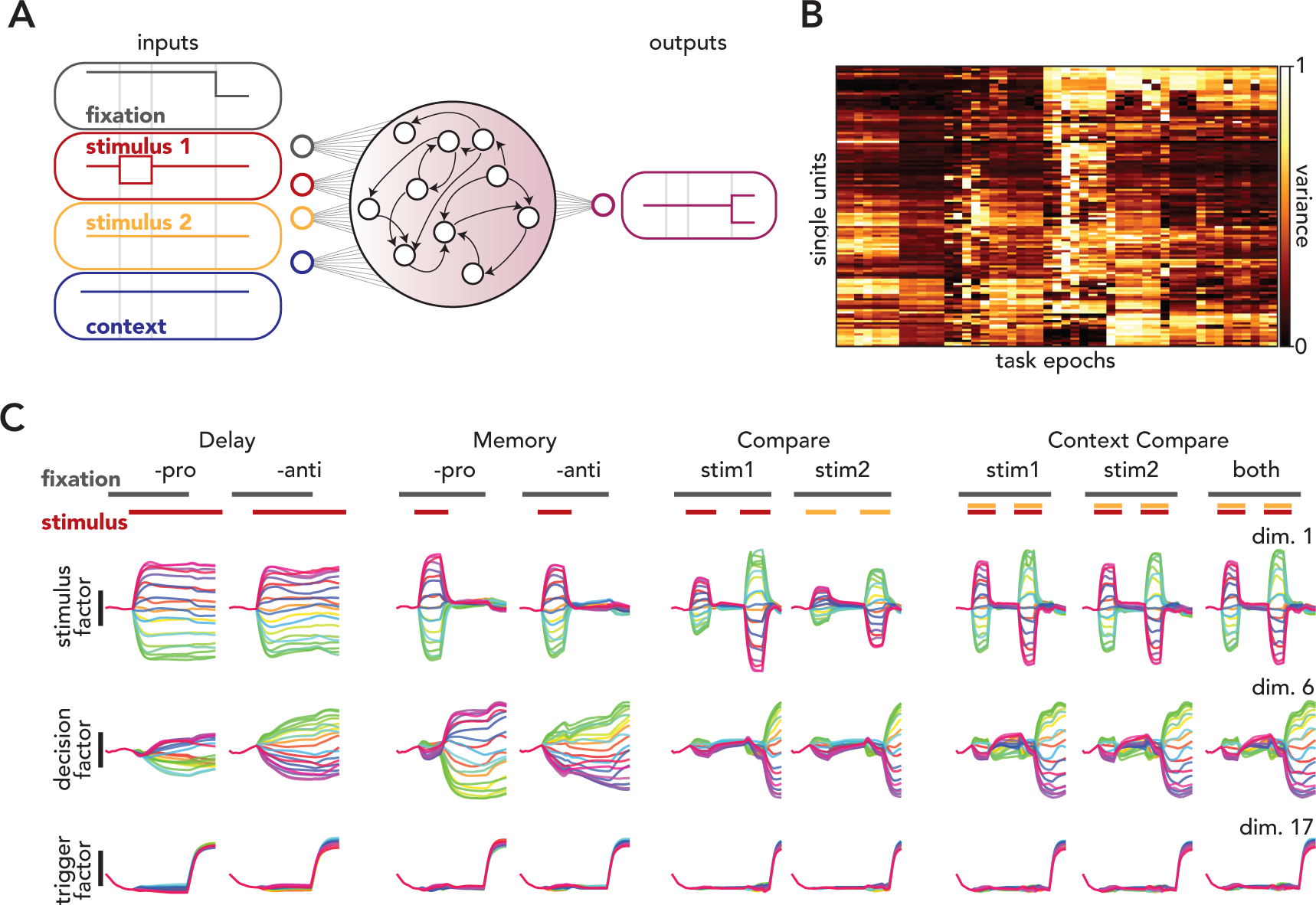
SCA identifies compositionality in a multitask network. **A.** Network architecture. **B.** Variance of individual units across task epochs. Epochs were delineated by the timing of the network inputs, and the number of epochs differed across tasks. **C.** Activity in three SCA dimensions during nine tasks. The first factor (top row) reflected the stimulus, the second (middle row) reflected the network’s ultimate decision, and the third (bottom row) reflected the timing of the network’s output. During the ‘Delay-pro’ task, the network received a directional stimulus and generated an output in the same direction after receiving a ‘go cue’. The ‘Delay-anti’ task was identical, except the network needed to generate an output that was 180° from the input stimulus. The ‘Memory-pro’ and ‘-anti’ tasks were the same as the ‘Delay-’ tasks, except the network did not receive a continual directional stimulus. The remaining five tasks required the network to compare the amplitude of two successive stimuli and respond in the direction of the larger of the two. For ‘Compare-stim 1’ and ‘Compare-stim 2’, the network compared the amplitude of stimuli delivered via a single pair of stimulus dimensions (stimulus 1 or stimulus 2). For the ‘ContextCompare-’ tasks, the network received stimuli from both pairs of stimulus dimensions and needed to ignore one (‘ContextCompareStim 1’ and ‘ContextCompareStim 2’) or compare across both pairs of inputs (‘ContextCompareStimuli’). During network training, the duration of the stimulus periods (and memory periods, when present) varied randomly from trial to trial to prevent the network from predicting task period transitions.

The factors identified by SCA reflected the compositionality present in the network and pointed towards a computational strategy that could be used to solve the tasks. To illustrate this strategy, we highlight the activity of three SCA factors during nine tasks (Fig. 8C, for a more thorough description of all recovered factors during all tasks, see Figure S20). The network’s putative strategy could be summarized as follows: use a single set of factors to represent the stimuli (e.g., Fig. 8C, dimension 1), a second set of factors to reflect the network’s decision (e.g., Fig. 8C, dimension 6), and condition-invariant factors to determine when activity flows into output-potent dimensions (e.g., Fig. 8C, dimension 17).

Across tasks, the network received directional input via two ‘channels’ (two, 2-dimensional directional inputs, Fig. 8A, *stimulus 1 and 2*). For some tasks, the network received input from only one of these channels (e.g., Delay-pro, Compare-stim2), while in others, both stimulus channels were active (e.g., ContextCompare). SCA identified a factor that reflected the stimulus across all tasks, regardless of input channel (Fig. 8C, dimension 1). Activity in this dimension reflected the time courses of the stimuli; during the Delay-tasks, activity in this dimension was constant after the stimulus was delivered. The same dimension reflects the stimuli regardless of the input source; activity in dimension 1 is highly similar during Compare-stim1 and Compare-stim2 even though the input sources are different in these two tasks. SCA also identified dimensions that only reflected the input from stimulus channel 2 (Figure S20, dimensions 3 and 4).

A second set of factors (e.g., Fig. 8C, dimension 6) reflected the network’s ultimate output. Consider activity during the Memory tasks. Here, the network receives a brief directional stimulus. In the Memory-pro task, the network must report this same direction after the go cue is delivered (i.e., when the fixation signal is removed) while in the Memory-anti task, the network must report the direction 180° from the instructed direction. After the stimulus is removed, activity grows in dimension 6, but with opposite ordering (*purple traces* are above the *green traces* in dimension 6 during Memory-pro, while the reverse is true during Memory-anti). This ordering reflects the ultimate ‘decision’ of the network. We use the term ‘decision’ loosely here, in reference to the fact that activity in dimension 6 reflects the ultimate output of the network, even before that output is produced. During Compare- and ContextCompare tasks, where the correct output is not known until the second of two stimuli is delivered (see the legend of Fig. 8C for more details), activity remains low until the second stimulus is removed.

Finally, the network uses a third set of factors to determine the timing of the output (e.g., Fig. 8C, dimension 17). The evolution of these factors was highly stereotyped across conditions and tasks, reminiscent of the condition-invariant ‘trigger signal’ that has been observed during reaching in non-human primates^75^, decision making in rodents^43^, and speech in humans^67^. In reaching monkeys, the trigger signal translates preparatory activity to a region of state space dominated by rotational dynamics^14,15,75^. While preparatory activity is output-null (with respect to motor cortical output^26,76^, execution-related activity has a non-zero projection onto output dimensions. Likewise, dimension 6 is output-null, and it is likely that the activity in dimension 17 translates the activity in these dimensions to new dimensions (not shown) that have non-zero projections onto the network’s readout dimensions.

While these three sets of SCA factors do not offer a full description of how the network solves these tasks (this would require direct interrogation and characterization of the network dynamics as in^13^), they suggest important aspects of the computational strategy. Such a clear representation of the network’s underlying computations could not be gleaned from the activity of single units (Fig. 8B) nor from PCA (Figure S21).

## Discussion

For many networks, computation is best understood in terms of population-level factors. This is known to be true of a sizable class of network models (e.g.,^11,12,14,17^, and strongly suspected to be true of many cortical (and perhaps subcortical) areas. The idea of a factor is simple: factors are the non-noise signals present in a network – i.e., signals that are reliable enough to subserve computation. Factors cannot be directly observed; they need to be estimated from the activity of many individual neurons. If one simply wishes to recover factors that can account for the activity of individual neurons, the choice of basis set does not matter (as long as it spans the full factor space). Yet this choice may matter a great deal to an experimenter attempting to understand the system; interpretation may be greatly aided by recovering factors that can be parcellated into sets that serve somewhat different functions. This has been a long-standing goal in our field, but gaining such interpretability has, historically, required supervision, either explicitly, in the form of supervised dimensionality reduction methods (e.g.,^12,31,32,77^) or implicitly, in the form of manual parsing of the neural activity into separate task epochs (e.g.,^17,19,26,66^).

Here, we introduced SCA, an unsupervised method that has many of the advantages of supervised methods, but with less need for the experimenter to anticipate or guess at the types of structure that might be observed. We found that, when applied to a variety of datasets, SCA often identified factors that reflected a single computational role. For example, when applied to data during reaching, SCA identified factors that could be grouped into one of three sets: one reflecting preparation, one reflecting execution, and one reflecting postural maintenance. This parcellation agreed with that previously found using supervised methods^26,48^, demonstrating that SCA can be useful in situations where one had previously needed supervised methods, and confirming that prior approaches identified naturally present divisions in the data. Across a variety of datasets, we repeatedly found that SCA identified nearly orthogonal sets of factors that had different computational ‘meanings’, in the sense that were associated with a particular behavior (e.g., cycling with one arm versus the other, Fig. 5), a particular aspect of a larger behavior (e.g., stopping at the end of a bout of cycling, Fig. 4), or internal process (e.g., reach preparation, Fig. 2). At a technical level, this demonstrates the utility of SCA. At a scientific level, it supports the emerging idea that a common network strategy is to use distinct factors, occupying orthogonal dimensions, to perform distinct computations or distinct aspects of a larger computation.

An advantage of SCA, and the reason we could usefully apply it to many datasets, is that it makes minimal, but powerful, assumptions: neural activity is low dimensional, factors associated with different computational roles are non-synchronous, and such non-synchronous factors tend to occupy unaligned (nearly orthogonal) subspaces. The assumption that factors are non-synchronous is much less stringent than it might initially appear. It is not necessary that factors be active at completely different times, merely that there be some flexibility in when they occur relative to one another. For example, SCA was able to segregate left-arm and right-arm factors even though these two sets of factors are simultaneously active during both bimanual cycling conditions (Fig. 5C,D). Additionally, SCA identified stopping factors during unimanual cycling (Fig. 4D), despite the fact that, whenever the stopping factors were active, steady-state factors were also active. SCA was able to separate these two sets of factors because there were times when steady-state factors were active but stopping factors were not.

Despite using a simple cost function, SCA allowed a surprisingly clear parcellation of the factors underlying multiple behaviors. This clarity was maintained across a broad range of hyperparameters (Figures S3, S4, S5). In addition to recovering ‘expected’ factors, SCA also found previously uncharacterized factors, such as preparatory cycling factors (Fig. 4) and bimanual-only cycling factors (Fig. 5). While the computational role of these (and any other newly characterized factors) cannot be determined without further study (e.g., more/different experimental conditions), their *potential* role is clear. This interpretability is what makes SCA a powerful method; in all examined datasets, SCA yielded factors that invited hypotheses of how a given neural computation was performed.

### Interpretability

‘Interpretable’ is a label that we, as researchers, use when we are able to assign meaning to or derive meaning from a particular neural signal or set of signals. The ‘meaning’ that we ascribe to SCA factors relates to their putative computational role; SCA factors are interpretable insofar as they aid us in making testable predictions about how a particular computation occurs. By seeking sparsity and orthogonality, SCA facilitates the grouping of factors into sets that ostensibly relate to distinct computational processes. Non-synchronous factors are assumed to reflect different computational roles (e.g., preparatory and execution-related factors), while synchronous factors are assumed to perform closely related roles (e.g., two posture-related factors). The factors identified by SCA tend to form natural groups (e.g., preparatory, posture, and execution factors) to which one may be able to assign meaning. Yet SCA does not itself determine how individual factors should be grouped, nor does the method assign ‘meaning’ to a set of factors; parcellation and interpretation of factors requires understanding of the task being performed and considerable scientific knowledge. For example, the execution-related factors for monkey B (Fig. 2E, *dimensions 6-8)* are not perfectly synchronous, yet we assume that they contribute to a single computational process because they are all only active during a reach and prior work has demonstrated how non-normal dynamics can be used to generate a reach^51^. The goal of SCA is not to obviate the need for scientific reasoning, but rather to make such reasoning easier by supplying factors that are likely to lend themselves to straightforward interpretations.

Alternatively, factors can be interpretable insofar as they correlate with a particular behavior. The ‘meaning’ that is assigned to such factors does not necessarily relate to an internal computational role, but rather references the close relationship between a factor and an externally measured signal. There are a wide variety of methods that seek such factors. For example, two recent methods, pi-VAE^78^ and CEBRA^79^, use generative modeling and contrastive learning, respectively, to find low-dimensional spaces in which similar locations in state-space correspond to similar behaviors. Two additional methods, preferential subspace identification (PSID)^80^ and targeted neural dynamical modeling (TNDM)^81^, use instead a dynamical systems approach to identify ‘behaviorally-relevant’ and ‘behaviorally-irrelevant’ factors (i.e., factors that do and do not correlate with a particular behavior). Note that the goals of SCA and methods that seek behaviorally-correlated factors are typically quite different and are thus complementary. For example, in the case of the cycling data (Fig. 4), the factors identified by SCA are those that one would likely wish to use when reverse engineering the solution used by a network (biological or artificial) to produce its output. However, if one’s goal is to predict behavior from neural activity, SCA is likely not the best choice, as the factors that are active during forwards and backwards cycling are quite different^82^ (e.g., Fig. S10). A natural choice would be to use a method like CEBRA to identify a single set of factors that are highly active during both forwards and backwards cycling.

### Factors and neurons

The degree to which computationally distinct factors relate to physically distinct neurons will vary by brain area and species. In *C. elegans*, where the connectivity of individual neurons is preserved across animals, SCA identifies factors that relate to specific component processes (behavioral motifs, Fig. 6A), and many of these factors primarily reflect the activity of a small number of neurons (e.g., Fig. 6B, *excretory pore detection* factors). In this system, distinct processes are often reliant on the activity of single neurons (e.g., ‘excretory pore detection’ requires PHA activity^68^) and can be understood, at the circuit level, in terms of interactions between individual neurons. In contrast, in monkeys, there exist distinct reach-preparation factors but very few ‘preparation only’ neurons; nearly all neurons that are active during preparation are active during execution, especially if many conditions are tested. Similarly, there exist distinct left-arm and right-arm factors, but relatively few ‘right-arm-only’ neurons, even in primary motor cortex^38^. This appears to be a common property in much of cortex^21,61,66,76^. Nevertheless, SCA identifies similar factors across animals (e.g., Fig. 2 and Figure S1 and Fig. 3 and Figure S9), reflecting the fact that the underlying factors that generate a behavior can be similar across animals^83^, even if there is not a one-to-one mapping between neurons.

### Alternative strategies and related methods

SCA is specifically designed to leverage particular structure within neural activity, yet there are alternative approaches for identifying non-synchronous factors. Traditionally, non-synchronous sets of factors are identified via careful task design. Controlling when different component processes occur (e.g., preparation during a reaching task) allows a researcher to use an unsupervised method to identify factors that only reflect a particular computational role. However, even with careful task design, it can be difficult to predict when different processes occur or even how many distinct processes contribute to a particular computation. For example, it is not obvious, from behavior alone, that stopping-related factors should exist (Fig. 4).

SCA uses end-to-end optimization to identify factors that can account for large amounts of neural variance and are temporally sparse. One could instead use a step-wise strategy – i.e., first identify a set of factors, then subsequently rotate this basis set to increase the sparsity of the factors. A recent study^84^ used such an approach to characterize population-level activity in human motor cortex during real and imagined movement. Here, the authors used PCA to identify a low-dimensional space, then used a varimax rotation to maximize the variance of the projections (across factors) at each time point. While the varimax rotation does not directly optimize for sparsity, such an approach may approximate the solution found by SCA. The results of Dekleva *et al.* provide further evidence of the utility of identifying non-synchronous factors.

SCA follows in a long line of dimensionality reduction methods that incorporate sparsity for the purposes of improved interpretability. For example, sparse PCA (sPCA)^85,86^) is a well-known method that seeks factors with sparse loadings, making this method particularly useful for clustering (e.g., identifying classes of cell types on the basis of electrophysiologic properties^87^). Both sPCA and SCA can be considered to fall under the broad umbrella of a class of methods known as dictionary learning^33,34^. Dictionary learning seeks to decompose a matrix into a product of two matrices, one of which is encouraged to be sparse; prior work has even referred to dictionary learning as sparse component analysis^88^. Within neuroscience, dictionary learning is not typically used for estimating latent factors but rather as a preprocessing tool (e.g., for extracting neural activity from calcium imaging data^89–91^ or for spike sorting^92^). ICA, which does not seek sparse factors *per se*, but to minimize the mutual information between factors^49^, can also be used to recover non-synchronous factors^50^. While standard dictionary learning and ICA may perform similarly to SCA under some hyperparameter choices (e.g., if a small number of factors are requested), we believe the lack of an orthogonality term limits the interpretability of the factors recovered by these methods. As noted in Figure S4, the majority of ICA factors recovered from reaching data had appreciable variance, regardless of the number of factors requested. This behavior results from ICA not being penalized for recovering non-orthogonal dimensions; as more factors are requested, ICA produces dimensions that are more collinear (Fig. S4). Consequently, using ICA factors to delineate different computational processes would be challenging, as the apparent boundaries between such processes would differ depending on the number of requested dimensions. SCA, on the other hand, does not exhibit such ‘over-sparsification’; if one requests ‘too many’ dimensions, SCA simply returns factors that reflect noise (Fig. S4).

### Conclusions

SCA facilitates, using analysis, what was more traditionally accomplished via careful task design: the segregation of neural activity related to distinct component processes. Once the boundaries between component processes have been established, each process can be characterized individually and the relationship between processes can be determined. The assumptions that SCA makes about neural computations appear to be widely applicable; neural computations occurring in monkeys, worms, and artificial networks were rendered more interpretable via the identification of non-synchronous factors. This success suggests that SCA can be a broadly useful tool for understanding the relationship between neural activity and behavior.

## Author Contributions

A.J.Z. and J.I.G. developed SCA; A.J.Z. and A.H.L. collected the reaching data; A.A.R. collected the unimanual cycling data; K.C.A. collected the bimanual cycling data; L.D., trained the multitask network; V.S., collected the *C. elegans* data; A.J.Z., X.A., V.S., L.D., J.I.G. performed the analyses, A.J.Z., M.M.C., and J.I.G. wrote the manuscript. All authors contributed to manuscript editing.

## Acknowledgements

We thank Y. Pavlova for expert animal care. This work was supported by the Grossman Center for the Statistics of Mind (M.M.C., J.P.C., and L.P.), the Simons Foundation (M.M.C., L.D., J.P.C., and L.P.), the McKnight Foundation (M.M.C. and J.P.C.), NSF Neuronex 520 Award DBI-1707398 (L.P.), the Gatsby Charitable Foundation (L.P. and J.P.C.), NIH Director’s New Innovator Award DP2 NS083037 (M.M.C.), NIH CRCNS R01NS100066 (M.M.C.), NIH 1U19NS104649 (M.M.C.), P30 EY019007 (M.M.C.), the National Science Foundation (A.J.Z.), the Kavli Foundation (M.M.C.), NIH R01NS113119 (V.S.), NIH ROO NS119787 (J.I.G.).

## Methods

### Sparse Component Analysis

#### Parameters and Cost Function

Let **X***^T×N^* be a matrix containing the activity of *N* neurons over *T* time bins. The goal of SCA is to find a low dimensional latent representation, with dimensionality *K,* that 1) accurately reconstructs the original data **X** via an affine mapping (linear mapping **V***^K×N^* and offset ***b_v_***^1×*N*^), 2) has temporally sparse latents, **Z***^T×K^*, and 3) encourages the mapping from the latent space to neural space to be orthonormal. We additionally include optional sample weighting, via the diagonal matrix **W***^T×T^*, which allows providing different importance to different time points within the cost function.

This cost function can be written as follows, with the summed terms corresponding to the three above goals, respectively:

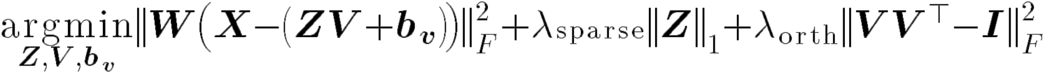

In practice, rather than directly optimizing the latents, we learn the latents via a mapping from the neural activity space (which can be viewed as a linear autoencoder, Fig. 1E).

This is accomplished with the following cost function:

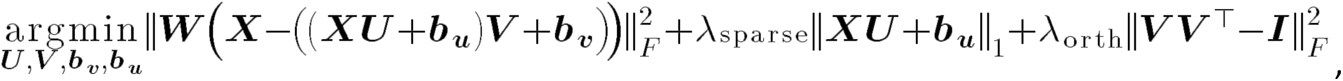

where **U***^N×K^* and ***b_u_***^1×*K*^ are the linear matrix and offset in the “encoder” part of the autoencoder, that map the neural activity to the low-dimensional space.

In addition to the orthonormality penalty, we also strictly constrain each row of **V** to have unit norm. Thus, even if λ*_orth_* is set to 0, optimization will not lead to a solution in which the magnitude of **Z** endlessly becomes smaller while **V** endlessly becomes larger.

We note that in our accompanying code package, we also include an option to enforce strict orthogonality in the recovered dimensions by performing manifold optimization^93^. However, we have found that using the orthogonality penalty (as opposed to an orthogonality constraint) is advantageous for two primary reasons. First, while the two options generally produced very similar factors, we found that, on occasion, constraining SCA to recover orthogonal dimensions could produce factors that were more ‘mixed’. For example, the separation between posture-related factors and preparatory factors was less complete for monkey C (Fig. S7) if we required SCA to recover orthogonal dimensions. The second benefit was purely practical; fitting SCA with manifold optimization was notably slower. For example, fitting SCA on the reaching dataset from monkey B was slowed by a factor of 1.8 if we used a strict orthogonality constraint.

#### Sample weighting

When we use sample weighting, we default the sample weight at a given time point to be inversely proportional to the sum squared activity of the neural population at that time point. This has the purpose of encouraging the model to still use latent dimensions to reconstruct lower-activity time points (e.g. to care as much about motor preparation as motor execution, when the former has substantially lower activity).

#### Optimization

All optimization is done in PyTorch with the Adam optimizer^94^ using default hyperparameters. We use learning rate scheduling, where the initial learning rate is set at 1e-3, and the learning rate is halved every time the loss does not decrease for 100 iterations, until a minimum learning rate of 5e-4.

#### Initialization of parameters and hyperparameters

We initialize the model parameters using sample-weighted PCA on the data. That is, if **Q***^N×K^* are the sample-weighted PCA loadings, we initialize with **U** = **Q**, **V**=**Q***^T^*, ***b_v_***=0, ***b_u_***=0. In practice, we find nearly equivalent solutions using a random initialization of parameters, but convergence generally takes significantly longer. Both sample-weighted PCA and random initializations are available within our code package.

In our code package, we have the following default settings for model hyperparameters. λ_orth_ is initialized so that, if the off-diagonal entries of ***VV***^T^ were all equal to 0.1, the orthonormality penalty would be 10% of the initial reconstruction cost (with sample-weighted PCA initialization). λ_sparse_ is initialized so that the initial cost associated with the sparsity penalty is 10% of the initial reconstruction cost. This serves to make sparsity and orthogonality important, but not at great expense of reconstructing the neural data. These defaults allow for easy out-of-the-box model-fitting without hyperparameter tuning. Still, we have found that it is often slightly beneficial to increase λ_sparse_ beyond this default value - i.e., further sparsity can be achieved without harming the reconstruction accuracy. Thus, for the demonstrations in this paper, we sometimes set λ_sparse_ above the default value - using λ_sparse_ values at the elbow of the plot of reconstruction accuracy versus λ_sparse_. We also provide a code notebook so that users can also find λ_sparse_ in this way. See below for a table of the hyperparameters used for each dataset.

#### Advantages of neural-network based optimization

We chose to optimize SCA as a linear autoencoder using standard neural network machinery for two primary reasons. 1) This framework scales well with increased dataset duration. The number of encoding and decoding weights depends only on the number of neurons and number of requested factors, whereas directly optimizing latents yields a parameter count that scales with the number of timepoints. Additionally, neural network machinery can easily allow for GPU use, providing improved speed for larger datasets. 2) Our approach makes the method readily extendable. For example, if a dataset contains neural activity from two brain regions, the model can be easily altered to identify interpretable factors within one brain region that predict activity in the second region. Additionally, one can easily change the model’s cost function (e.g. including Poisson statistics) or architecture (e.g. adding nonlinearities). This flexibility will encourage further development of interpretable dimensionality reduction tools for neural population activity.

### Sample-weighted Principal Components Analysis

Sample-weighted PCA (wPCA) is a variant of principal component analysis that includes providing a weight for each sample (time point). Formulated as a cost function, it aims to solve:

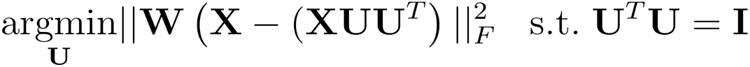

**U** is found by running singular value decomposition (SVD) on the matrix **WX**, in the same way that standard PCA runs SVD on the matrix **X**.

### Datasets

All analyzed datasets (center-out reaching^27^, unimanual cycling^54^, bimanual cycling^38^, center-out reaching network^14^, multi-task network^13^, and *C. elegans* mating^68^) have previously been analyzed. Detailed descriptions of each can be found in the respective citations. Here, we briefly describe each dataset.

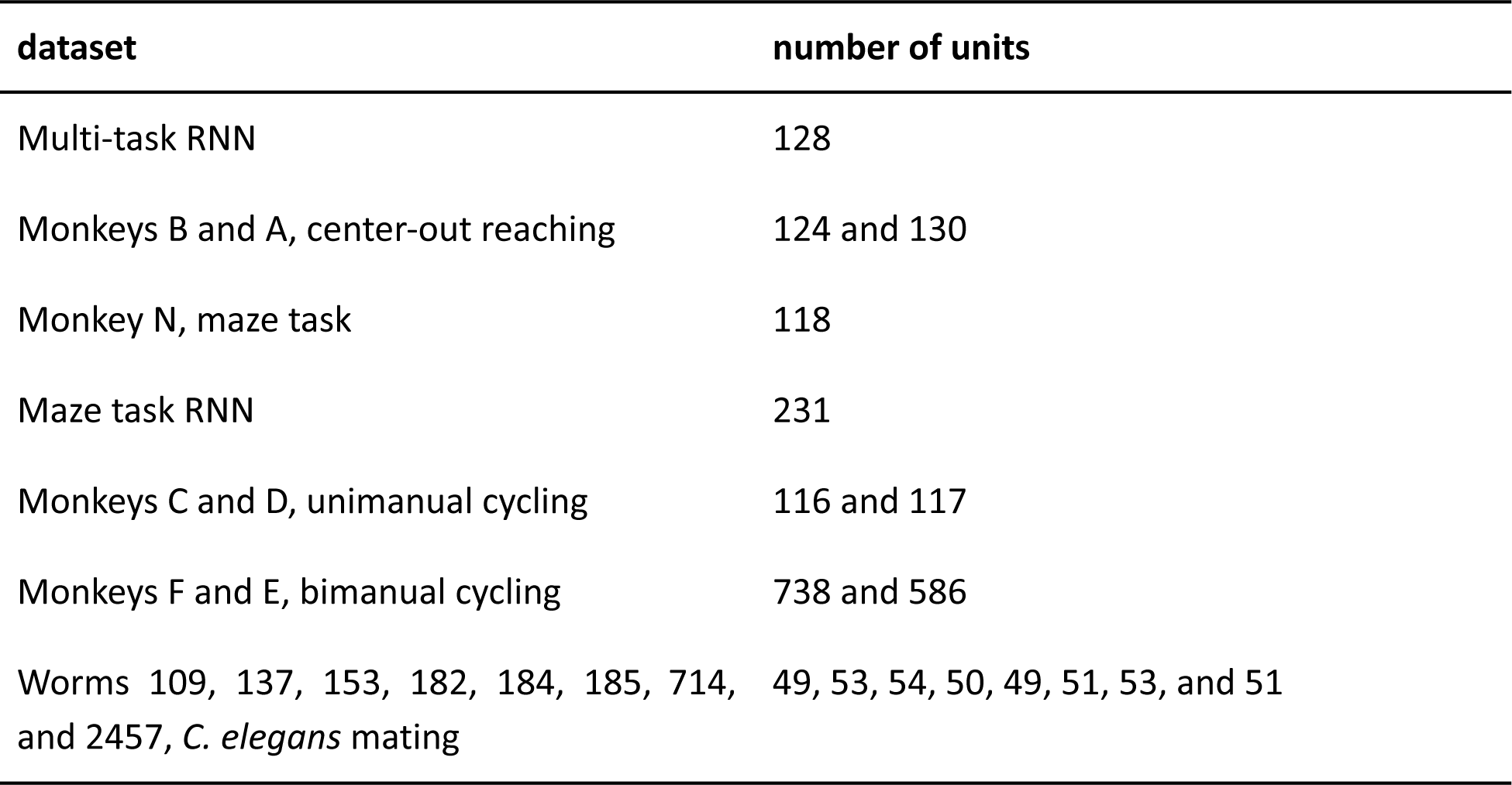

### Center-out reaching

The animal protocols relating to the experiments involving rhesus macaques were approved by the Columbia University Institutional Animal Care and Use Committee. Single neuron and well-isolated multi-unit activity was collected from motor cortex (dorsal premotor cortex and primary motor cortex) while monkeys A and B performed center-out reaches. All recordings were performed in the hemisphere contralateral to the performing arm. Animals reached in eight directions during two contexts: cue-initiated and quasi-automatic. In the cue-initiated context, the animals performed standard center-out reaches with a random, unpredictable delay between 0-1000 ms. In the quasi-automatic context, the delay period was again 0-1000 ms long, and the peripheral target began to move outward from the center of the screen once the go-cue was delivered. In both conditions, the peripheral target needed to be held for 600 ms, after which reward was delivered. Following reward delivery, the monkeys reached back to the central touch-point to begin the next trial.

Due to extensive training, the monkeys’ behavior during cue-initiated and quasi-automatic conditions is extremely similar. Additionally, activity in motor cortex is also extremely similar^27^. We therefore combined cue-initiated and quasi-automatic trials when creating trial-averaged PSTHs. When creating PSTHs, we excluded failed trials and trials where the monkeys’ behavior deviated from normal. This included trials where the outward reaction time was greater than 500 ms, the monkey’s kinematics were substantially different from the median kinematics, or the monkey’s hand moved during the dwell period between outward and return reaches. In practice, we primarily excluded trials because of behavior during the return reach, where behavior was less constrained by task requirements. All analyses, excepting Figure S11, only included data from conditions with a long (i.e., 300 ms) delay.

To construct trial-averaged PSTHs, we spliced neural activity from around target onset, outward reach onset, and return reach onset together to form a single trace that was continuous in time. For each unit, we first aligned spikes from a single trial, within a window of time, to the relevant task events. We used windows of –200 to 350 ms, -225 to 1000 ms, and -225 to 1000 ms for peri-target, peri-reach, and peri-return activity, respectively. We then concatenated each peri-event spike train, convolved each with a 25 ms Gaussian filter, and averaged across trials. Prior to performing SCA or PCA, we performed two additional preprocessing steps. First, we ‘soft-normalized’ the activity of each neuron; each neuron’s mean rate (across times and conditions) was normalized by the range of the rate (across times and conditions) + 5. We standardly use soft-normalization (e.g.,^15,30,54^) to balance the desire for SCA or PCA to explain the responses of all neurons with the desire that weak responses not contribute on an equal footing with robust responses. As a second preprocessing step, the responses for each neuron were mean-centered at each time. We calculated the mean activity across all conditions of each neuron at each time point, and subtracted this mean activity from each condition’s response. This step ensures that dimensionality reduction focuses on dimensions where responses are selective across conditions, rather than dimensions where activity varies in a similar fashion across all conditions^75^.

### Unimanual cycling

Monkeys C and D performed unimanual cycling movements to traverse a virtual track^54^. All recordings were performed in motor cortex, contralateral to the performing arm. Here, only well-isolated single neurons were included in the analyses.

Each trial began with the monkey acquiring an initial target. A final target was then shown in the distance, cueing the monkey as to the number of required cycles. After a variable delay period (500-1000 ms), a go-cue was delivered. After acquiring the final target, the monkey needed to remain stationary for 1500 ms, after which they received a reward. Animals performed 20 unique conditions, each requiring 0.5, 1, 2, 4, or 7 cycles, forwards or backwards movements of the arm, starting at either the top or the bottom of a cycle.

To create trial-averaged PSTHs, we used a previously described time-warping procedure^54^. While behavior during the reaching task was brief, movements during the cycling task could unfold over multiple seconds, necessitating a more detailed alignment procedure. Briefly, the time-base for each individual trial was scaled such that the virtual position for that trial closely matched the mean position across all trials of a given condition. This procedure aligned neural activity (and behavior) across trials of a given condition. We then took a subsequent step (not used in^54^) to better align data across conditions; we applied a second scaling to the time-base calculated for each condition, such that the duration of each individual cycle was 500 ms (0.5 cycle conditions were not adjusted). Because both animals tended to cycle at approximately 2 Hz, this second adjustment was fairly minor. Prior to performing SCA, responses were soft-normalized, as described above.

### Bimanual cycling

Monkeys E and F performed a cycling task similar to that described above; the animals performed cycling movements to traverse a virtual track^38^. While monkeys C and D performed only unimanual cycling movements, monkeys E and F performed both unimanual and bimanual conditions. As above, each condition required the monkey to move between two virtual targets, with a given condition requiring either forwards or backwards arm movements, beginning at either the top or bottom of a cycle. Unlike the task performed by monkeys C and D, each condition was seven cycles long and required movements of either the left, right, or both arms.

For unimanual conditions, only one arm (the ‘performing arm’) was allowed to move; a trial was aborted if the ‘non-performing arm’ moved outside of a small window centered around the bottom of the cycle (±0.05 and ±0.07 cycles for monkey E and F, respectively). During bimanual conditions, the animal was required to move both arms, maintaining either a 0-degree (bimanual0 conditions) or 180-degree (bimanual180 conditions) phase offset between the two hands. Virtual position was determined by the average position of the two arms. Critically, the pedals were not ‘locked’ together; the monkeys needed to actively maintain a particular phase offset. If the angle between the two hands exceeded 180 degrees from the desired offset (i.e., if one arm became more than one half-cycle ahead of the other) the trial was aborted. While the position of the pedals were not yoked, we did provide a weak restorative force to help the animals maintain the instructed phase offset.

Recordings were made in the primary and premotor cortices of both hemispheres and included both single neurons and well-isolated multi-units. To generate trial-averaged PSTHs, we aligned single-trial spikes to movement onset and convolved each spike-train with a 25 ms Gaussian. We then used the same alignment procedure described previously^38^. This procedure results in an adjusted time-base in which each individual cycle, for each condition, is 500 ms in duration. As was true for monkeys C and D, monkeys E and F cycled at approximately 2 Hz for all conditions, so the warping procedure provided only a small adjustment. All analyses of the bimanual cycling dataset only included data from the middle cycles (cycles 2-6). Prior to performing SCA, responses were soft-normalized, as described above.

### *C. elegans* mating

Here, we analyzed calcium imaging data recorded from the posterior brain of eight male worms during mating^68^. All worms co-expressed GCaMP6s and mNeptune in all neuronal nuclei. Individual neurons were identified via their location, morphology, and expression of specific fluorescent markers, and the occurrence of specific behavioral motifs was detected automatically using various kinematic measures (e.g., swimming velocity, tail velocity) as well as the orientation and position of the male worms relative to the hermaphrodites. All analyses involved single-trial fluorescence signals; for details of the preprocessing steps see^68^. Each neuron’s response was fully normalized by its dynamic range prior to performing SCA or PCA.

### Multitask RNN

The multitask network^13^ consisted of 128 recurrently connected units and received three inputs: fixation (1-dimensional), stimulus (4-dimensional), and context (15-dimensional). The fixation input served both as a ‘hold’ signal and a ‘go cue’, depending on the context. The set of stimuli contained two separate two-dimensional vectors composed of *Asin*(*θ*) and *Acos*(*θ*), where each vector encoded a different one-dimensional circular variable (*θ*_1_, *θ*_2_) scaled by an amplitude *A*. The context input indicates the current task on any given trial. Context input was encoded in a one-hot vector where the index associated with the current task was one and all other indices were zero. The network generated two outputs: a fixation output (1-dimensional) and a response angle (2-dimensional, the sine and cosine of the response angle). During training, the duration of the stimulus epochs (and memory epochs, when present) was determined by a random draw from a uniform distribution to prevent the network from predicting task period transitions.

The RNN was defined by standard functions:

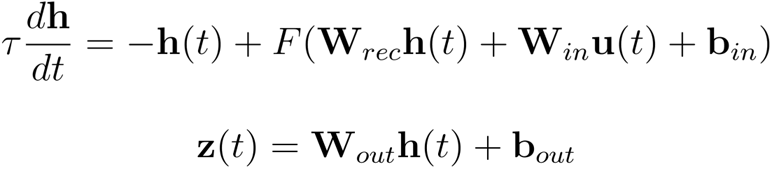

where *F* is a tanh activation function, **h** is a vector of network activations, **u** are the inputs, **b** are offset terms, and **z** is the output vector. **W***_rec_*, **W***_in_*, and **W***_out_* are the recurrent, input, and output weight matrices, respectively. The network was trained with standard L2 penalties on the weights and rates. Additionally, private neuron noise and input noise were added during training. The weights were trained via backpropagation through time using the Adam optimizer^94^. During training, the network performed all tasks in an interleaved order.

Below, we summarize the 15 tasks performed by the network.

Delayed Response: Delay-pro: Move in the same direction as stimulus (*φ_response_* = *θ_stimulus_*) after a delay. Delayed-anti: Move in opposite direction as stimulus (*φ_response_* = *θ_stimulus_* + π) after a delay. Stimulus remains on throughout stimulus and response periods.

Memory Response: Same as above, except the stimulus disappears during the memory and response period.

Reaction Timed: Same as Delayed Response, but a response is required as soon as the stimulus is delivered.

Compare: Move in the direction of the stimulus with the largest amplitude. Compare-stim1: Compare stimuli delivered via stimulus modality 1. Compare-stim2: Compare stimuli delivered via stimulus modality 2.

Context Compare: Same as above, except stimuli are delivered using both modalities. ContextCompare-stim1: compare stimuli delivered via modality 1, ignore modality 2. ContextCompare-stim2: compare stimuli delivered via modality 2, ignore modality 1. ContextCompare-both: attend both modalities.

Category Match: Network is delivered two sequential stimuli (using the same stimulus modality). Category-pro: immediately respond in the direction of the second stimulus if the two stimuli belong to the same category (*θ_stim1_*, *θ_stim2_* < π or *θ_stim1_*, *θ_stim2_* > π). Category-anti: immediately respond in the direction of the second stimuli if the two stimuli belong to opposite categories.

MatchToSample: Network is delivered two sequential stimuli (using the same stimulus modality). MatchToSample-match: immediately respond in the direction of the second stimulus if the two stimuli are the same (*θ_stim1_* = *θ_stim2_*). MatchToSample-nonmatch: immediately respond in the direction of the second stimulus if the two stimuli are 180° apart (*θ_stim1_* = *θ_stim2_* + π)

### SCA hyperparameters

As demonstrated in Figure S3, SCA produced highly similar results across a wide range of hyperparameter choices. Below we list the specific hyperparameters used for each dataset. ‘Default’, and rationale for non-default values is described above.

| dataset | $\lambda_{sparse}$ | $\lambda_{orth}$ | number of factors |
| --- | --- | --- | --- |
| Multi-task RNN | default | default | 25 |
| center-out reaching | default | default | 8, 16, or 24 |
| maze task | default | default | 12 |
| Maze task RNN | default | default | 12 |
| unimanual cycling | 0.03 (~10x default) | default | 40 |
| bimanual cycling | 0.01 (~10x default) | default | 50 |
| *C. elegans* mating | 0.05 (~10x default) | default | 15 |

### Reaching analyses

#### Occupancy and occupancy concentration

For the reaching datasets, we computed the occupancy (variance across conditions) of each dimension, *k*, at each timepoint, *t*, where *t* began with target onset and ended 300 ms after the onset of the return reach:

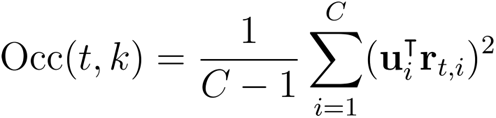

where *C* is the number of conditions, **u** is a vector of PCA or SCA loadings, and **r** is a vector of firing rates. For each dimension, we calculated the fraction of the total occupancy accounted for by the preparatory, execution, or postural epochs (e.g., Fig. 2D, *left*). For example, the fractional occupancy accounted for by the preparatory epoch is defined as:

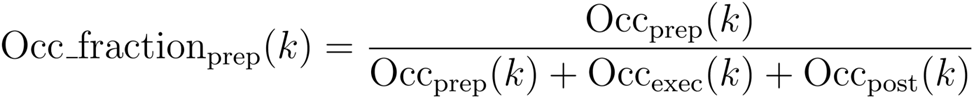

Where Occ_prep_,Occ_exec_, and Occ_post_ are the sum of the occupancies during the preparatory, execution, and posture epochs. These three epochs are defined below.

| epoch | time window(s) |
| --- | --- |
| preparatory | $T_{target} : T_{reach},$<br>$T_{return} - 300 : T_{return}$ |
| execution | $T_{reach} : T_{reach} + 300,$<br>$T_{return} : T_{return} + 300$ |
| posture | $T_{reach} + 300 : T_{return} - 300$ |

*T_target_*, *T_reach_*, and *T_return_* are the times of target onset, outward reach onset, and return reach onset, and the time ranges are defined in milliseconds.

To summarize the distribution of occupancy within a single dimension, we calculated ‘occupancy concentration’: the sum of the absolute difference between all fractional occupancy pairs:

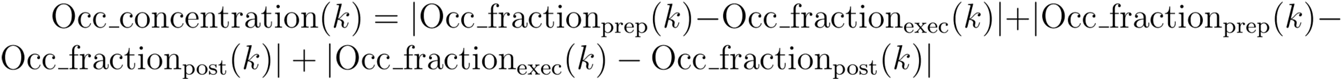

The maximum possible occupancy concentration for a single dimension would be 2 (e.g., fractional occupancies of 1,0, and 0 for the preparatory, execution, and posture epochs, respectively). To summarize occupancy concentration across all dimensions, we simply calculated the median concentration across all dimensions (Fig. 2D, *right*).

#### Supervised dimensionality reduction

To compare SCA and PCA latents to those recovered by a supervised method, we used the dimensionality reduction method documented in^26^(Fig. 2E). This method identifies orthogonal sets of dimensions that capture a large fraction of variance during user-defined epochs:

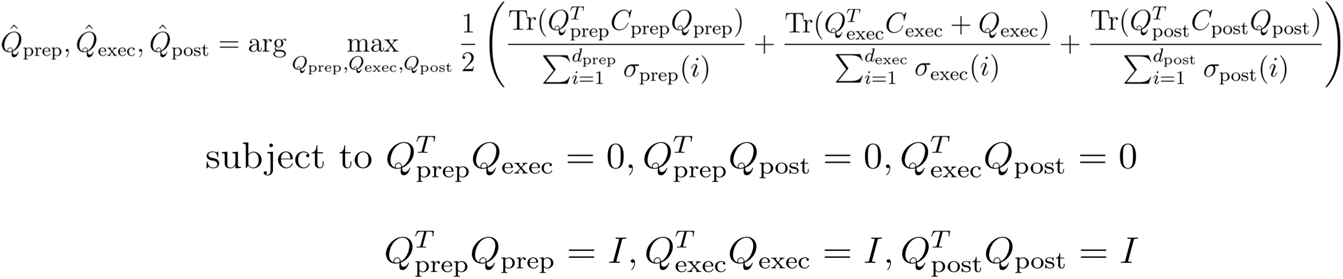

Where *Q* are the loadings for the preparatory, execution, and posture-related spaces, *C* are the covariance matrices of neural activity during preparatory, execution, and posture-related epochs, and σ(*i*) is the *i^th^* singular value of *C*.

#### Bi-cross-validation

As a stringent test of whether SCA recovers structure that exists across conditions during reaching, we performed a bi-cross-validation analysis, which required generalization to both held-out conditions and held-out neurons. We first fit SCA using data from all conditions and a set of training neurons (90% of all neurons). We then used linear regression to fit the loadings between test-neurons and factors, using activity from 7 of 8 conditions. Finally, we use factor activity from the held-out condition and the loadings recovered via regression to predict the activity of the held-out neurons during the held-out condition. We repeated this process 80 times (10-fold validation for the neurons, 8-fold validation for the conditions).

### Bimanual cycling analyses

#### Validating existence of a single speed dimension during bimanual cycling

Training SCA on bimanual cycling activity from monkey E yielded a single dimension that was highly correlated with the monkey’s average cycling speed over a cycle (Figure S15E). To validate this result, we used linear regression to identify a single speed dimension from the neural activity from one condition. We then asked whether activity in this dimension was correlated with cycling speed during a second condition. More specifically, we used linear regression to solve for **b**:

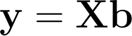

Where **y** is a vector corresponding to the angular cycling speed for a single condition after low-pass filtering (third-order Butterworth, cutoff frequency: 0.8 Hz) and **X** is a ***TxN*** matrix of trial-averaged firing rates for one bottom-start condition. To prevent overfitting, we used ridge-regression with *λ* = 0.01. After calculating **b**, we projected neural activity from all top-start conditions into the same dimension and calculated the *R*^2^ between the projection and low-pass filtered angular speed for each condition.

### C. elegans analyses

#### Correlating factors with behavior

The magnitude of individual SCA factors often correlated with the onset of specific behavioral motifs (e.g., Fig. 6C). To quantify this relationship (Fig. 6D), we generated boolean vectors that corresponded to when each individual behavioral motif occurred for each worm. We convolved each vector with a gamma distribution (shape parameter: 2, scale parameter: 5) to produce a continuous behavioral trace, then calculated the correlation between these behavioral traces and each SCA (or PCA) factor.

#### Generating pseudo-factors

SCA often recovered qualitatively similar loading vectors for the same behavioral motif in different worms (e.g., Fig. 7B). We quantified this relationship in two ways. First, we measured the angle between loading vectors (Figure S18). Second, we generated ‘pseudo-factors’ from the loadings of one worm and the neural activity of a second. If SCA is recovering truly similar loading vectors for different worms, then the magnitude of the pseudo-factor should correlate with the behavior of the worm that provided the neural activity.

For Figure 7C,D, we generated pseudo-factors using the loading vectors related to factors with high correlations to behavior. Specifically, we identified the loading vectors associated with the highest 25% of these correlations. We then generated pseudo-factors for each second worm (i.e., the worm other than the one that provided the loading vector) that exhibited the relevant behavioral motif. The critical result – that pseudo-factors generated from SCA loadings were more correlated than those generated from PCA loadings – held if we constructed pseudo-factors from all of the original loading vectors (mean correlation(standard error): 0.34(0.009) vs. 0.29(0.008), p < 0.001, SCA vs. PCA, rank sum test).

### Software Availability

A code package for SCA is available at https://github.com/glaserlab/sca.

---

**Figure S1.**
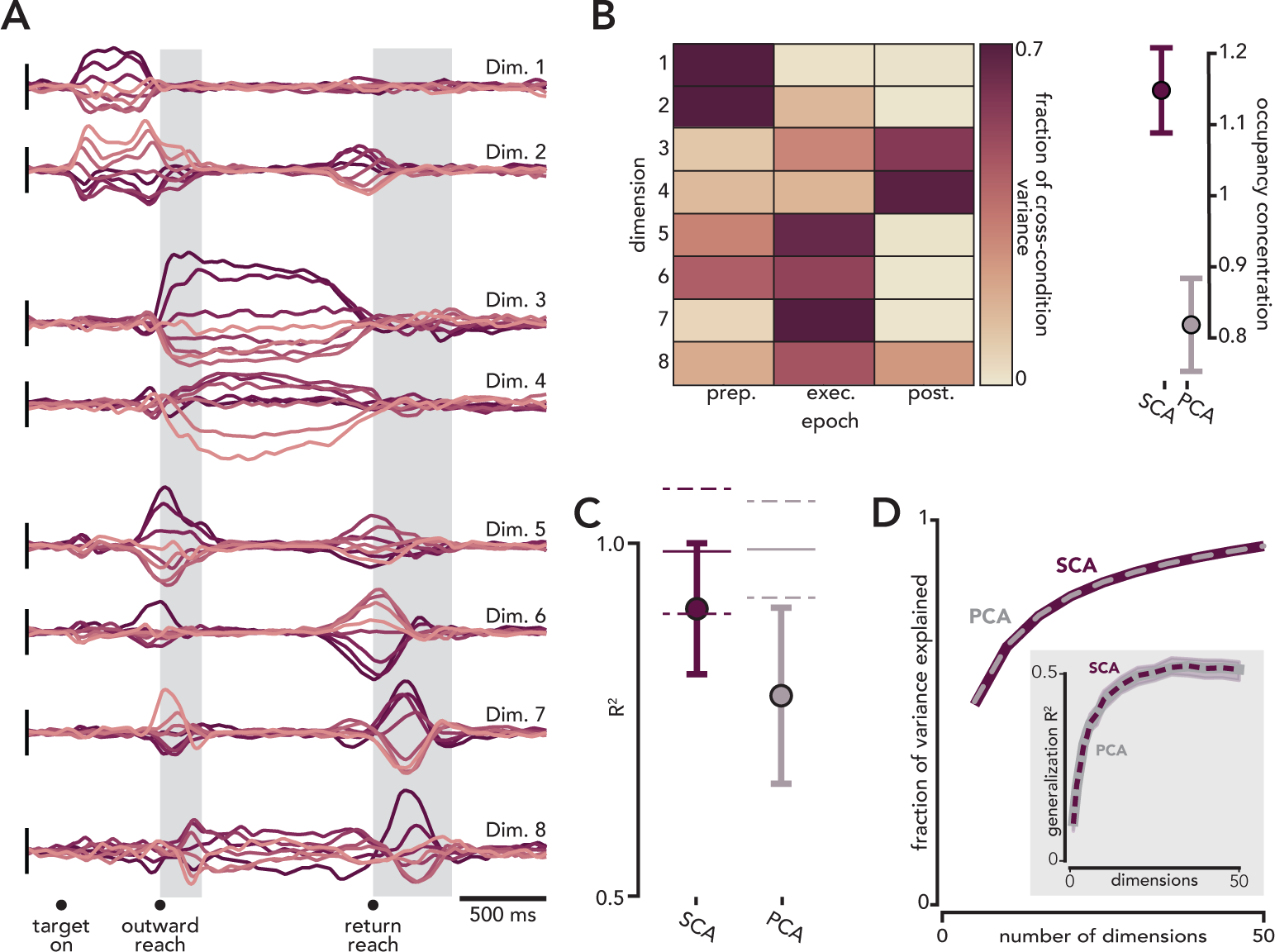
SCA applied to motor cortical reaching data from a second monkey. Same format as Figures 2,3. **A.** Eight SCA dimensions **B.** Left: fraction of occupancy in each SCA dimension across three task epochs. Right: Occupancy concentration (mean and standard deviation, calculated across redrawn populations, p < 0.01, n=100 populations). **C.** Reconstructing individual SCA factors using activity in preparatory, execution-related, or posture-related factors found via a supervised method. The reconstruction of factors from a single set of supervised factors was significantly higher for SCA than PCA (p < 0.01, *bootstrap test*), and the difference in R^2^ between predicting factors using a single set of supervised factors vs. all supervised factors was significantly smaller for SCA than PCA (p < 0.01, *bootstrap test*). **D**. Cumulative fraction of variance explained. Inset. Reconstructing activity of held-out neurons during held-out conditions from SCA or PCA factors.

**Figure S2.**
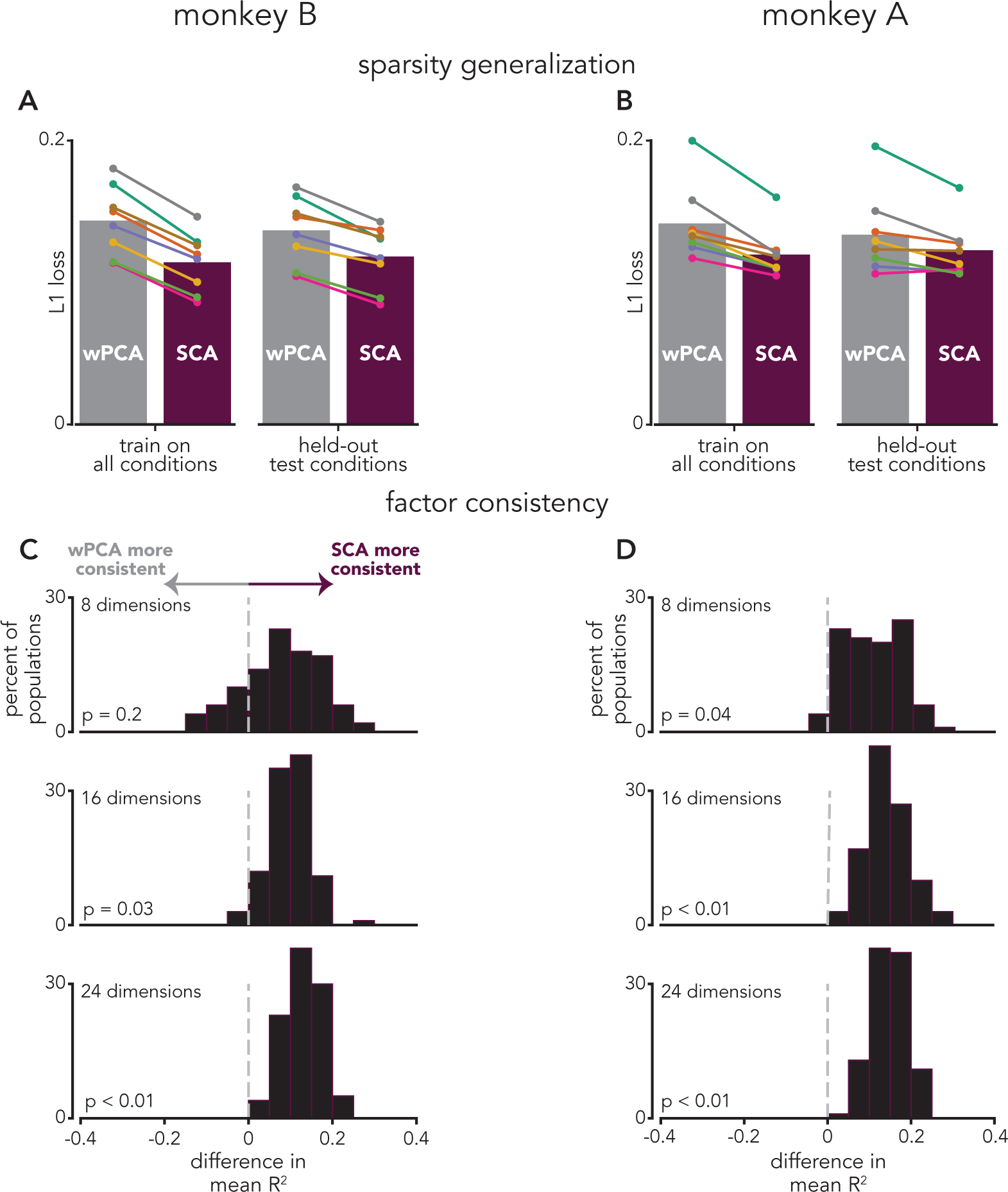
SCA sparsity generalizes to held-out conditions and SCA factors are more consistent than PCA factors. **A.** Testing generalization of sparsity. Left: wPCA and SCA were performed on all conditions (8 wPCA and SCA factors). *Colored circles* indicate L1 loss for each condition. As expected, SCA factors were significantly more sparse than wPCA factors (p = 0.006, *sign-rank test*). Right: wPCA and SCA were performed on seven of the eight conditions, and the L1 loss was calculated for the held-out condition; here too, the SCA factors were significantly more sparse than the wPCA factors (p = 0.006, *sign-rank test*). **B.** Same as **A**, monkey A (p = 0.006 and p = 0.013, wPCA L1 loss vs. SCA L1 loss, left and right, respectively). **C.** SCA factors are more consistent across resampled populations than PCA factors. Bootstrapped populations were constructed by resampling single units from the original population. SCA (or PCA) was performed on the bootstrapped population and the maximum correlation between each bootstrapped factor and all original (i.e., non-bootstrapped) factors were calculated. The average maximum R^2^ was then calculated for PCA and SCA. This process was repeated 100 times, and significance was determined by counting the number of bootstrapped populations that produced a higher mean R^2^ value for PCA than SCA. **D.** Same as **C**, but for monkey A.

**Figure S3.**
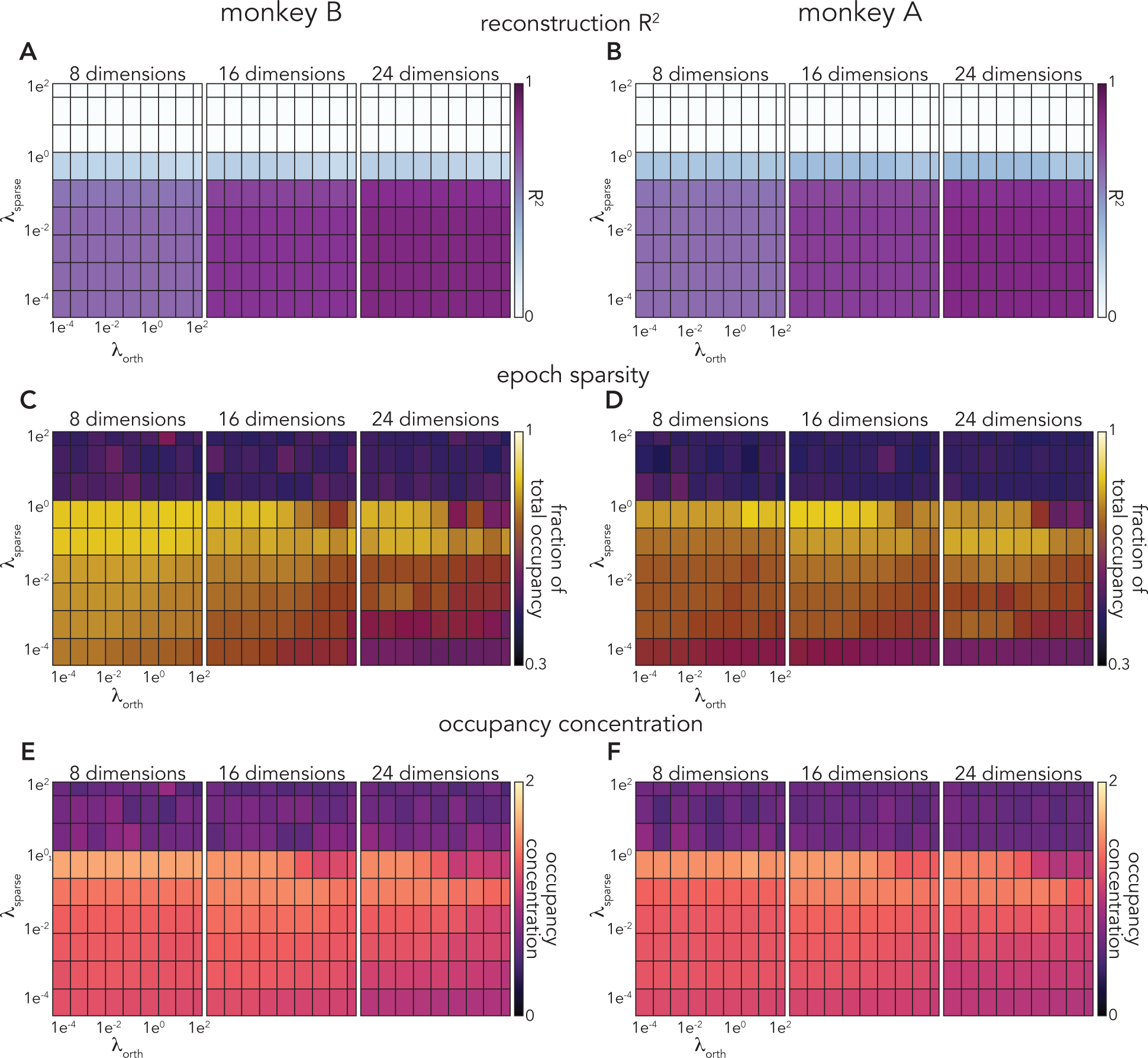
SCA performs similarly across a wide range of hyperparameters. To quantify the effects of changing λ_orth_ and λ_sparse_, we swept these parameters across six orders of magnitude. We found a relatively minor quantitative differences across the entire range of λ_orth_values. Increasing λ_sparse_ had similarly small quantitative effects across roughly four orders of magnitude. However, above some threshold value ( λ_sparse_≈ 0.1, *white rectangles*, in A and B), SCA prioritizes sparsity over minimizing reconstruction error, and the magnitude of the recovered factors tends towards 0. **A,B.** R^2^ between trial-averaged neural activity and SCA reconstruction, monkeys B and A, respectively. **C,D.** For each SCA dimension, we first calculated the maximum fraction of the total occupancy accounted for within a single epoch (preparatory, execution, or posture). We then took the median (across all SCA dimensions) of the maximum fractional occupancy values. **C** and **D** correspond to monkey B and A, respectively. **E,F.** Occupancy concentration (calculated as outlined in *Methods*), monkeys B and A, respectively.

**Figure S4.**
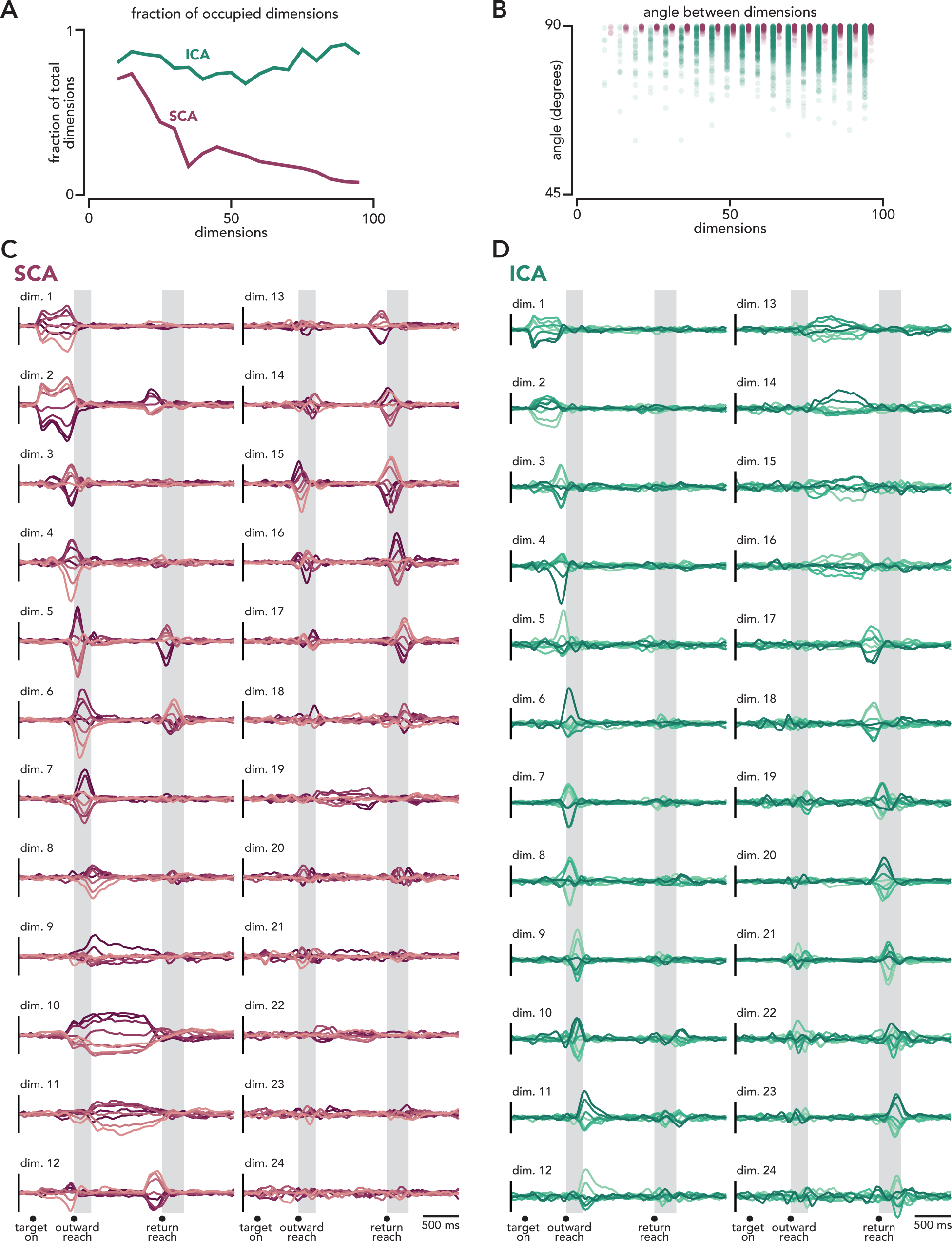
SCA is less likely to generate ‘overly-sparse’ factors than ICA. **A.** Fraction of dimensions with appreciable occupancy as function of total dimensions. A dimension was considered ‘appreciably occupied’ if the sum of the cross-condition variance within that dimension was greater than 30% than that of the dimension with the largest cross-condition variance. **B.** Angle between all individual dimensions. As a larger number of dimensions are requested, both SCA (*maroon*) and ICA (*green*) tend to find dimensions that are more aligned. However, this effect is much more dramatic for ICA. Note that even for a moderate number of dimensions (e.g., less than 20), ICA dimensions are more aligned than SCA dimensions. C. Projections in twenty-four SCA dimensions. These SCA dimensions share many of the same features as those plotted in Figure 3, namely dimensions that are primarily occupied during preparatory, execution, or posture epochs. In the set of 24 SCA dimensions however, there are more dimensions that are only occupied prior to (or during) outward or return reaches. Note the number of dimensions that are largely unoccupied (e.g., dimensions 21-24). **D.** Projections in twenty-four ICA dimensions. Nearly all dimensions have substantial activity and are only occupied prior to/during outward or return reaches.

**Figure S5.**
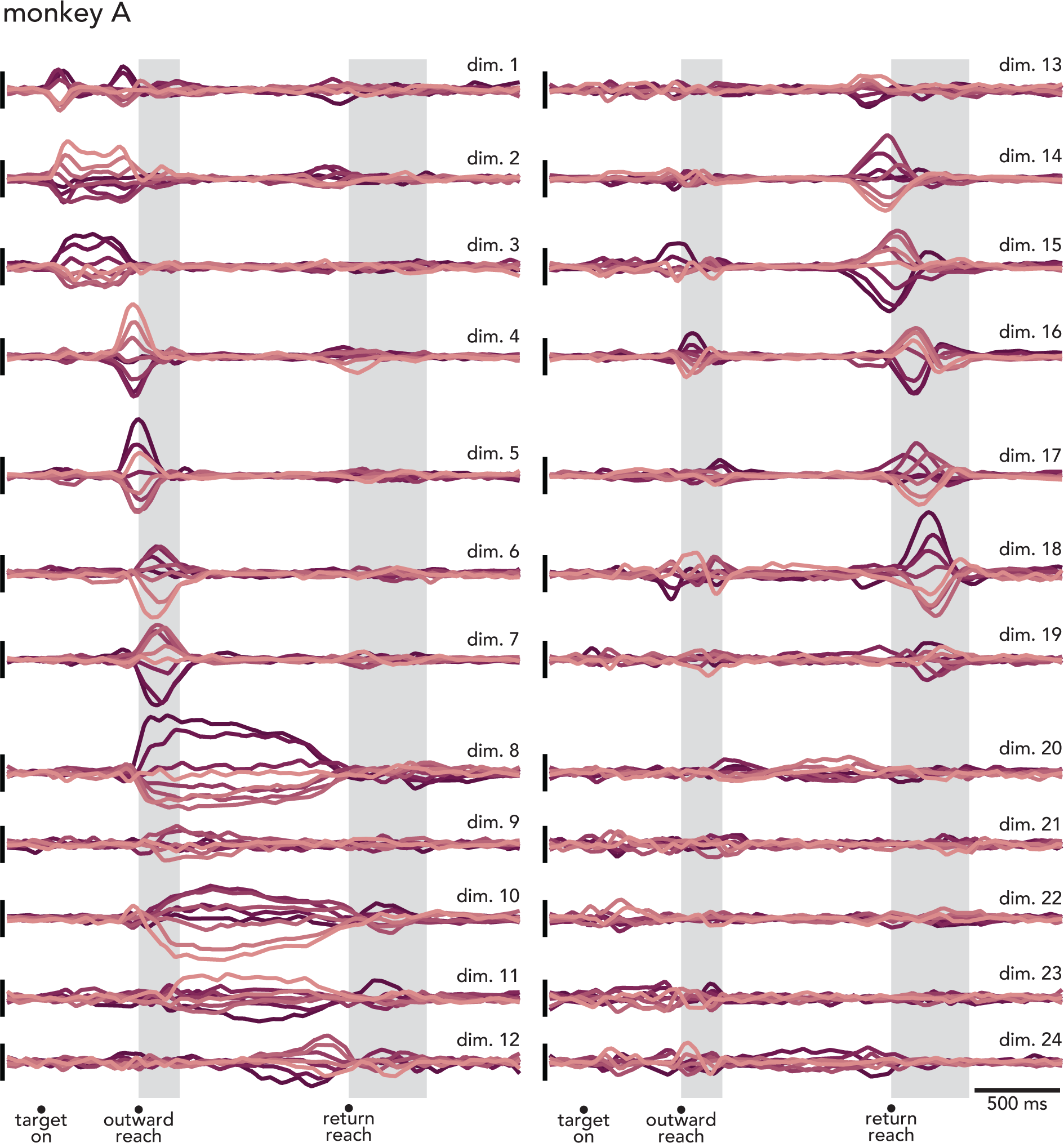
Twenty-four SCA dimensions, motor cortical reaching data, monkey. **A.** Unlike monkey B, the SCA dimensions found for monkey A tended to be occupied prior to or during outward or return reaches, but not both. This difference is due to the greater dissimilarity between outward and return reaches for monkey A. For monkey A, return reaches were roughly half as fast as outward reaches (the mean peak speed for return reaches was 49% that of outward reaches). Relatedly, the alignment index for outward and return reaches was 0.29. For monkey B, the alignment index for outward and return reaches was 0.43.

**Figure S6.**
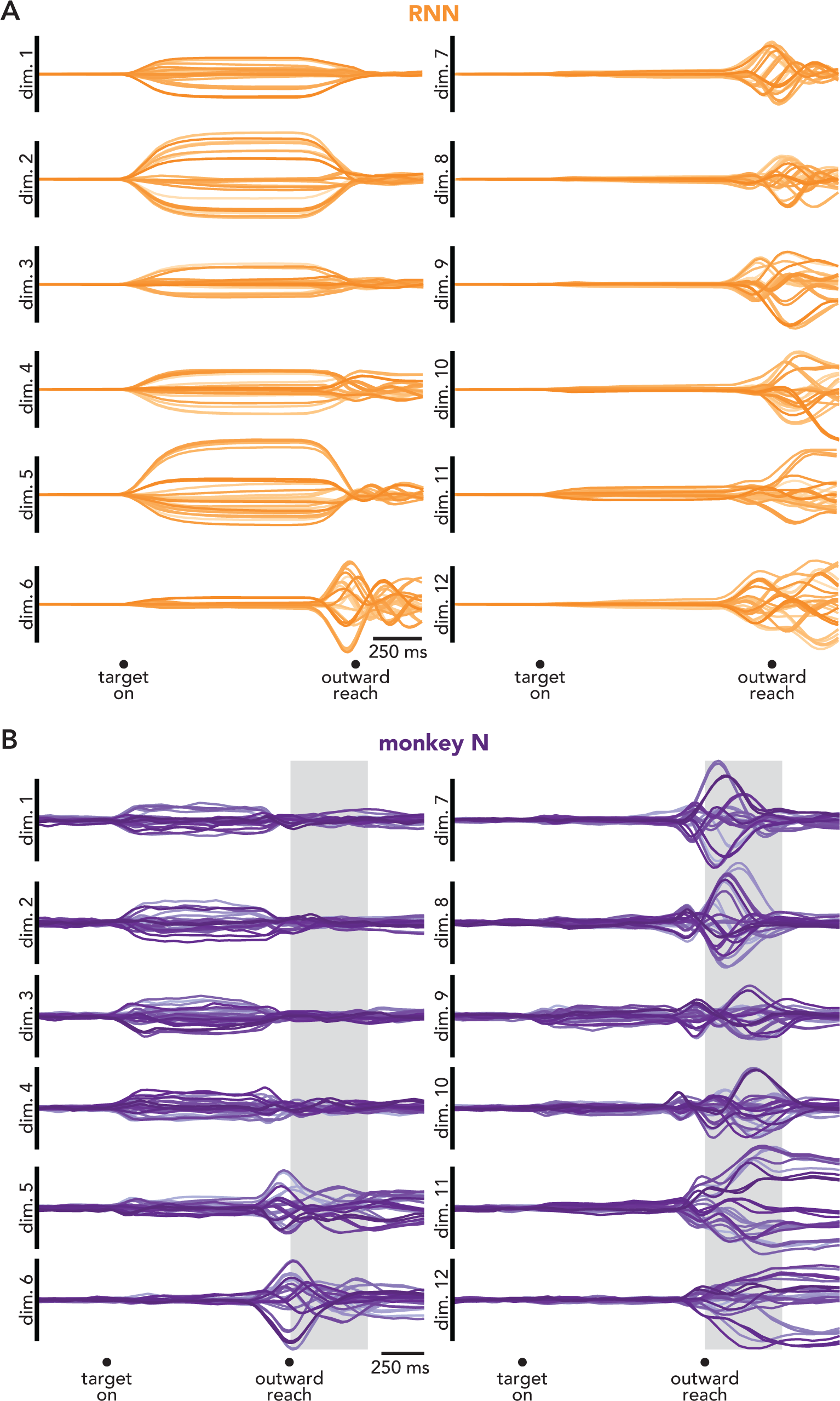
SCA can capture factors reflecting quasi-oscillatory dynamics. An obvious concern regarding SCA is that it might ‘over-sparsify’ data, separating activity into more dimensions than necessary. This concern interacts with present uncertainty regarding the nature of rotational dynamics in motor cortex during reaching. Neural network models of reach generation reproduce the empirical rotations using nonlinear dynamics whose linear approximation can be either quasi-oscillatory (Sussillo et al. 2015) or non-normal (Hennequin et al. 2014). With oscillatory dynamics, neural trajectories exit dimension one, rotate into dimension two, then subsequently rotate back into dimension one (typically decaying as they do so). With non-normal dynamics, neural trajectories exit dimension one, rotate into dimension two, and then rotate into a third dimension without returning to dimension one (Murphy and Miller 2009; Goldman 2009). Networks can blend both strategies (Zimnik and Churchland 2021) and the distinction can become blurry for nonlinear dynamics (e.g., time-varying eigenvectors can cause oscillatory dynamics to exhibit non-normal-like features). Yet the solutions are different enough that it is worth distinguishing between them when possible. The data in Figure 2E, analyzed via SCA, are overall most consistent with non-normal dynamics. There are some biphasic aspects to responses, suggesting a modest oscillatory component, but overall activity tends to be monophasic and to rotate from some dimensions (e.g., dimension 6) into others (e.g., dimension 8) without returning. A key question is whether this feature – largely monophasic factors – simply appears because SCA optimizes for sparsity and monophasic factors are more sparse than multiphasic factors. Here we address that question in two ways. First, we analyzed population activity from a recurrent neural network (RNN) that is known to use a quasi-oscillatory solution. Second, we re-analyzed population activity that had previously been identified as having a sizable quasi-oscillatory component. If SCA works as desired – i.e., if it does not over-sparsify data – then it should return multiphasic factors in both cases. This was indeed the case. **A.** Twelve SCA dimensions, identified from the activity of an RNN (Sussillo et al. 2015) trained to generate empirical patterns of muscle activity, recorded from a monkey performing a task that required both curved and straight reaches (i.e, the ‘maze’ task (Churchland et al. 2012)). Each *orange trace* represents one condition. Previous work (Sussillo et al. 2015) has thoroughly characterized the dynamics of this network. During reach execution, across all conditions, the network exhibits a single fixed point. The linearized dynamics, local to this fixed point, involve a small number of oscillatory modes. In agreement with this ground truth, SCA identifies multiphasic factors. Also in agreement with a quasi-oscillatory (rather than non-normal) solution, activity tends not to leave a dimension once it enters it. Most dimensions that are active just before movement onset (dimensions 6-12) remain active >250 ms after movement onset. This argues that the mostly monophasic factors in Figure 2E are not a distortion; if the empirical dynamics had been strongly quasi-oscillatory, SCA would have found more strongly multiphasic factors as it does for the network model shown here. For the data in Figure 2E, a better approximation is likely to be non-normal dynamics. Alternatively, oscillatory dynamics may involve eigenvectors that change swiftly as the overall location in state-space changes (Kaufman et al. 2016). **B.** Twelve SCA dimensions identified in motor cortex activity from a monkey performing the maze task. Each *purple trace* represents a condition. Data are the same as those analyzed in Figure 3 and Supplementary Movie 3 of (Churchland et al. 2012). For this monkey, the factors identified by SCA resemble those when analyzing the network’s population response. Many individual factors are multiphasic and are active across fairly long durations. This is in contrast to the data in Figure 2E (from monkey B) where execution-active factors are mostly monophasic and short-lived. The fact that SCA identified multiphasic factors for monkey N demonstrates that SCA recover this structure when it exists in the data. These results also suggest the two monkeys (N and B) use modestly different internal solutions despite performing similar tasks. The similarities and differences of solutions can be compared by considering what occurs in plots of the sort shown in Figure 2F. In a quasi-oscillatory solution (e.g., monkey N), the set of preparatory states (arranged along the vertical axis) rotates into execution dimensions (gray plane) and then continues to rotate, producing multiphasic factors. In a non-normal solution (e.g., monkey B), the set of preparatory states rotates into execution dimensions (as before). However, as rotations continue, they continuously enter new dimensions and leave old ones, resulting in monophasic factors. Mixed solutions are also possible. Indeed, neither monkey appears to use an entirely pure solution. There is some biphasic activity in the factors of monkey B, and monkey N shows some factors that are primarily active early in execution and others that are primarily active late. Presumably these two solutions lie on a spectrum of ‘good’ solutions that can be used by nonlinear recurrent networks. A potentially related phenomenon was found in the case of the cycling task (Figure S7, S10). Some monkeys used largely distinct factors for forward and backward cycling (e.g., monkey D, Figure S10) and others used a few SCA factors during both conditions (e.g., monkey C, Figure S7). Whether such differences have meaningful computational or behavioral consequences, or whether they simply reflect redundancy in the available solutions, is a reasonable future question.

**Figure S7.**
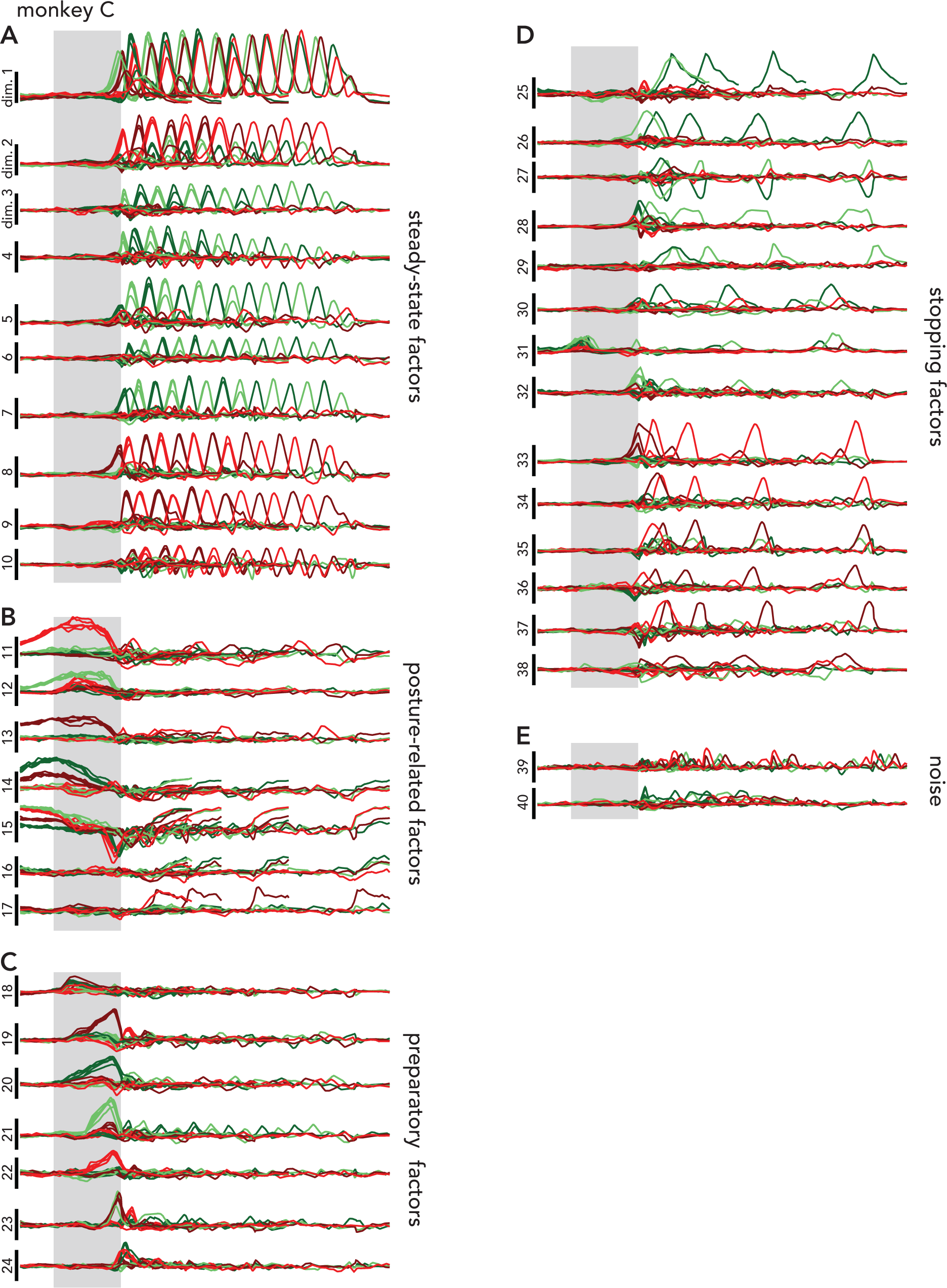
All SCA dimensions identified for monkey C. Dimensions are manually grouped by putative function. **A.** Steady-state factors. **B.** Posture-related factors. **C.** Preparatory factors. **D.** Stopping-related factors **E.** Noise.

**Figure S8.**
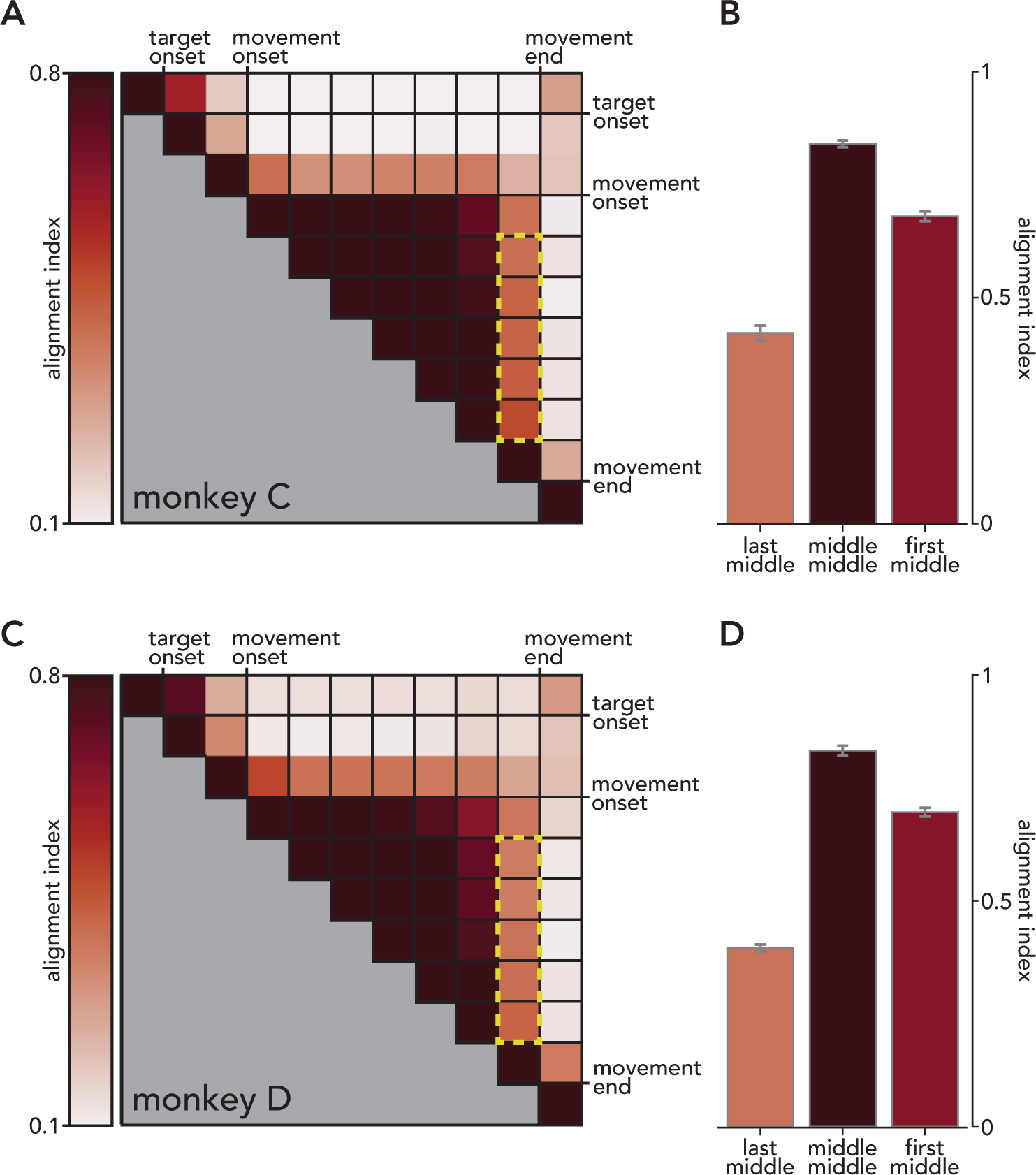
Alignment Indices for motor cortical activity during unimanual cycling. **A.** Pairwise alignment indices for all times during seven-cycle forwards movements. Each square corresponds to 500 ms (the duration of one cycle). The *dashed yellow box* corresponds to the alignment indices between the middle cycles and the final cycle. **B.** Mean alignment index (and standard error) for all pairs of individual cycles within four or seven cycle conditions (i.e., half, one, and two cycle conditions did not have middle cycles). **C,D.** Same as A,B but for monkey D.

**Figure S9.**
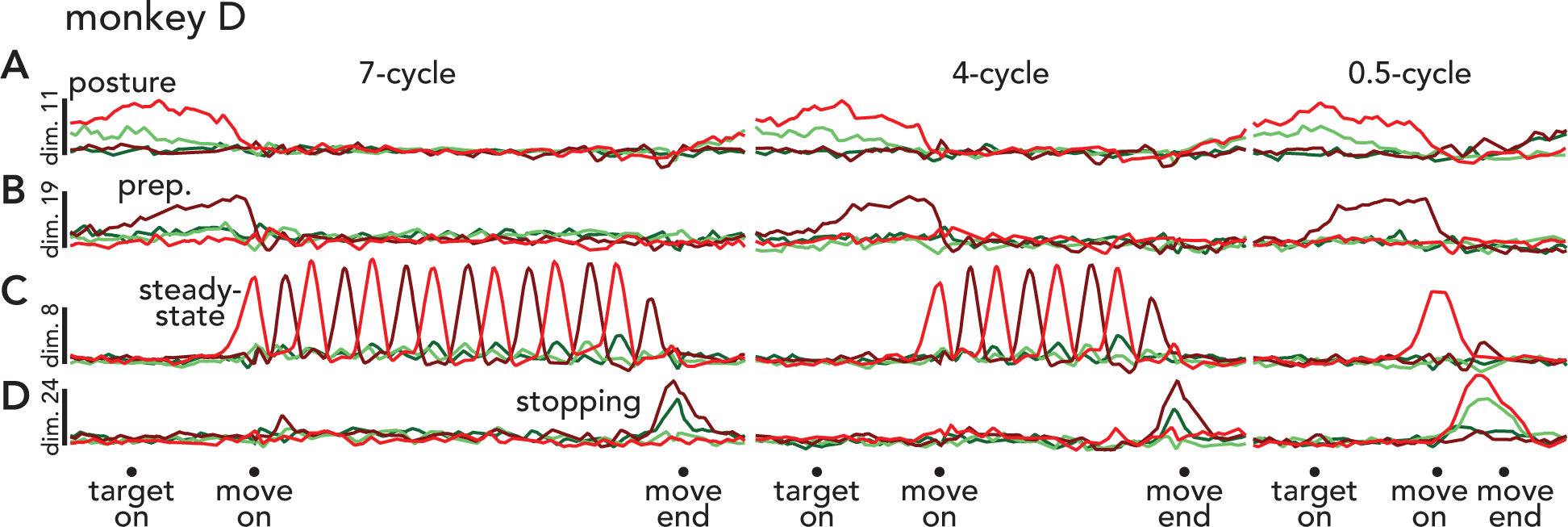
Example SCA dimensions from a second cycling monkey. Same format as Figure 4. **A.** Projections from all seven cycle (left), four cycle (middle), and half cycle (right) conditions in a single posture-related SCA dimension. During seven and four cycle conditions, activity for top-start conditions (*light traces*) is high at the start of the trial and begins to increase after movement ends. However, during half cycle conditions, activity related to bottom start conditions (*dark traces*) begins to increase after movement onset. This is expected, as the monkey’s arm ends at the top of a cycle during bottom start, half cycle conditions. **B.** Projections from all seven, four, and half-cycle conditions in a single preparatory dimension. **C.** Projections in a steady-state dimension. **D.** Projections in a stopping-related dimension.

**Figure S10.**
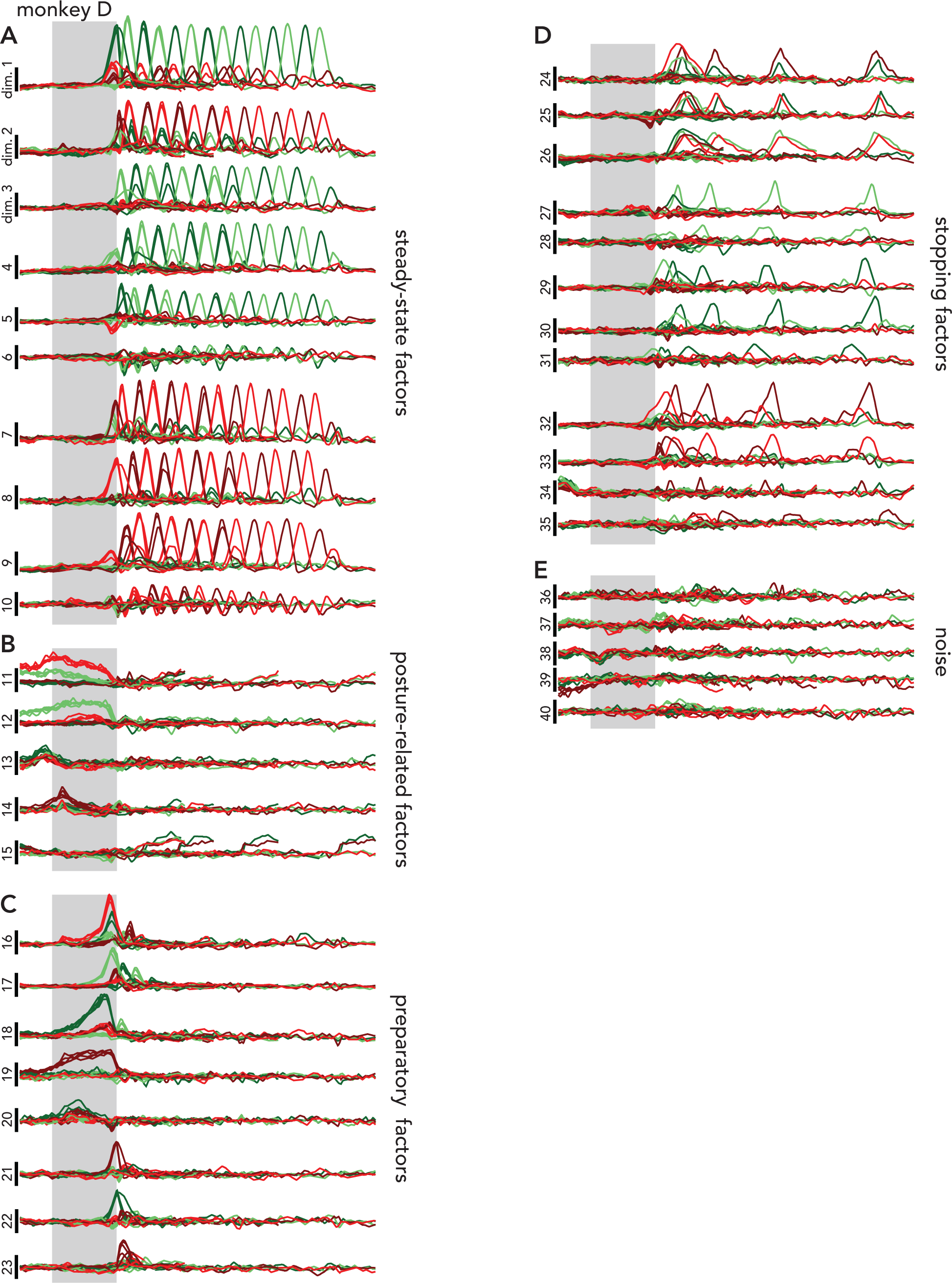
All SCA dimensions identified for monkey D. Dimensions are manually grouped by putative function. **A.** Steady-state factors. **B.** Posture-related factors. **C.** Preparatory factors. **D.** Stopping-related factors **E.** Noise. When comparing the SCA factors for the two unimanual cycling monkeys (monkey C and monkey D), the most notable difference was a higher degree of overlap between forward and backward steady-state activity in monkey C than in monkey D (compare steady-state factors in Figure S7 and Figure S10). In agreement with this difference in steady-state factors, the alignment between forward and backward cycling differed in the two animals. As stated in the main text, forwards and backwards cycling activity occupied partially overlapping spaces for monkey C (alignment index: 0.40±0.03 (mean and standard deviation, calculated across conditions), while the alignment index for these conditions was half as large for monkey D (alignment index: 0.20±0.01). While determining the functional significance of this difference (if any) is beyond the scope of this work, the fact that SCA preserves this structure demonstrates its utility as an exploratory tool.

**Figure S11.**
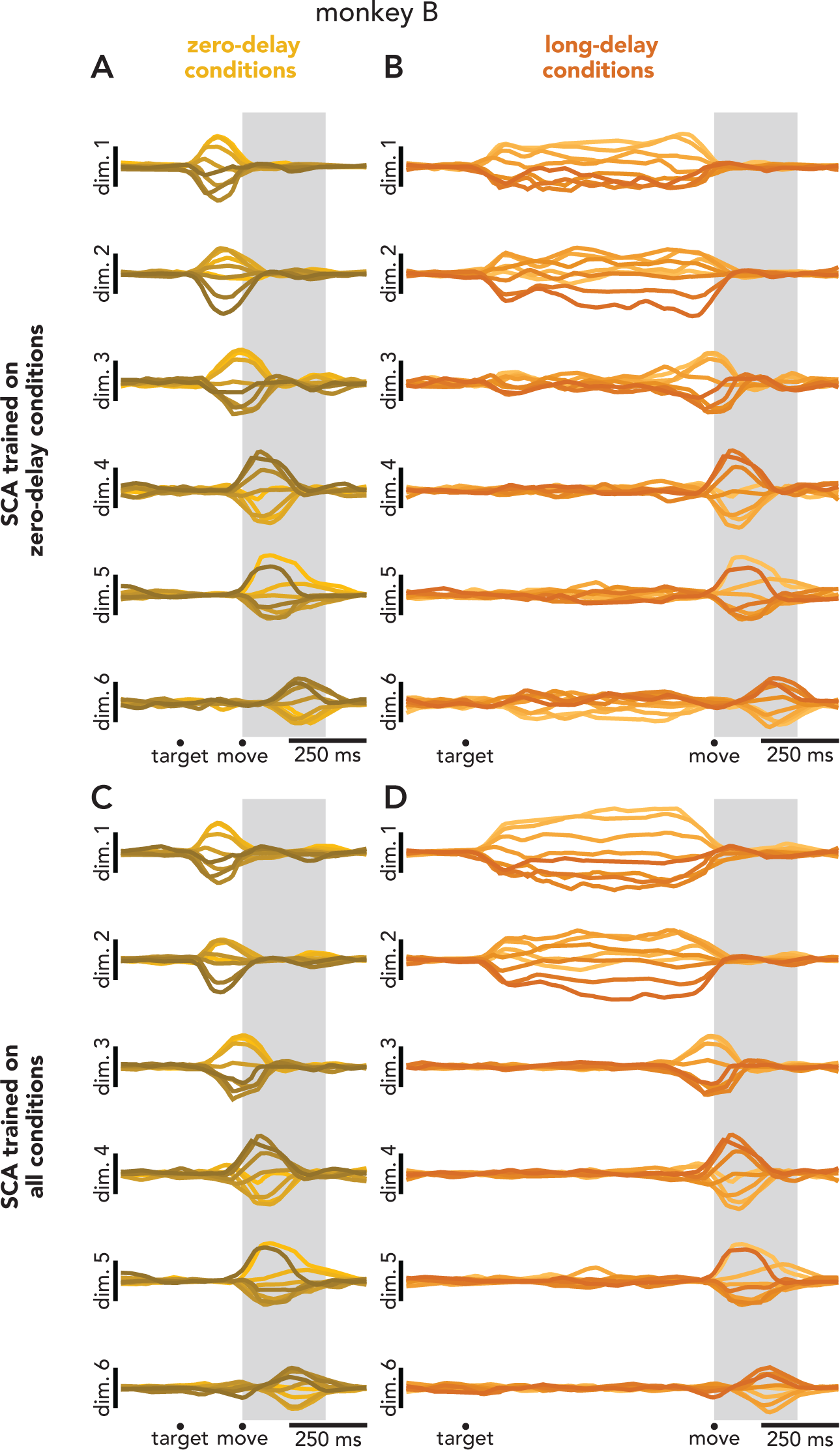
Experimental conditions affect the interpretability of SCA factors. **A.** Projections of eight non-delayed reaching conditions into six SCA dimensions. For these conditions, the target began to fly off of the screen as soon as it was displayed. Therefore, the monkey was required to initiate a reach as soon as the target was presented (i.e., without an instructed delay period, Lara et al. 2018). SCA was trained only on the non-delayed conditions. From these factors, it is unclear whether preparation occurs; activity in dimensions 1 and 2 could be preparatory or early execution-related activity. **B.** Projections of eight delayed conditions into the same six SCA dimensions as in **A**. These conditions were standard, delayed center-out reaching conditions. By projecting the delayed conditions into the same SCA dimensions, it becomes clear that dimensions 1 and 2 are indeed preparatory. **C,D**. Same as **A**,**B**, except SCA was trained on both delayed and non-delayed conditions.

**Figure S12.**
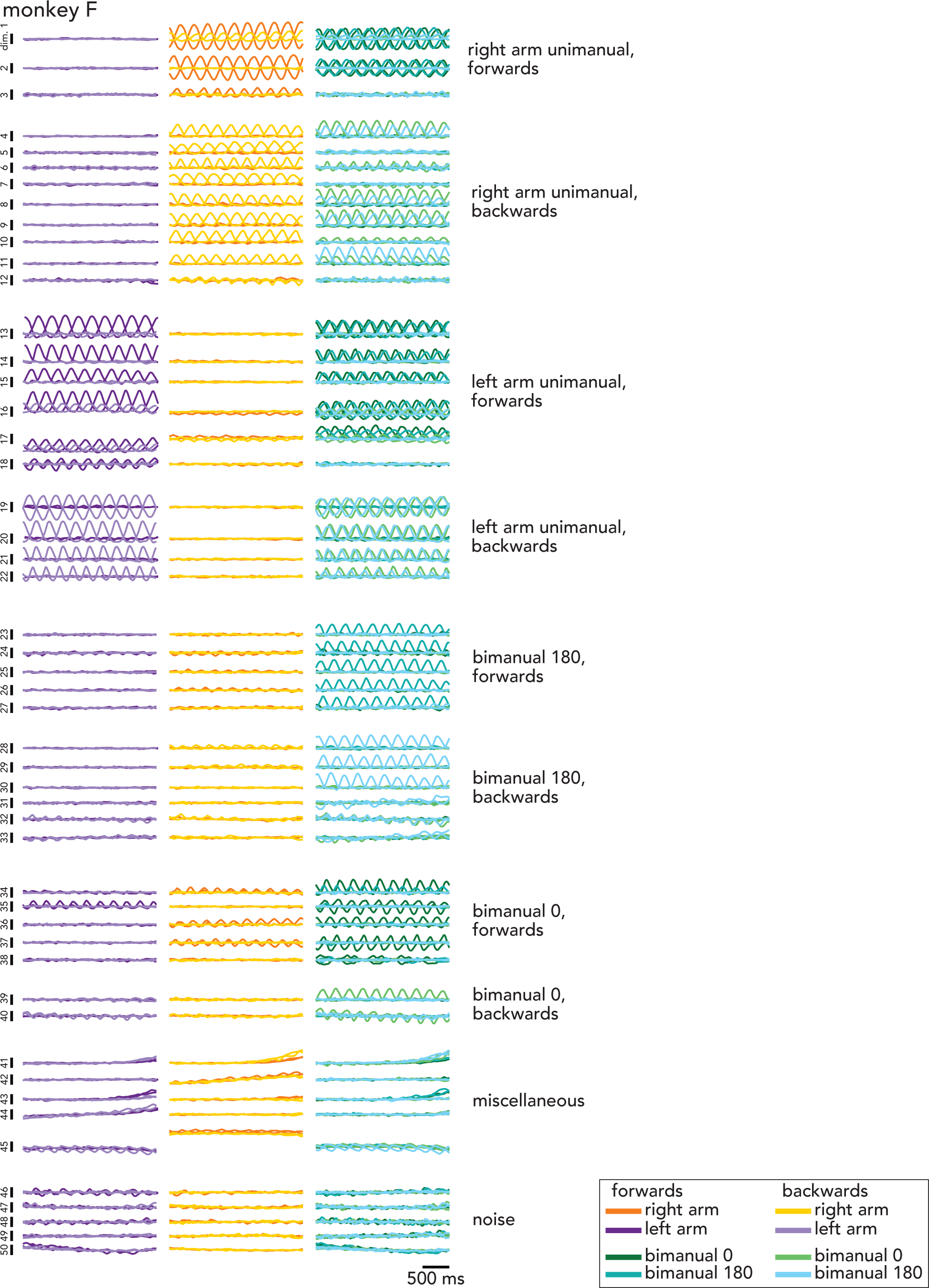
All SCA dimensions for monkey F. Factors were grouped manually (and imperfectly) by the conditions during which they were most active. A factor was placed in a ‘unimanual’ group if the factor was strongly active during a unimanual condition. A factor was placed in a ‘bimanual’ group if it was strongly active during a bimanual condition and weakly active during all unimanual conditions.

**Figure S13.**
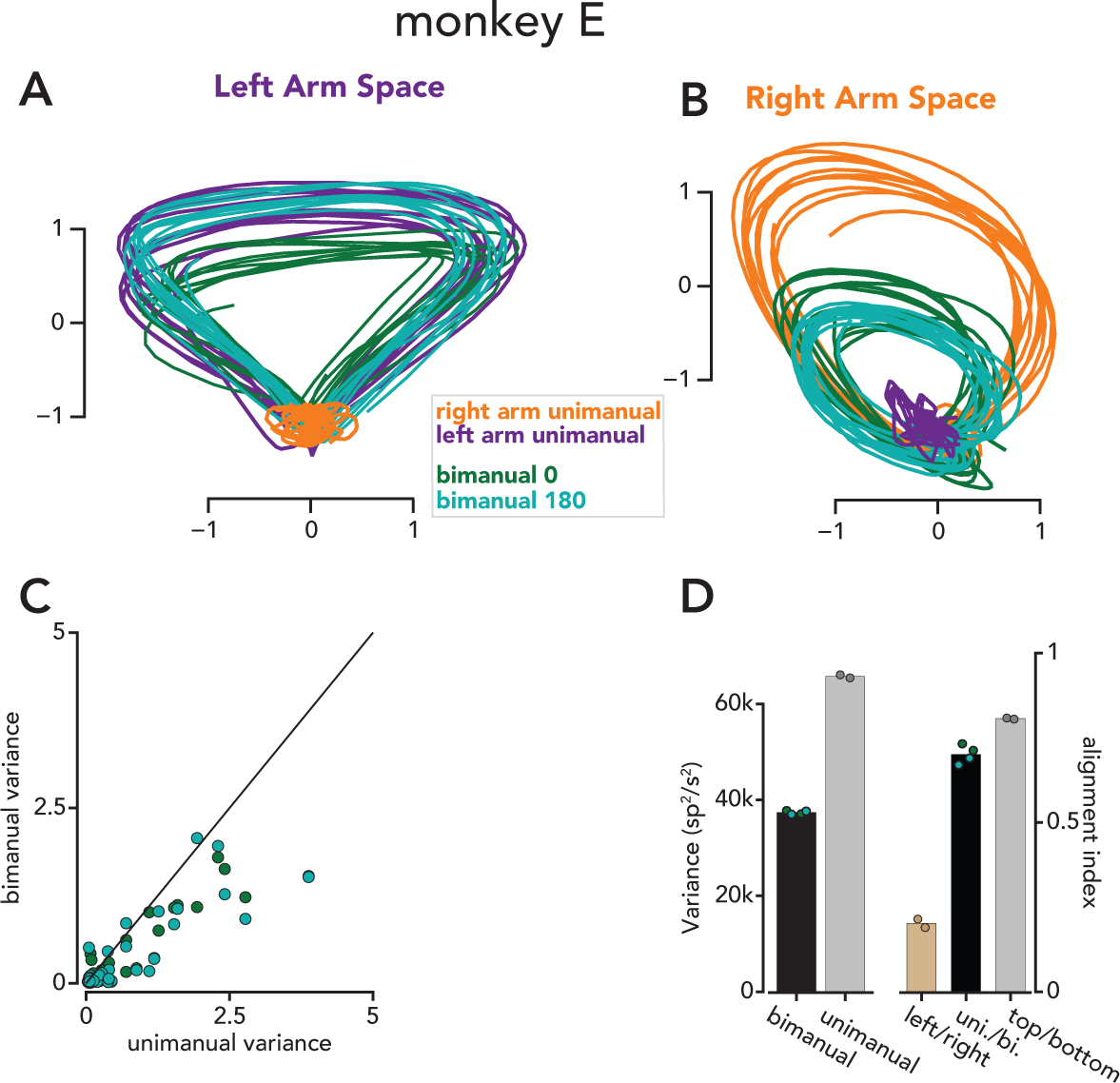
Summary of SCA results from a second bimanual cycling monkey. Same format as Figure 5. **A.** All forward conditions plotted in a space that is spanned by the two SCA dimensions that accounted for the largest fraction of left arm unimanual variance. **B.** Same as **A**, but for SCA dimensions that captured a large fraction of right arm unimanual variance. **C.** Bimanual and unimanual variance in 50 SCA dimensions. **D.** Left: total neural variance during bimanual and unimanual conditions. Right: Alignment indices between left and right unimanual conditions.

**Figure S14.**
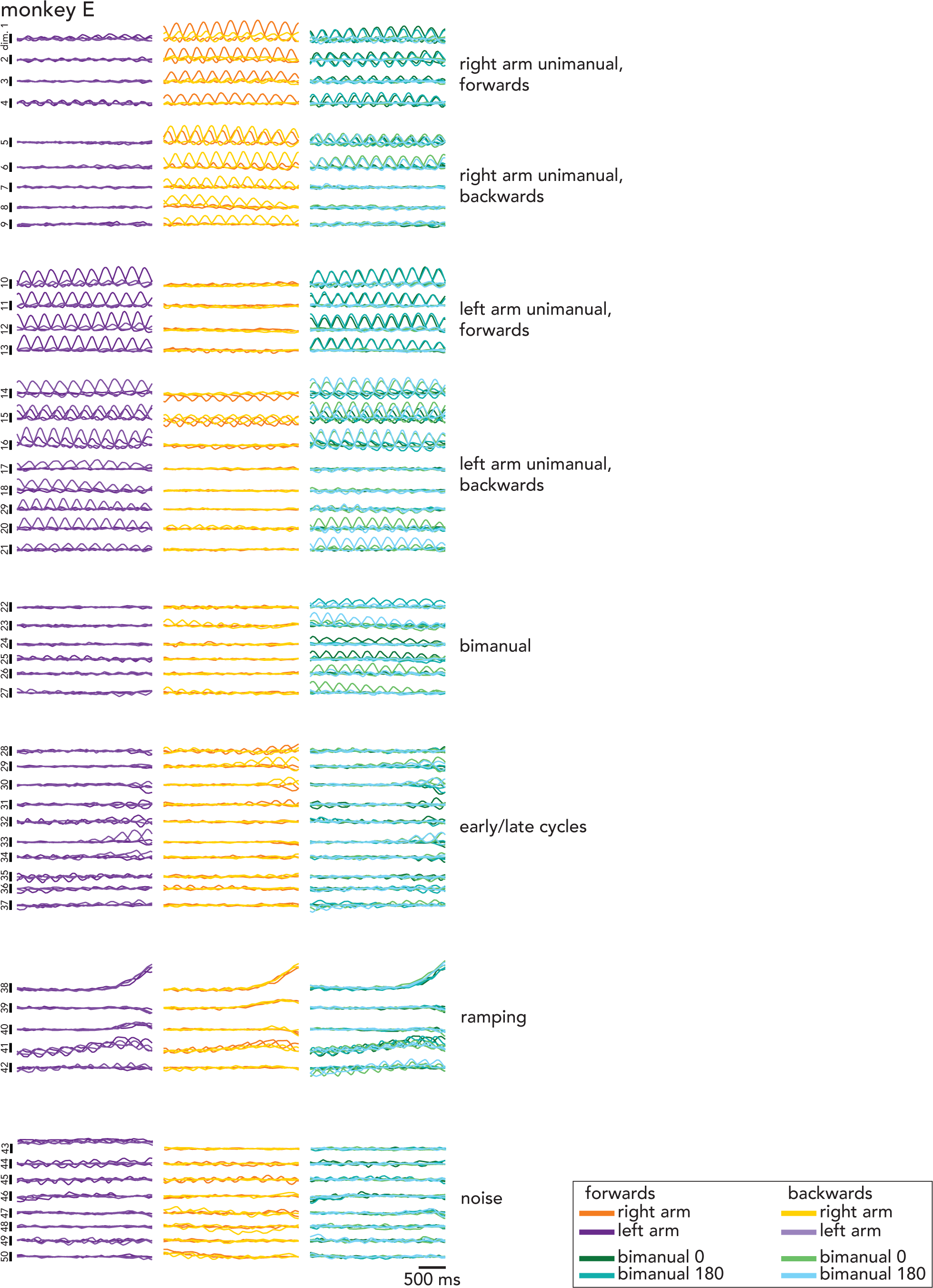
All SCA dimensions for monkey E. Same format as Figure S12. Across the four cycling monkeys, the degree of overlap between the neural dimensions occupied during forward and backward cycling differed fairly substantially; the alignment index between forwards and backwards cycling varied from 0.16 to 0.47 across monkeys. This difference in alignment was mirrored by the degree to which individual SCA factors were active for forward or backward cycling (compare Figures. S7, S10, S12, and S14). Whether these monkey-to-monkey differences reflect meaningful computational differences, with different, perhaps subtle, behavioral consequences, remains to be seen. Alternatively, the differences in overlap between forward and backward cycling could be largely arbitrary; networks may be able to fulfill computational requirements (e.g., generate neural trajectories with low tangling (Russo et al. 2018)) using either partially-aligned or largely-unaligned subspaces.

**Figure S15.**
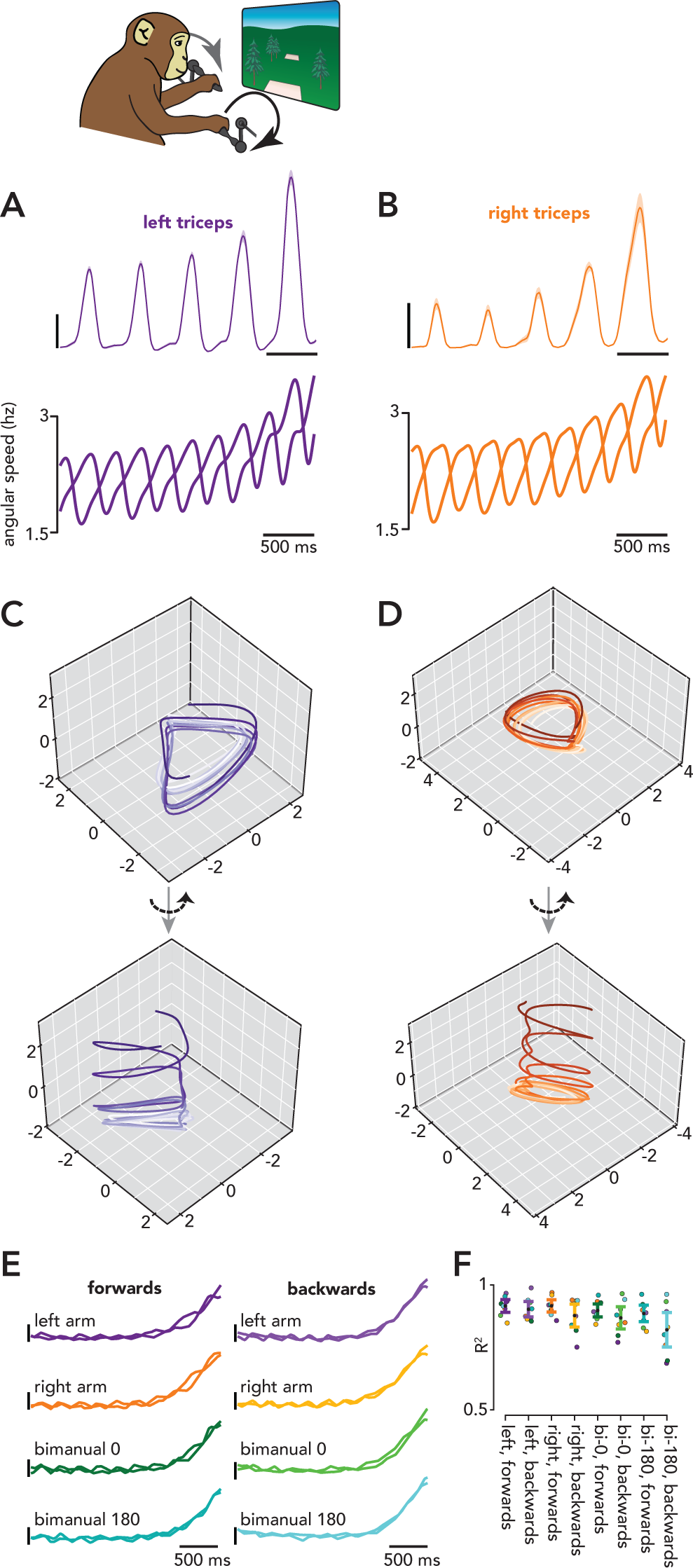
SCA identifies a shared ‘speed’ dimension in motor cortical bimanual cycling data. Prior work has explored the mechanisms underlying cycling at different speeds in both motor cortex and artificial networks (Saxena et al. 2022). Networks trained to empirical muscle activity recorded during cycling exhibit large, oscillatory signals that are correlated with the phase of the cycle; faster cycling speeds are associated with faster rotations in network activity. However, if network activity were to traverse the same region of state space at different speeds, ‘tangling’ (Russo et al. 2018) would be high, and the network would be susceptible to noise (Russo et al. 2018). To maintain low tangling, the networks translate the oscillatory signals associated with cycling at different speeds and/or ‘tilt’ these signals into different dimensions; such structure was also observed in motor cortex activity (Saxena et al. 2022). These prior results make the straightforward prediction that if a monkey were to steadily increase its cycling speed over time (rather than cycle at different speeds on different trials, as was done in Saxena et al.), activity in motor cortex should be helical. During the bimanual cycling task, one monkey (monkey E) exhibited the idiosyncratic behavior of substantially increasing his cycling speed towards the end of a trial. **A.** Top: trial-averaged EMG from the lateral head of the left triceps during a left arm unimanual condition. Bottom: Trial-averaged angular speed of the left arm during two left arm unimanual cycling conditions. **B.** Same as **A**, but for the right arm during right arm unimanual conditions. We performed SCA on all 16 conditions (2 cycling directions x 4 arm conditions x 2 starting pedal positions) and recovered the helical structure hypothesized by Saxena et al. SCA recovered a single dimension in which activity correlated with average cycling speed (rho = 0.92±0.03, mean and standard deviation, across conditions). To illustrate the geometry of these signals, we have plotted activity in two three-dimensional state spaces (**C,D**); two of the dimensions capture the large, oscillatory signals that are present throughout the trial, and the third dimension is the ‘speed dimension’. The projections can be oriented such that the trajectories trace out a repeating, rotational pattern (**C,D**, top). However, rotating the projection reveals that the trajectories actually form a helix, with the first three cycles evolving along the base of the helix and the final two cycles ascending the helix (**C,D**, bottom). Monkey E increased his cycling speed toward the end of all conditions. **E.** Projections from all sixteen conditions in a single SCA dimension that correlates with intra-cycle speed. The two traces in each dimension correspond to top start and bottom start conditions. Given this result, two possibilities exist. Either activity truly does primarily evolve along a single speed dimension, or there are multiple speed dimensions, and SCA has found a rotation of these dimensions in which activity evolves along a stereotyped trajectory. To disambiguate these possibilities, we performed a regression analysis. For each combination of moving arm and cycling direction, we used activity from the top-start condition to learn a set of neural weights that predicted low-pass filtered cycling speed for that condition. We then predicted the (low-pass filtered) cycling speed for all other bottom-start conditions. **F.** Validation of a single ‘speed axis’ across conditions. For each bottom-start condition, regression was used to identify a single neural dimension that correlated with the angular speed during that condition (see *Methods*). Neural activity from all top start conditions was then projected into this dimension, and the R2 between the projection and angular speed of that top start condition was calculated (*colored dots*). *Error bars* correspond to mean and standard deviation across conditions. While the decoder often performed the best for the matched bottom-start condition (compare the colors of the *error bars* and the highest *circle* in each column of **F**), this was a minor improvement over the non-matched conditions, indicating that the neural activity largely translated along a single dimension as the monkey cycled faster, regardless of condition.

**Figure S16.**
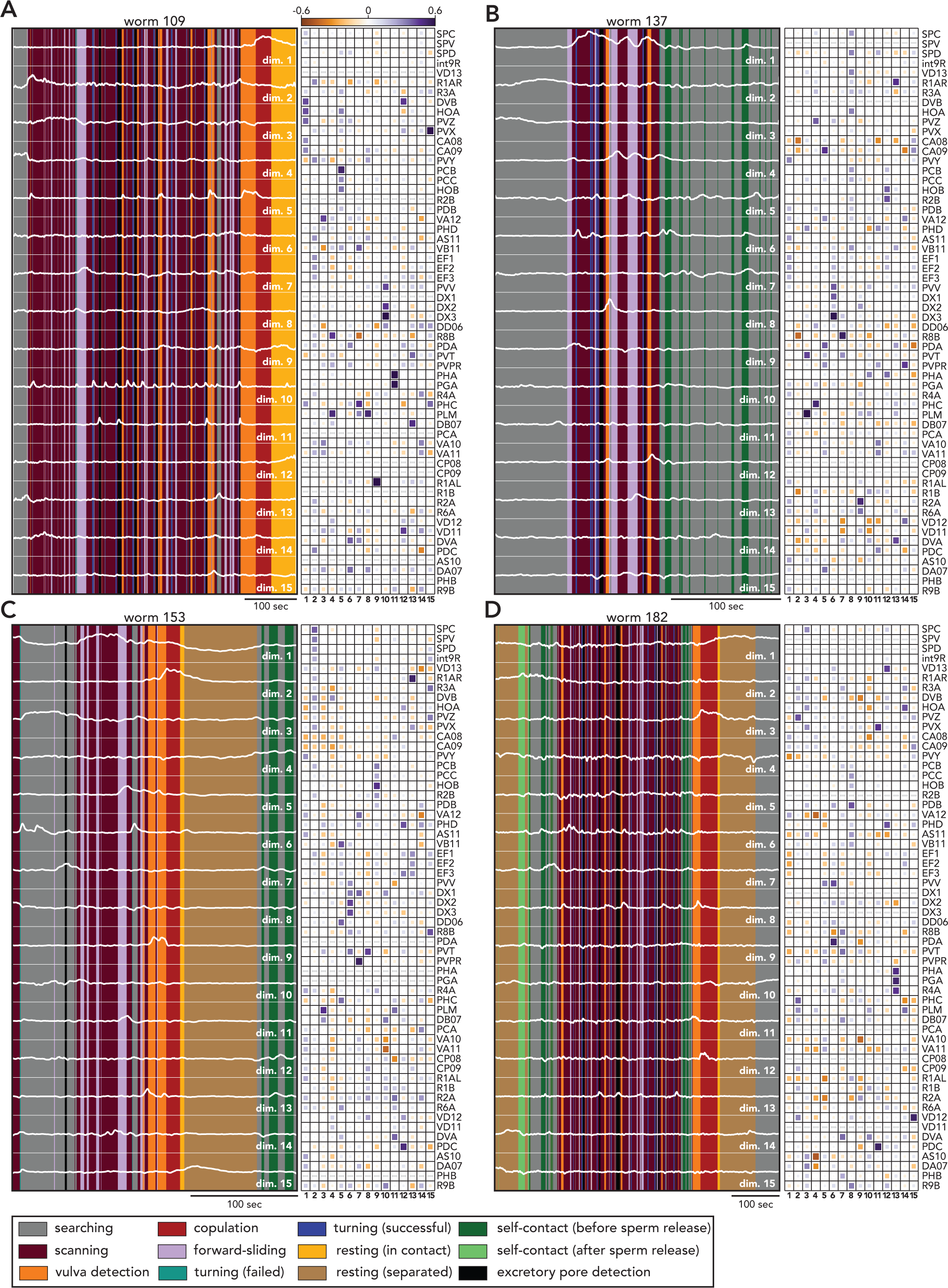
All SCA dimensions for four worms. **A.** Left: Projections in all SCA dimensions (*white traces*), plotted on top of ethogram. Right: SCA loadings. Neurons that were not recorded are indicated with a *gray dash*. **B-D.** Same as **A**, for three additional worms.

**Figure S17.**
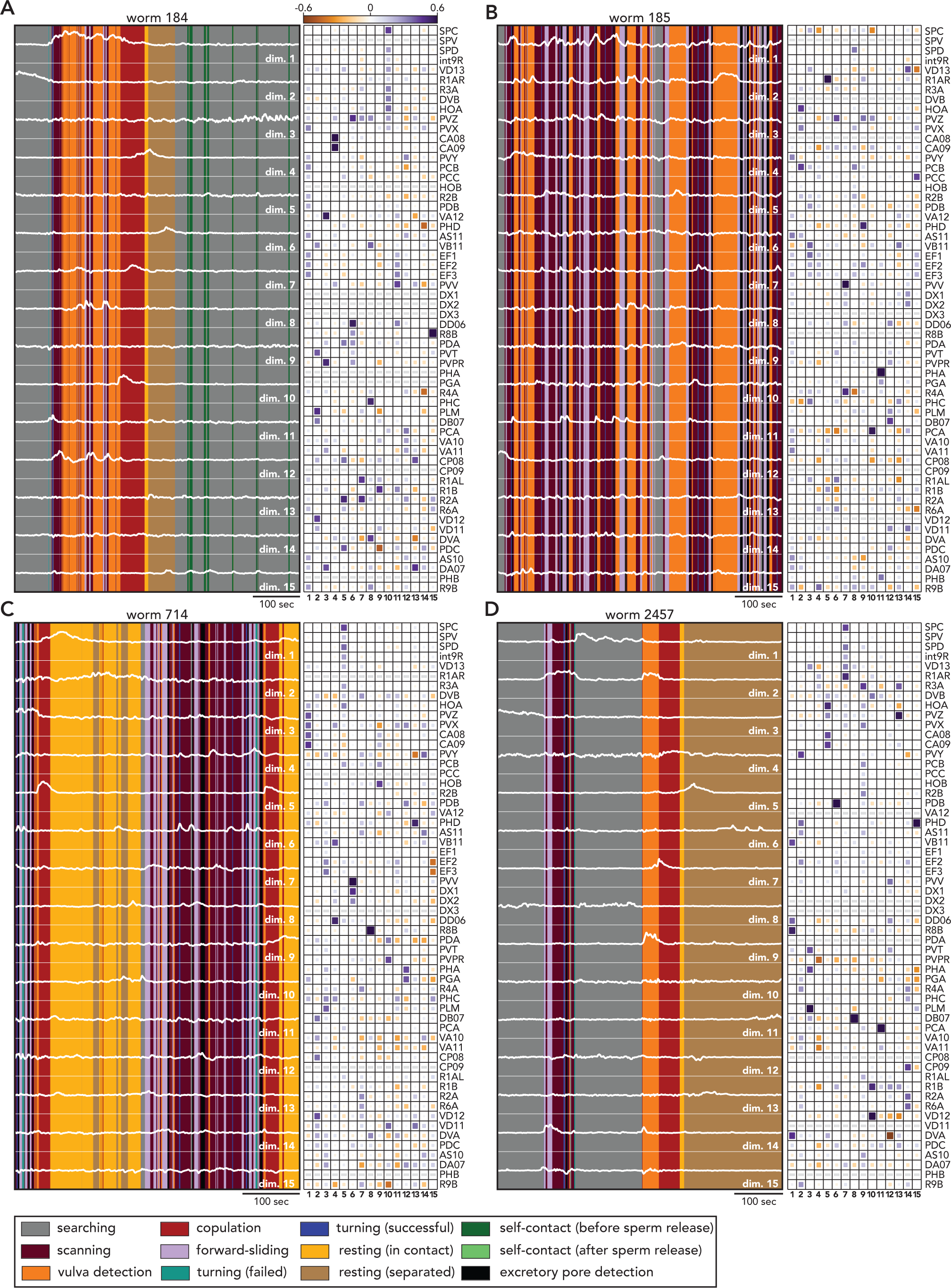
All SCA dimensions for remaining worms. Same format as Figure S16.

**Figure S18.**
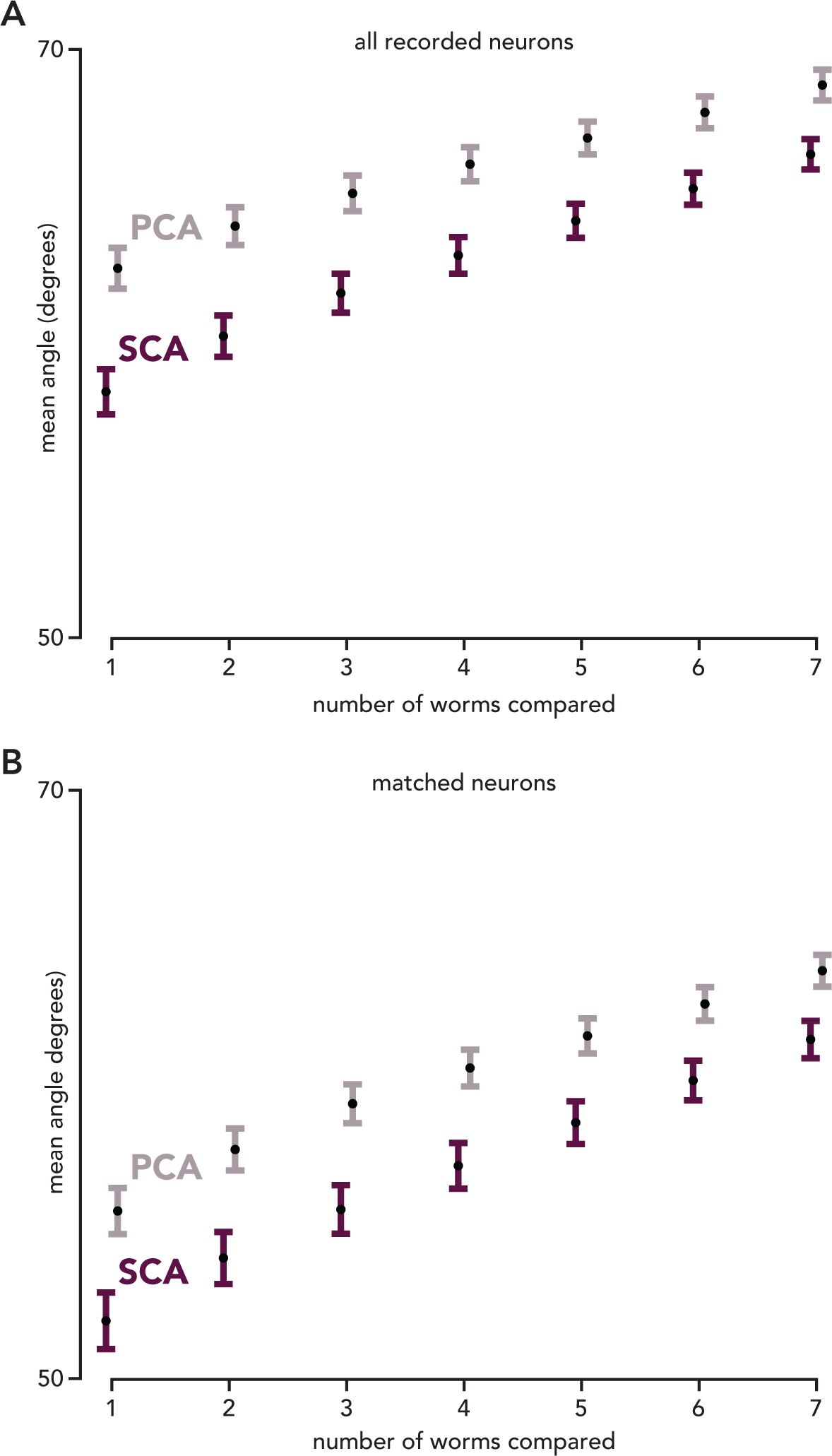
SCA loading vectors are more similar between worms than PCA loading vectors. **A.** For each SCA (or PCA) loading vector for each worm, we identified the seven most similar loading vectors from all other worms (each worm could only contribute a single vector). We then calculated the mean angle between the most similar vectors, the two most similar vectors, etc. Here, the loadings for missing neurons were set to 0. Error bars correspond to standard error, p < 0.001 for all SCA-PCA comparisons, *rank-sum test*. **B.** Same as above, except the analysis is limited to the 33 neurons that were shared between all 8 worms; p < 0.01 for all comparisons, *rank-sum test*.

**Figure S19.**
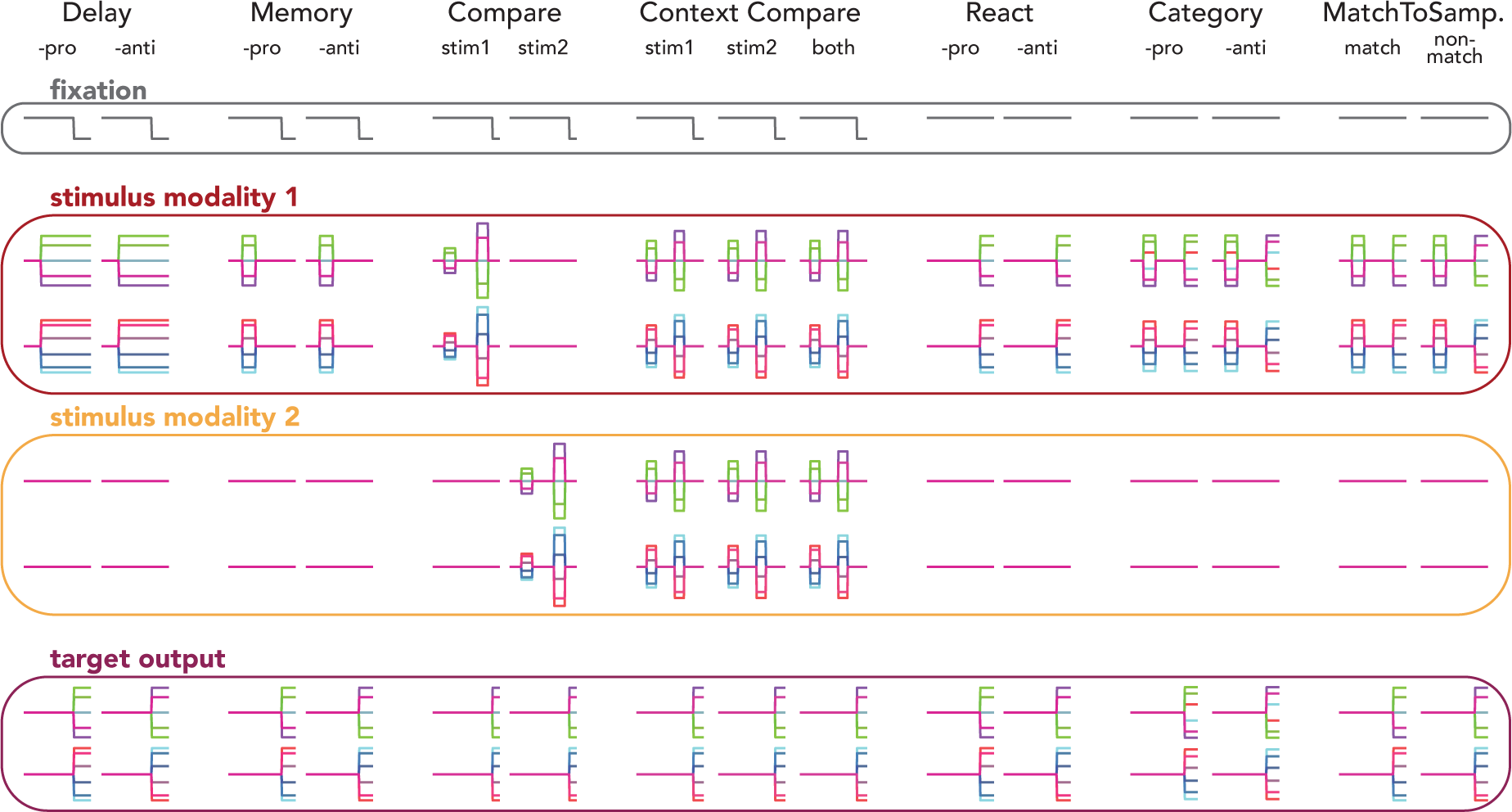
Inputs and target outputs for the multitask network. Across all tasks, the network received three inputs: a 1-dimensional fixation signal (*gray box*), a 4-dimensional stimulus (corresponding to two, 2-dimensional circular stimuli, both scaled by amplitude, *red and yellow boxes*), and a 15-dimensional context cue (not shown). On each trial, the network generated a 2-dimensional output (*maroon box*), corresponding to a single circular variable (the sine and cosine of an angle). The removal of the fixation signal acted as a go cue during Delay-, Memory-, Compare-, and ContextCompare-tasks. During the remaining tasks, output was triggered by the delivery of directional stimuli. The context cue was provided throughout each trial and determined how the network should respond to the directional stimuli. Depending on the task, the network needed to attend to stimulus modality 1 (*red box*), stimulus modality 2 (*yellow box*), or both. For a complete description of each task, see *Methods*.

**Figure S20.**
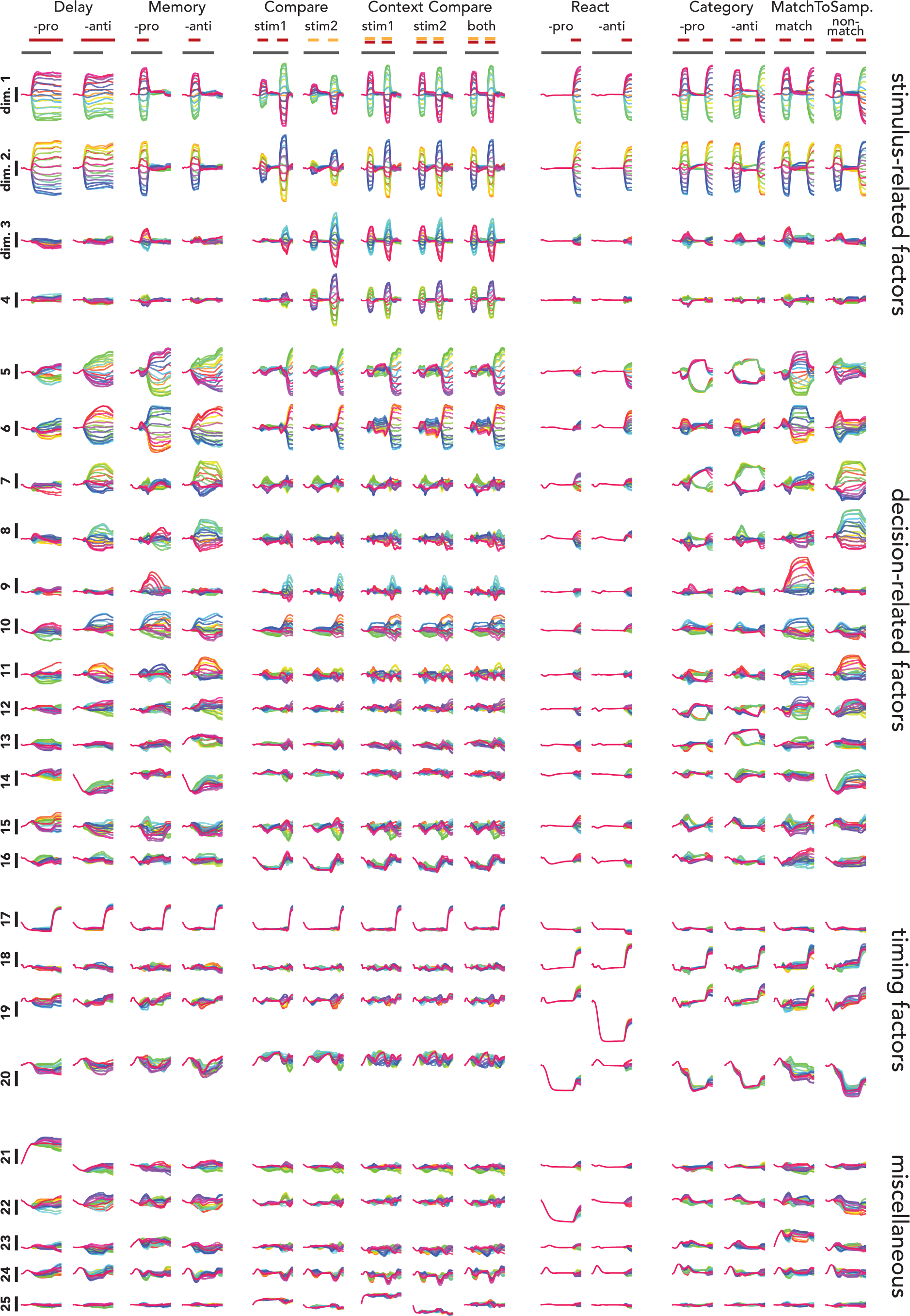
SCA factors identified from a multitask network. SCA was trained on the activity from all fifteen tasks (see *Methods* for a description of each task). Dimensions 1-4 reflected the directional stimuli. Dimensions 1 and 2 were more closely related to input from stimuli 1 (e.g., CompareStim 1), while dimensions 3 and 4 more strongly reflected input from stimuli 2 (e.g., CompareStim 2). In contrast to dimensions 1-4, dimensions 5-16 reflected the network’s ultimate output (i.e., decision). Interestingly, there was only partial overlap between the dimensions that reflected the network’s decision during tasks that involved an external go cue (Delay-, Memory-, Compare-, and Context Compare-) and the dimensions that carried the network’s decision during tasks without an explicit go cue (React-, Category-, and MatchToSample). Consider dimensions 5 and 6. As outlined in Results, these dimensions strongly reflect the network’s decision during the first nine tasks (i.e., the across-condition variance in these dimensions after the go cue is delivered is substantial). During the Category-tasks, however, across-condition variance in these dimensions decreases after the go cue. However, during the Category-tasks, variance increases after the go cue in dimensions 7 and 8. The partial overlap of the ‘decision dimensions’ for different tasks likely relates to the fact that the network seemed to use different sets of factors to trigger output generation, depending on which task was being performed. During the first nine tasks (Delay-, Memory-, Compare-, and ContextCompare-) the fixation cue was removed at the end of each trial, acting as a go cue. During the final six tasks (React-, Category-, and MatchToSamp.-) the fixation cue was never removed; here, the timing of the network’s output was determined by the directional stimuli. SCA recovered factors that reflected two different triggering mechanisms. As discussed in Results, the condition-invariant activity in dimension 17 resembles the ‘trigger signal’ observed in neural data (Kaufman et al. 2016). Note that during the final six tasks, factor 17 was not active. However, a different factor (factor 18) was only active during tasks without a fixation cue. During these tasks, factor 18 also exhibited condition-invariant activity that was time-locked to the production of an output, suggesting that it was acting as a ‘trigger signal’ during tasks without a dedicated external go cue.

**Figure S21.**
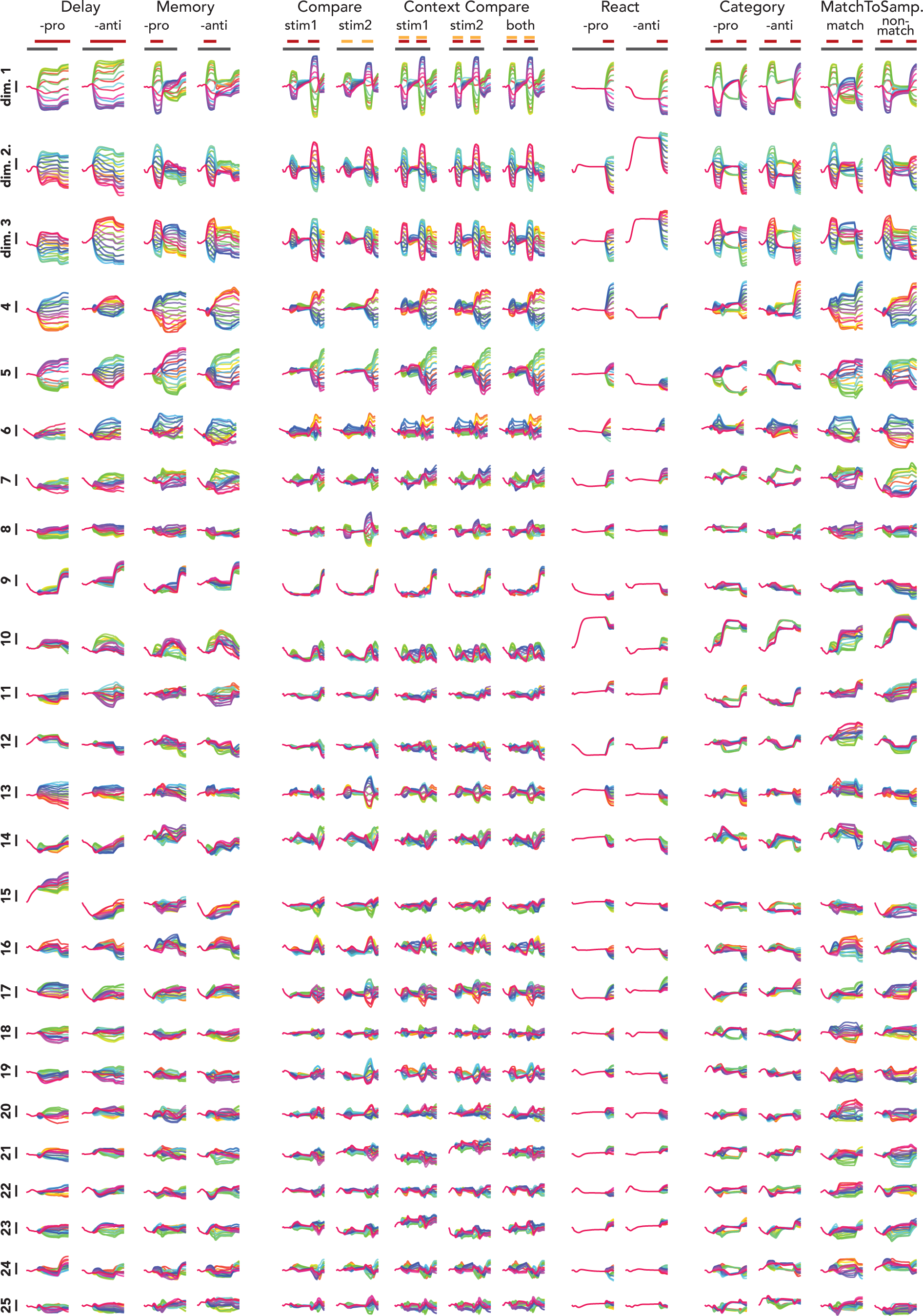
PCA factors identified from a multitask network. Factors identified via PCA do not offer the same clear view of the underlying computations provided by the SCA factors. While some individual dimensions are similar to single SCA dimensions (e.g., dimensions 9 appears to be the ‘trigger signal’ observed in SCA dimension 17, Figure S20), PCA factors more often appear to be ‘mixed’ versions of SCA factors. For example, dimensions 1 and 2 appear to be a mix of stimulus and decision related activity.

